# A novel computational pipeline for *var* gene expression augments the discovery of changes in the *Plasmodium falciparum* transcriptome during transition from *in vivo* to short-term *in vitro* culture

**DOI:** 10.1101/2023.03.21.533599

**Authors:** Clare Andradi-Brown, Jan Stephan Wichers-Misterek, Heidrun von Thien, Yannick D. Höppner, Judith A. M. Scholz, Helle Hansson, Emma Filtenborg Hocke, Tim-Wolf Gilberger, Michael F. Duffy, Thomas Lavstsen, Jake Baum, Thomas D. Otto, Aubrey J. Cunnington, Anna Bachmann

**Affiliations:** Section of Paediatric Infectious Disease, Department of Infectious Disease, Imperial College London, UK; Department of Life Sciences, Imperial College London, South Kensington, London, SW7 2AZ, UK; Centre for Paediatrics and Child Health, Imperial College London, UK; Bernhard Nocht Institute for Tropical Medicine, Bernhard-Nocht-Strasse 74, 20359 Hamburg, Germany; Centre for Structural Systems Biology, Hamburg, Germany, Notkestraße 85, 22607 Hamburg, Germany; Biology Department, University of Hamburg, Hamburg, Germany; Center for Medical Parasitology, Department of Immunology and Microbiology, University of Copenhagen, 2200 Copenhagen, Denmark; Department of Infectious Diseases, Copenhagen University Hospital, 2200 Copenhagen, Denmark; Department of Microbiology and Immunology, University of Melbourne, Melbourne/Parkville VIC 3052, Australia; School of Biomedical Sciences, Faculty of Medicine & Health, UNSW, Kensington, Sydney, 2052, Australia; School of Infection & Immunity, MVLS, University of Glasgow, UK; German Center for Infection Research (DZIF), partner site Hamburg-Borstel-Lübeck-Riems, Germany

## Abstract

The pathogenesis of severe *Plasmodium falciparum* malaria involves cytoadhesive microvascular sequestration of infected erythrocytes, mediated by *P. falciparum* erythrocyte membrane protein 1 (PfEMP1). PfEMP1 variants are encoded by the highly polymorphic family of *var* genes, the sequences of which are largely unknown in clinical samples. Previously, we published new approaches for *var* gene profiling and classification of predicted binding phenotypes in clinical *P. falciparum* isolates (Wichers *et al*., 2021), which represented a major technical advance. Building on this, we report here a novel method for *var* gene assembly and multidimensional quantification from RNA-sequencing that outperforms the earlier approach of Wichers *et al*., 2021 on both laboratory and clinical isolates across a combination of metrics. Importantly, the tool can interrogate the *var* transcriptome in context with the rest of the transcriptome and can be applied to enhance our understanding of the role of *var* genes in malaria pathogenesis. We applied this new method to investigate changes in *var* gene expression through early transition of parasite isolates to *in vitro* culture, using paired sets of *ex vivo* samples from our previous study, cultured for up to three generations. In parallel, changes in non-polymorphic core gene expression were investigated. Modest but unpredictable *var* gene switching and convergence towards *var2csa* were observed in culture, along with differential expression of 19% of the core transcriptome between paired *ex vivo* and generation 1 samples. Our results cast doubt on the validity of the common practice of using short-term cultured parasites to make inferences about *in vivo* phenotype and behaviour.

## Introduction

Malaria is a parasitic life-threatening disease caused by species of the *Plasmodium* genus. In 2021, there were an estimated 619,000 deaths due to malaria, with children under 5 accounting for 77% of these (WHO, 2022). *Plasmodium falciparum* causes the greatest disease burden and most severe outcomes, but our efforts to combat the disease are challenged by its complex life cycle and its sophisticated immune evasion strategies. *P. falciparum* has several highly polymorphic variant surface antigens (VSA) encoded by multi-gene families, with the best studied high molecular weight *Plasmodium falciparum* erythrocyte membrane protein 1 (PfEMP1) family of proteins known to play a major role in the pathogenesis of malaria (Leech *et al*., 1984, Wahlgren *et al*., 2017). About 60 polymorphic *var* genes per parasite genome encode different PfEMP1 variants, which are exported to the surface of parasite-infected erythrocytes, where they mediate cytoadherence to host endothelial cells (Leech *et al*., 1984, Su *et al*., 1995, Smith *et al*., 1995, Baruch *et al*., 1995, Rask *et al*., 2010). *Var* genes are expressed in a mutually exclusive pattern, resulting in each parasite expressing only one *var* gene, and therefore one PfEMP1 protein, at a time (Scherf *et al*., 1998). Due to the exposure of PfEMP1 proteins to the host immune system, switching expression between the approximately 60 *var* genes in the genome is an effective immune evasion strategy, which can result in selection and dominance of parasites expressing particular *var* genes within each host (Smith *et al*., 1995).

Despite their sequence polymorphism, *var* genes could be classified into four categories (A, B, C, and E) according to their chromosomal location, transcriptional direction, type of 5’-upstream sequence (UPSA–E), and encoded protein domains with associated binding phenotype (Figure 1) (Lavstsen *et al*., 2003, Kraemer & Smith, 2003, Kyes *et al*., 2007, Rask *et al*., 2010). PfEMP1 proteins have up to 10 extracellular domains, with the N-terminal domains forming a semi-conserved head structure complex typically containing the N-terminal segment (NTS), a Duffy binding-like domain of class α (DBLα) coupled to a cysteine-rich interdomain region (CIDR). C-terminally to this head structure, PfEMP1 proteins exhibit a varying but semi-ordered composition of additional DBL and CIDR domains of different subtypes (Figure 1c). The PfEMP1 family divides into three main groups based on the receptor specificity of the N-terminal CIDR domain: (i) PfEMP1 proteins with CIDRα1 domains bind endothelial protein C receptor (EPCR), while (ii) PfEMP1 proteins with CIDRα2–6 domains bind CD36 and (iii) the atypical VAR2CSA PfEMP1 proteins bind placental chondroitin sulphate A (CSA) (Salanti *et al*., 2004). In addition to these, a subset of PfEMP1 proteins have N-terminal CIDRβ/γ/δ domains of unknown function. This functional diversification correlates with the genetic organization of the *var* genes. Thus, UPSA *var* genes encode PfEMP1 proteins with domain sub-variants NTSA-DBLα1-CIDRα1/β/γ/δ, whereas UPSB and UPSC *var* genes encode PfEMP1 proteins with NTSB-DBLα0-CIDRα2–6. One exception to this rule is the B/A chimeric *var* genes, which encode NTSB-DBLα2-CIDRα1 domains. The different receptor binding specificities are associated with different clinical outcomes of infection. Pregnancy-associated malaria is linked to parasites expressing VAR2CSA, whereas parasites expressing EPCR-binding PfEMP1 are linked to severe malaria and parasites expressing CD36-binding PfEMP1 are linked to uncomplicated malaria (Turner *et al*., 2013, Lavstsen *et al*., 2012, Avril *et al*., 2012, Claessens *et al*., 2012, Tonkin-Hill *et al*., 2018, Wichers *et al*., 2021). The clinical relevance of PfEMP1 proteins with unknown binding phenotypes of the N-terminal head structure and C-terminal PfEMP1 domains is largely unknown, albeit specific interactions with endothelial receptors and plasma proteins have been described (Tuikue Ndam *et al*., 2017, Quintana *et al*., 2019, Stevenson *et al*., 2015). Each parasite genome carries a similar repertoire of *var* genes, which in addition to the described variants include a highly conserved *var1* variant of either type 3D7 or IT, which in most genomes occurs with a truncated or absent exon 2. Also, most genomes carry the unusually small and highly conserved *var3* genes, of unknown function (Figure 1c) (Otto *et al*., 2019).

**Figure 1:**
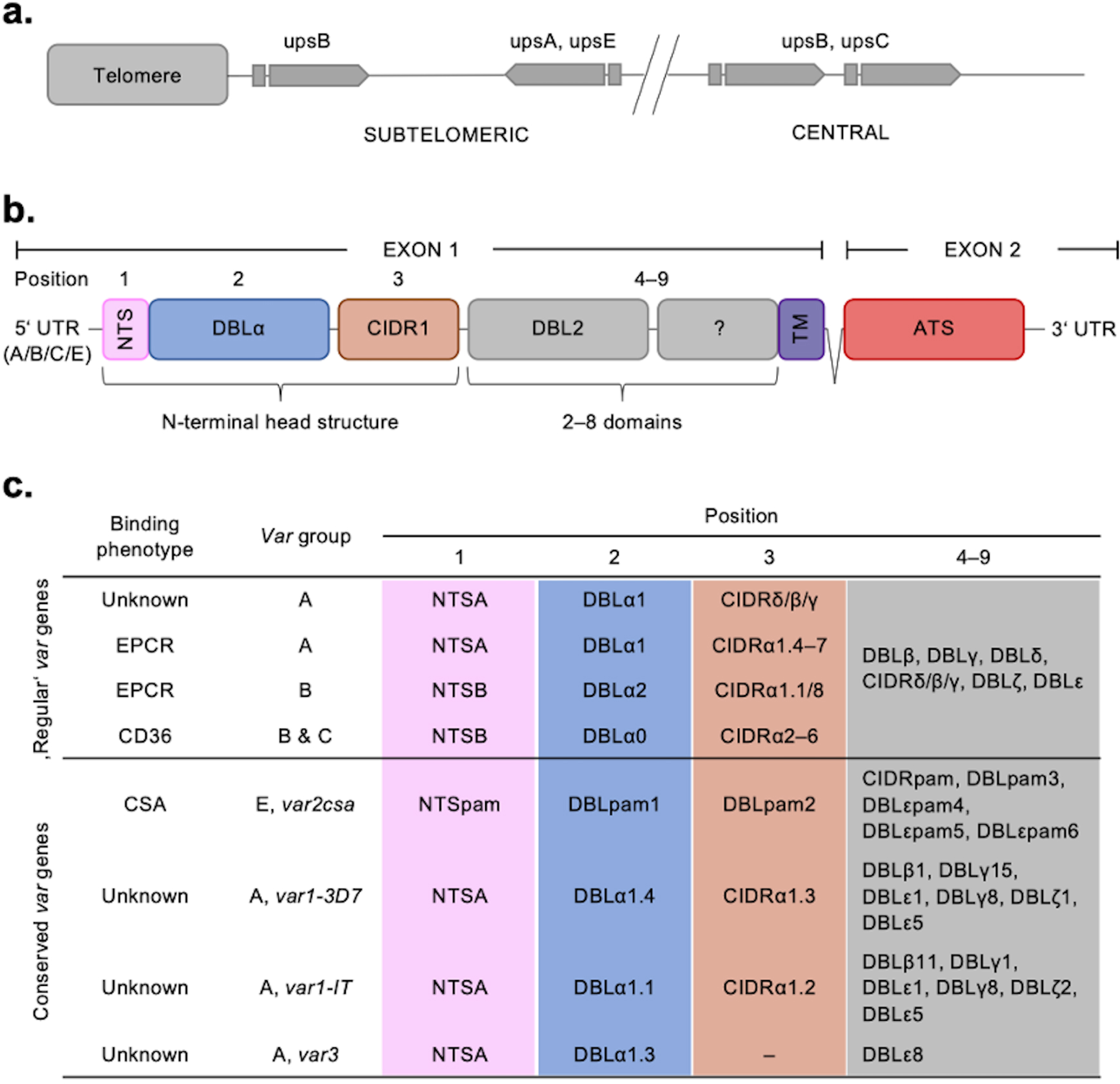
Summary of the *var* chromosomal location, *var* gene, PfEMP1 protein structure, and PfEMP1 binding phenotypes. **a)** Chromosomal position and transcriptional direction (indicated by arrows) of the different *var* gene groups, designated by the respective type of upstream sequence (Kraemer and Smith et al., 2003, Lavstsen et al., 2003). **b)** Structure of the *var* gene which encodes the PfEMP1 protein. The *var* gene is composed of two exons, the first, around 3–9.4 kb, encodes the highly variable extracellular region and the transmembrane region (TM) of PfEMP1. Exon 2 is shorter with about 1.2 kb and encodes a semi-conserved intracellular region (acidic terminal segment, ATS). The PfEMP1 protein is composed of an N-terminal segment (NTS), followed by a variable number of Duffy binding-like (DBL) domains and cysteine-rich interdomain regions (CIDR) (Rask et al., 2010). **c)** Summary of PfEMP1 proteins encoded in the parasite genome, their composition of domain subtypes and associated N-terminal binding phenotype. Group A and some B proteins have an EPCR-binding phenotype; the vast majority of group B and C PfEMP1 proteins bind to CD36. Group A proteins also include those that bind a yet unknown receptor, as well as VAR1 and VAR3 variants with unknown function and binding phenotype. VAR2CSA (group E) binds placental CSA.

Comprehensive characterisation and quantification of *var* gene expression in field samples have been complicated by biological and technical challenges. The extreme polymorphism of *var* genes precludes a reference *var* sequence. *Var* genes can be lowly expressed or not expressed at all, contain repetitive domains and can have large duplications (Otto *et al*., 2019). Consequently, most studies relating *var* gene expression to severe malaria have relied on primers with restricted coverage of the *var* family, use of laboratory-adapted parasite strains or have predicted the downstream sequence from DBLα domains (Sahu *et al*., 2021, Storm *et al*., 2019, Shabani *et al*., 2017, Mkumbaye *et al*., 2017, Kessler *et al*., 2017, Bernabeu *et al*., 2016, Jespersen *et al*., 2016, Lavstsen *et al*., 2012). This has resulted in incomplete *var* gene expression quantification and the inability to elucidate specific or detect atypical *var* sequences. RNA-sequencing has the potential to overcome these limitations and provide a better link between *var* expression and PfEMP1 phenotype in *in vitro* assays, co-expression with other genes or gene families and epigenetics. While approaches for *var* assembly and quantification based on RNA-sequencing have recently been proposed (Wichers *et al*., 2021; Stucke et al., 2021; Andrade et al., 2020; Tonkin-Hill *et al*., 2018, Duffy et al., 2016), these still produce inadequate assembly of the biologically important N-terminal domain region, have a relatively high number of misassemblies and do not provide an adequate solution for handling the conserved *var* variants (Table 1).

**Table 1:**
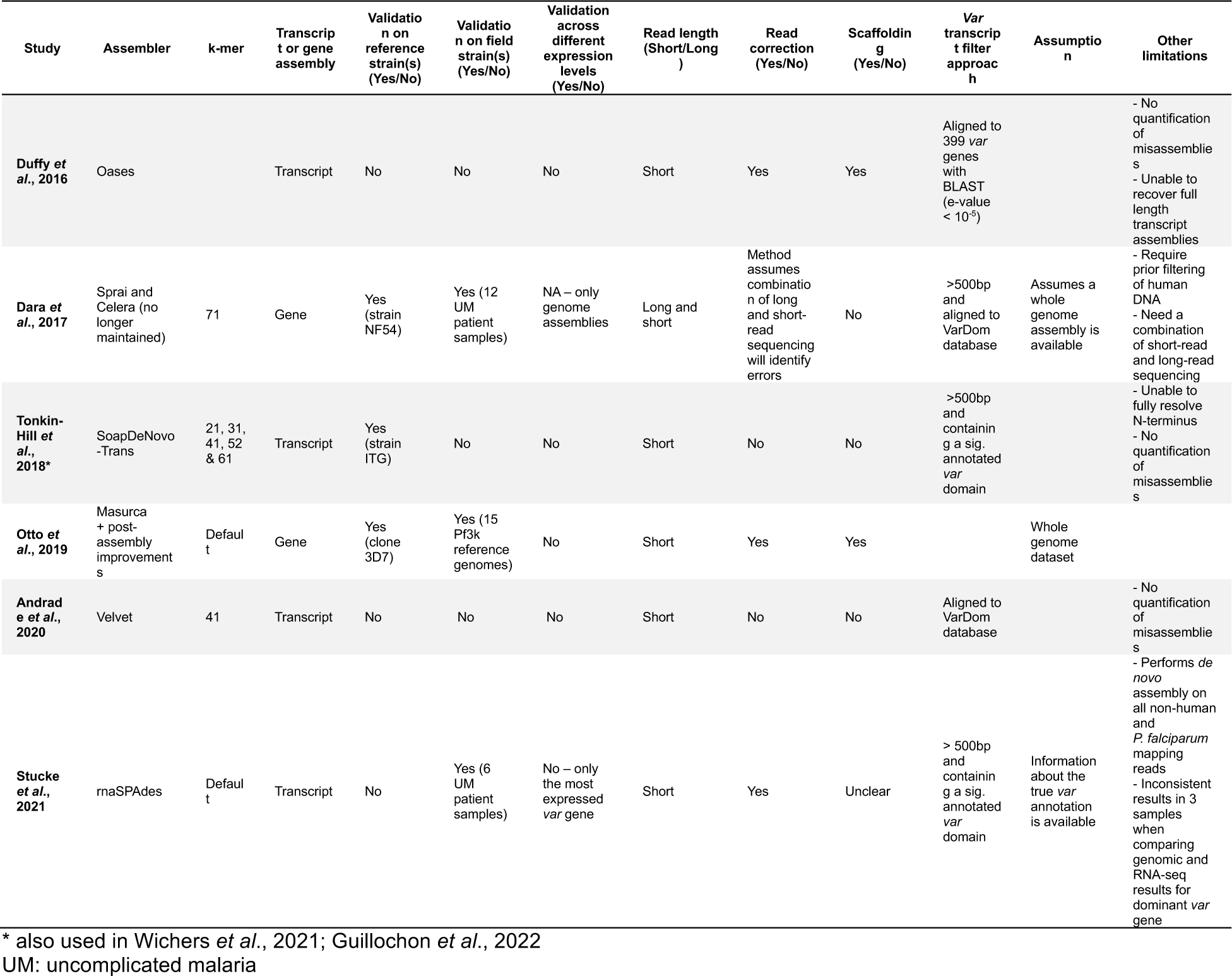
Comparison of previous *var* assembly approaches based on DNA- and RNA-sequencing.

*Plasmodium* parasites from human blood samples are often adapted to or expanded through *in vitro* culture to provide sufficient parasites for subsequent investigation of parasite biology and phenotype (Brown & Guler, 2020). This is also the case for several studies assessing the PfEMP1 phenotype of parasites isolated from malaria-infected donors (Pickford *et al*., 2021, Joste *et al*., 2020, Storm *et al*., 2019, Tuikue Ndam *et al*., 2017, Bruske *et al*., 2016, Claessens *et al*., 2012, Lavstsen *et al*., 2005, Jensen *et al*., 2004, Kirchgatter & Portillo Hdel, 2002, Dimonte *et al*., 2016, Hoo *et al*., 2019). However, *in vitro* conditions are considerably different to those found *in vivo,* for example in terms of different nutrient availability and lack of a host immune response (Brown & Guler, 2020). Previous studies found inconsistent results in terms of whether *var* gene expression is impacted by culture and, if so, which *var* groups were the most affected (Zhang *et al*., 2011, Peters *et al*., 2007). Similar challenges apply to the understanding of changes in *P. falciparum* non-polymorphic core genes in culture, with the focus previously being on long-term laboratory adapted parasites (Claessens *et al*., 2017, Mackinnon *et al*., 2009). Consequently, direct interpretation of a short-term cultured parasite’s transcriptome remains a challenge. It is fundamental to understand *var* genes in context with the parasite’s core transcriptome. This could provide insights into *var* gene regulation and phenomena such as the proposed lower level of *var* gene expression in asymptomatic individuals (Almelli *et al*., 2014, Andrade *et al*., 2020).

Here we present an improved method for assembly, characterization, and quantification of *var* gene expression from RNA-sequencing data. This new approach overcomes previous limitations and outperforms current methods, enabling a much greater understanding of the *var* transcriptome. We demonstrate the power of this new approach by evaluating changes in *var* gene expression of paired samples from clinical isolates *of P. falciparum* during their early transition to *in vitro* culture, across several generations. The use of paired samples, which are genetically identical and hence have the same *var* gene repertoire, allows validation of assembled transcripts and direct comparisons of expression. We complement this with a comparison of changes which occur in the non-polymorphic core transcriptome over the same transition into culture. We find a background of modest changes in *var* gene expression with unpredictable patterns of *var* gene switching, favouring an apparent convergence towards *var2csa* expression. More extensive changes were observed in the core transcriptome during the first cycle of culture, suggestive of a parasite stress response.

## Results

To extend our ability to characterise *var* gene expression profiles and changes over time in clinical *P. falciparum* isolates, we set out to improve current assembly methods. Previous methods for assembling *var* transcripts have focussed on assembling whole transcripts (Tonkin-Hill *et al*., 2018, Wichers *et al*., 2021, Guillochon *et al*., 2022, Andrade *et al*., 2020). However, due to the diversity within PfEMP1 domains, their associations with disease severity and the fact different domain types are not inherited together, a method focussing on domain assembly first was developed. In addition, a novel whole transcript approach, using a different *de novo* assembler, was developed and their performance compared to the method of Wichers *et al*. (hereafter termed “original approach”, Figure 2) (Wichers *et al*., 2021). The new approaches made use of the MalariaGEN *P. falciparum* dataset, which led to the identification of additional multi-mapping non-core reads (a median of 3,955 reads per sample) prior to *var* transcript assembly (MalariaGen *et al*., 2021). We incorporated read error correction and improved large scaffold construction with fewer misassemblies (see Methods). We then applied this pipeline to paired *ex vivo* and short-term *in vitro* cultured parasites to enhance our understanding of the impact of short-term culturing on the *var* transcriptome (Figure 3). The *var* transcriptome was assessed at several complementary levels: first, changes in the dominantly expressed *var* gene and the homogeneity of the *var* expression profile in paired samples were investigated; second, changes in *var* domain expression through culture were assessed; and third, *var* group and global *var* gene expression changes were evaluated. All these analyses on *var* expression were accompanied by analysis of the core transcriptome at the transition to short-term culture.

**Figure 2:**
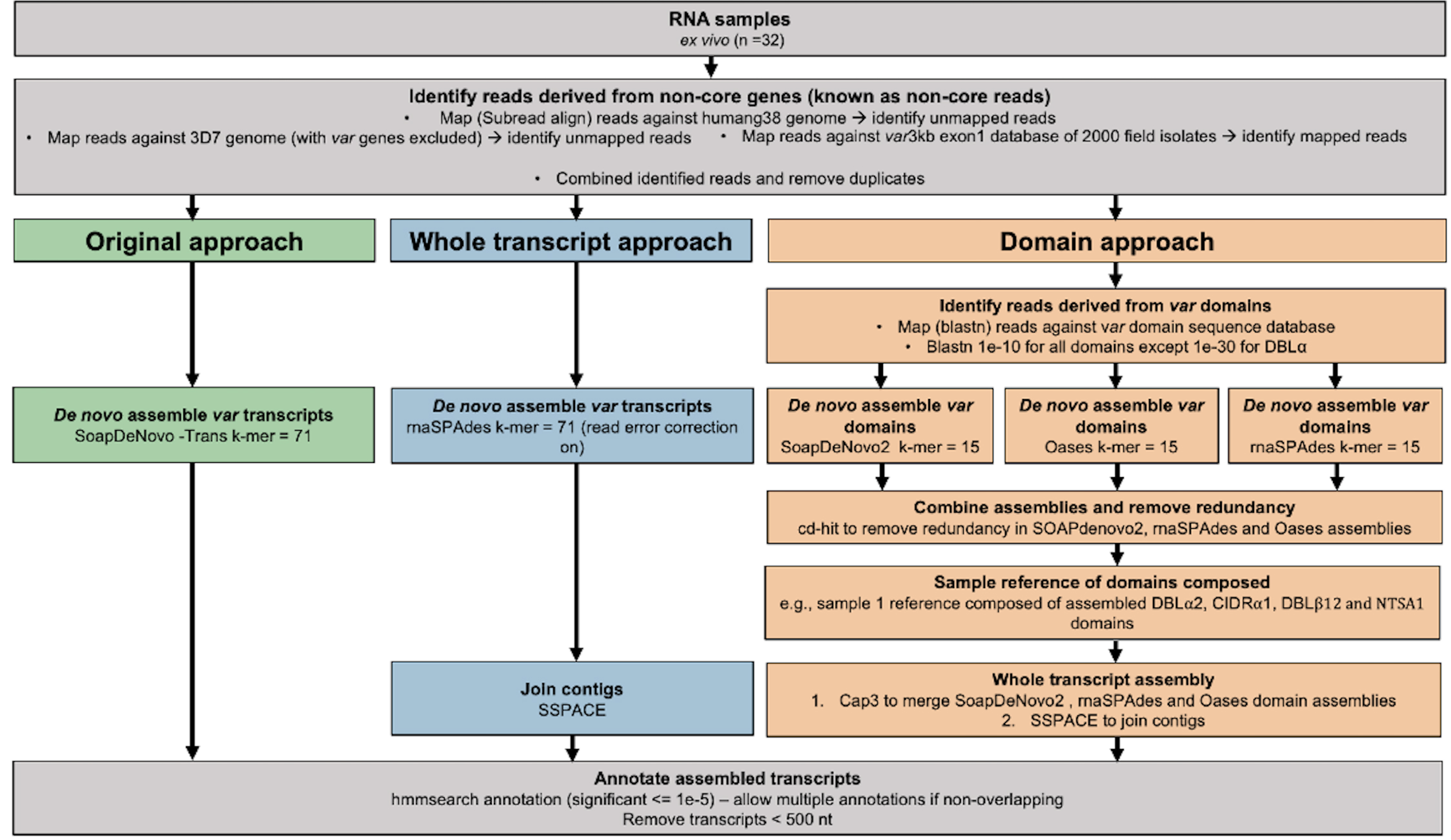
Overview of novel computational pipelines for assembling *var* transcripts. The original approach (green) used SoapDeNovo-Trans (k=71) to perform whole *var* transcript assembly. The whole transcript approach (blue) focused on assembling whole *var* transcripts from the non-core reads using rnaSPAdes (k = 71). Contigs were then joined into longer transcripts using SSPACE. The domain approach (orange) assembled *var* domains first and then joined the domains into whole transcripts. Domains were assembled separately using three different *de novo* assemblers (SoapDeNovo2, Oases and rnaSPAdes). Next, a reference of assembled domains was composed and cd-hit (at sequence identity = 99%) was used to remove redundant sequences. Cap3 was used to merge and extend domain assemblies. Finally, SSPACE was used to join domains together. HMM models built on the Rask *et al*., 2010 dataset were used to annotate the assembled transcripts (Rask *et al*., 2010). The most significant alignment was taken as the best annotation for each region of the assembled transcript (significance <= 1e-5) identified using cath-resolve-hits0. Transcripts < 500nt were removed. A *var* transcript was selected if it contained at least one significantly annotated domain (in exon 1). *Var* transcripts that encoded only the more conserved exon 2 (ATS domain) were discarded. The three pipelines were run on the 32 malaria patient *ex vivo* samples from Wichers *et al*., 2021 (Wichers *et al*., 2021).

**Figure 3:**
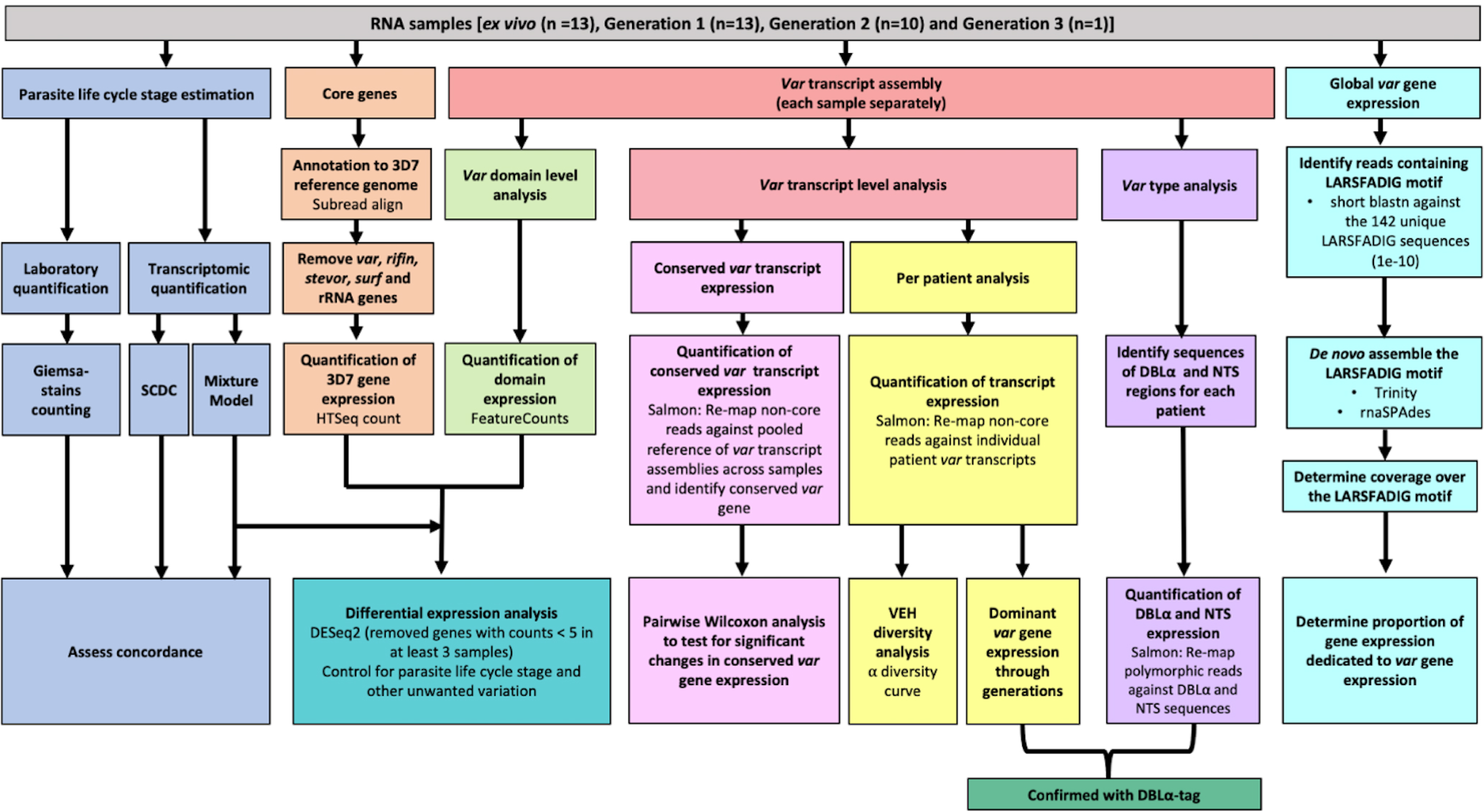
Summary of analyses of *var* and core gene transcriptome changes in paired *ex vivo* and short-term *in vitro* cultured parasites. From a total of 13 parasite isolates, the *ex vivo*samples (Wichers *et al*., 2021) and the corresponding *in vitro*-cultured parasites of the first (n=13), second (n=10) and third (n=1) replication cycle were analysed by RNA sequencing. The expression of non-polymorphic core genes and polymorphic *var* genes was determined in different analysis streams: (1) Non-polymorphic core gene reads were mapped to the 3D7 reference genome, expression was quantified using HTSeq and differential expression analysis performed (orange); (2) Non-core reads were identified, whole transcripts were assembled with rnaSPAdes, expression of both *var* transcripts (red) and domains (light green) was quantified, and *var* domain differential expression analysis was performed. “Per patient analysis” (yellow) represents combining all assembled *var* transcripts for samples originating from the same *ex vivo*sample only. For each conserved *var* gene (*var1-3D7, var1-IT, var2csa* and *var3*) all significantly assembled conserved *var* transcripts were identified and put into a combined reference (pink). The normalised counts for each conserved gene were summed. Non-core reads were mapped to this and DESeq2 normalisation performed. *Var* type (group A vs group B and C) expression (purple) was quantified using the DBLα and NTS assembled sequences and differences across generations were assessed. Total *var* gene expression (turquoise) was quantified by assembling and quantifying the coverage over the highly conserved LARSFADIG motif, with the performance of assembly using Trinity and rnaSPAdes assessed. DBLα-tag data was used to confirm the results of the dominant *var* gene expression analysis and the *var* type analysis (dark green). *Var* expression homogeneity (VEH) was analysed at the patient level (α diversity curves). All differential expression analyses were performed using DESeq2.To ensure a fair comparison of samples, which may contain different proportions of life cycle stages, the performance of two different *in silico* approaches was evaluated by counting Giemsa-stained thin blood smears (blue).

## Improving *var* transcript assembly, annotation and quantification

A laboratory and a clinical dataset were used to assess the performance of the different *var* assembly pipelines (Figure 2). The laboratory dataset was a *P. falciparum* 3D7 time course RNA-sequencing dataset (European nucleotide archive (ENA): PRJEB31535) (Wichers *et al*., 2019). The clinical dataset contained samples from 32 adult malaria patients, hospitalised in Hamburg, Germany (National Center for Biotechnology Information (NCBI) BioProject ID: PRJNA679547). Fifteen were malaria naïve and 17 were previously exposed to malaria. Eight of the malaria naïve patients went on to develop severe malaria and 24 had non-severe malaria (Wichers *et al*., 2021).

Our i) new whole transcript approach, ii) domain assembly approach, and iii) modified version of the original approach (see material and methods) were first applied to a *P. falciparum* 3D7 time course RNA-sequencing dataset to benchmark their performance (Wichers *et al*., 2019) (Figure 2 – Figure supplement 1). The whole transcript approach performed best, achieving near perfect alignment scores for the dominantly expressed *var* gene (Figure 2 – Figure supplement 1a). The domain and the original approach produced shorter contigs and required more contigs to assemble the *var* transcripts at the 8 and 16 hour post-invasion time points, when *var* gene expression is maximal (Figure 2 – Figure supplement 1c, f, g and h). However, we found high accuracies (> 0.95) across all approaches, meaning the sequences we assembled were correct (Figure 2 – Figure supplement 1b). The whole transcript approach also performed the best when assembling the lower expressed *var* genes (Figure 2 – Figure supplement 1e) and produced the fewest *var* chimeras compared to the original approach on *P. falciparum* 3D7. Fourteen misassemblies were observed with the whole transcript approach compared to 19 with the original approach (Table S1). This reduction in misassemblies was particularly apparent in the ring-stage samples.

Next, the assembled transcripts produced from the original approach of Wichers *et al*., 2021 were compared to those produced from our new whole transcript and domain assembly approaches for *ex vivo* samples from German travellers. Summary statistics are shown in Table 2. The whole transcript approach produced the fewest transcripts, but of greater length than the domain approach and the original approach (Figure 2 – Figure supplement 2). The whole transcript approach also returned the largest N50 score (more than doubling the N50 of the original approach), which means that it was the most contiguous assembly produced. Remarkably, with the new whole transcript method, we observed a significant decrease (2 vs 336) in clearly misassembled transcripts with, for example, an N-terminal domain at an internal position.

**Table 2:** Statistics for the different approaches used to assemble the *var* transcripts. *^Var^* ^assembly approaches were applied to malaria patient *ex vivo* samples (n=32) from (Wichers *et al*., 2021) and statistics determined. Given are the total number of assembled *var* transcripts longer than 500 nt containing at least one significantly annotated *var* domain, the maximum length of the longest assembled *var* transcript in nucleotides and the N50 value, respectively. The N50 is defined as the sequence length of the shortest *var* contig, with all *var* contigs greater than or equal to this length together accounting for 50% of the total length of concatenated *var* transcript assemblies. Misassemblies represents the number of misassemblies for each approach. *Number of misassemblies were not determined for the domain approach due to its poor performance in other metrics.^

|  | Number of contigs<br>≥500nts | Maximum length<br>(nt) | Average contig<br>length (nt) | N50 | Number of<br>misassemblies |
| --- | --- | --- | --- | --- | --- |
| Original approach | 6,441 | 10,412 | 1,621 | 2,302 | 336 |
| Domain approach | 4,691 | 5,003 | 954 | 1,088 | NA* |
| Whole transcript<br>approach | 3,011 | 12,586 | 2,771 | 5,381 | 2 |

When genome sequencing is not available, concordance of different *var* profiling approaches can support the validation of an approach. Here, the same methods used in the original analysis were applied for quantifying the expression of the assembled *var* transcripts and domains. This suggests any concordance in expression estimates likely reflects concordance at the domain annotation level. The original approach and the new whole transcript approach gave similar results for domain expression in each sample with greater correlation in results observed between the highly expressed domains (Figure 2 – Figure supplement 3). As expected, comparable results were also seen for the differentially expressed transcripts identified in the original analysis between the naïve vs pre-exposed and severe vs non-severe comparisons, respectively (Figure 2 – Figure supplement 4).

Overall, the new whole transcript approach performed the best on the laboratory 3D7 dataset (ENA: PRJEB31535) (Wichers *et al*., 2019), had the greatest N50, the longest *var* transcripts and produced concordant results with the original analysis on the clinical *ex vivo* samples (NCBI: PRJNA679547) (Wichers *et al*., 2021). Therefore, it was selected for all subsequent analyses unless specified otherwise.

## Establishing characterisation of *var* transcripts from *ex vivo* and *in vitro* samples

Of the 32 clinical isolates of *P. falciparum* from the German traveller dataset, 13 underwent one replication cycle of *in vitro* culture, 10 of these underwent a second generation and one underwent a third generation (Table 3). Most (9/13, 69%) isolates entering culture had a single MSP1 genotype, indicative of monoclonal infections. All samples were sequenced with a high read depth, although the *ex vivo* samples had a greater read depth than the *in vitro* samples (Table 3). Figure 3 shows a summary of the analysis performed.

**Table 3:**
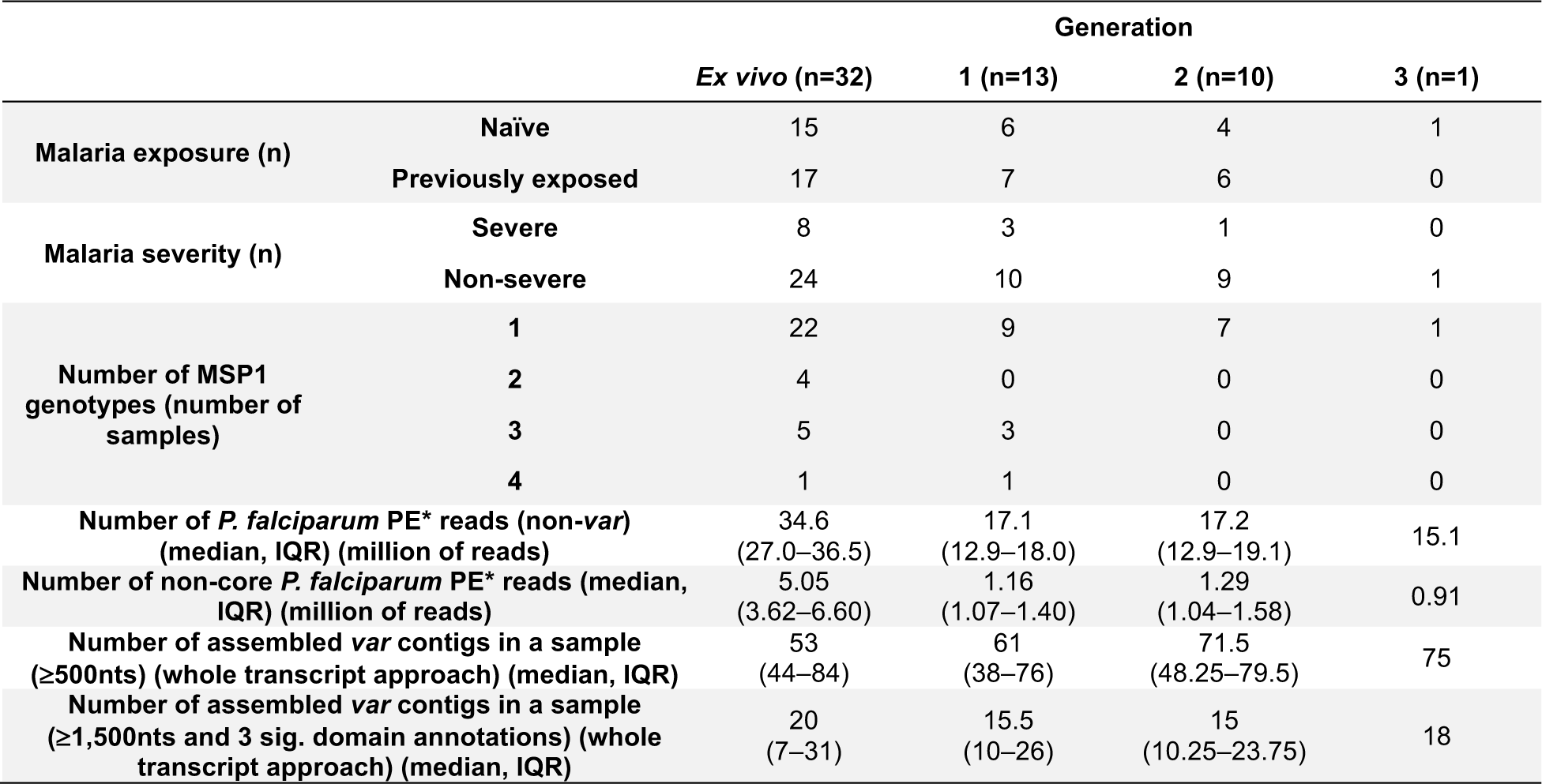
Summary of the clinical dataset used to analyze the impact of parasite culturing on gene expression. ^RNA-sequencing was performed on 32 malaria infected German traveler samples (Wichers *et al*., 2021). The 32 *ex vivo* samples were used to compare the performance of the *var* assembly approaches. Parasites from 13 of these *ex vivo* samples underwent one cycle of *in vitro* replication, 10 parasite samples were also subjected to a second cycle of replication *in vitro*, and a single parasite isolate was also analyzed after a third cycle of replication. For the *ex vivo* vs short-term *in vitro* cultivation analysis only paired samples were used. The number of assembled *var* contigs represents results per sample using the whole transcript approach, and shows either the number of assembled *var* contigs significantly annotated as *var* gene and > =500nt in length, or the number of assembled *var* transcripts identified with a length >= 1500nt and containing at least 3 significantly annotated *var* domains. *PE; paired-end reads.^

To account for differences in parasite developmental stage within each sample, which are known to impact gene expression levels (Bozdech *et al*., 2003), the proportions of life cycle stages were estimated using the mixture model approach of the original analysis (Tonkin-Hill *et al*., 2018, Wichers *et al*., 2021). As a complementary approach, single cell differential composition analysis (SCDC) with the Malaria Cell Atlas as a reference was also used to determine parasite age (Dong *et al*., 2021, Howick *et al*., 2019). SCDC and the mixture model approaches produced concordant estimates that most parasites were at ring stage in all *ex vivo*and *in vitro*samples (Figure 3 – Figure supplement 1a,b). Whilst there was no significant difference in ring stage proportions across the generations, we observed a slight increase in parasite age in the cultured samples. Overall, there were more rings and early trophozoites in the *ex vivo* samples compared to the cultured parasite samples and an increase of late trophozoite, schizont and gametocyte proportions during the culturing process (Figure 3 – Figure supplement 1c). The estimates produced from the mixture model approach showed high concordance with those observed by counting Giemsa-stained blood smears (Figure 3 – Figure supplement 1d). Due to the potential confounding effect of differences in stage distribution on gene expression, we adjusted for developmental stage determined by the mixture model in all subsequent analyses.

Our new approach was applied to RNA-sequencing samples of *ex vivo* and short-term *in vitro* cultured parasites from German travellers (Wichers *et al*., 2021). Table S2 shows the assembled *var* transcripts on a per sample basis. Interestingly, we observed SSPACE did not provide improvement in terms of extending *var* assembled contigs in 9/37 samples. We observed a significant increase in the number of assembled *var* transcripts in generation 2 parasites compared to paired generation 1 parasites (p_adj_ = 0.04, paired Wilcoxon test). We observed no significant differences in the length of the assembled *var* transcripts across the generations. Three different filtering approaches were applied in comparison to maximise the likelihood that correct assemblies were taken forward for further analysis and to avoid the overinterpretation of lowly expressed partial *var* transcripts (Table S3). Filtering for *var* transcripts at least 1500nt long and containing at least 3 significantly annotated *var* domains was the least restrictive, while the other approaches required the presence of a DBLα domain within the transcript. All three filtering approaches generated the same maximum length *var* transcript and similar N50 values. This suggests minimal differences in the three filtering approaches, whilst highlighting the importance of filtering assembled *var* transcripts.

In the original approach of Wichers *et al*., 2021, the non-core reads of each sample used for *var* assembly were mapped against a pooled reference of assembled *var* transcripts from all samples, as a preliminary step towards differential *var* transcript expression analysis. This approach returned a small number of *var* transcripts which were expressed across multiple patient samples (Figure 3 – Figure supplement 2a). As genome sequencing was not available, it was not possible to know whether there was truly overlap in *var* genomic repertoires of the different patient samples, but substantial overlap was not expected. Stricter mapping approaches (for example, excluding transcripts shorter than 1500nt) changed the resulting *var* expression profiles and produced more realistic scenarios where similar *var* expression profiles were generated across paired samples, whilst there was decreasing overlap across different patient samples (Figure 3 – Figure supplement 2b,c). Given this limitation, we used the paired samples to analyse *var* gene expression at an individual subject level, where we confirmed the MSP1 genotypes and alleles were still present after short-term *in vitro*cultivation. The per patient approach showed consistent expression of *var* transcripts within samples from each patient but no overlap of *var* expression profiles across different patients (Figure 3 – Figure supplement 2d). Taken together, the per patient approach was better suited for assessing *var* transcriptional changes in longitudinal samples. However, it has been hypothesised that more conserved *var* genes in field isolates increase parasite fitness during chronic infections, necessitating the need to correctly identify them (Dimonte *et al*., 2020, Otto *et al*., 2019). Accordingly, further work is needed to optimise the pooled sample approach to identify truly conserved *var* transcripts across different parasite isolates in cross-sectional studies.

## Longitudinal analysis of *var* transcriptome from *ex vivo* to *in vitro* samples

To assess the changes in the *var* transcriptome induced by parasite culturing, we performed a series of analyses, all of which addressed different aspects: (i) changes in individual *var* gene expression pattern and *var* expression homogeneity (“per patient analysis”), (ii) changes in the expression of *var* variants conserved between strains, (iii) changes in the expression of PfEMP1 domains, (iv) changes in expression at the *var* group level, and (v) at the overall *var* expression level. We validated our results using the DBLα-tag approach and complemented the *var* -specific analysis by also examining changes in the core transcriptome.

To investigate whether dominant *var* gene expression changes through *in vitro* culture, rank analysis of *var* transcript expression was performed (Figure 4, Figure 4 – Figure supplement 1). In most cases a single dominant *var* transcript was detected. The dominant *var* gene did not change in most patient samples and the ranking of *var* gene expression remained similar. However, we observed a change in the dominant *var* gene being expressed through culture in isolates from three of 13 (23%) patients (#6, #17 and #26). Changes in the dominant *var* gene expression were also observed in the DBLα-tag data for these patients (described below). In parasites from three additional patients, #1, #7 and #14, the top expressed *var* gene remained the same, however we observed a change in the ranking of other highly expressed *var* genes in the cultured samples compared to the *ex vivo* sample. Interestingly, in patient #26 we observed a switch from a dominant group A *var* gene to a group B and C *var* gene. This finding was also observed in the DBLα-tag analysis (results below). A similar finding was seen in patient #7. In the *ex vivo* sample, the second most expressed *var* transcript was a group A transcript. However, in the cultured samples expression of this transcript was reduced and we observed an increase in the expression of group B and C *var* transcripts. A similar pattern was observed in the DBLα-tag analysis for patient #7, whereby the expression of a group A transcript was reduced during the first cycle of cultivation. Overall, the data suggest that some patient samples underwent a larger *var* transcriptional change when cultured compared to the other patient samples and that culturing parasites can lead to an unpredictable *var* transcriptional change.

**Figure 4:**
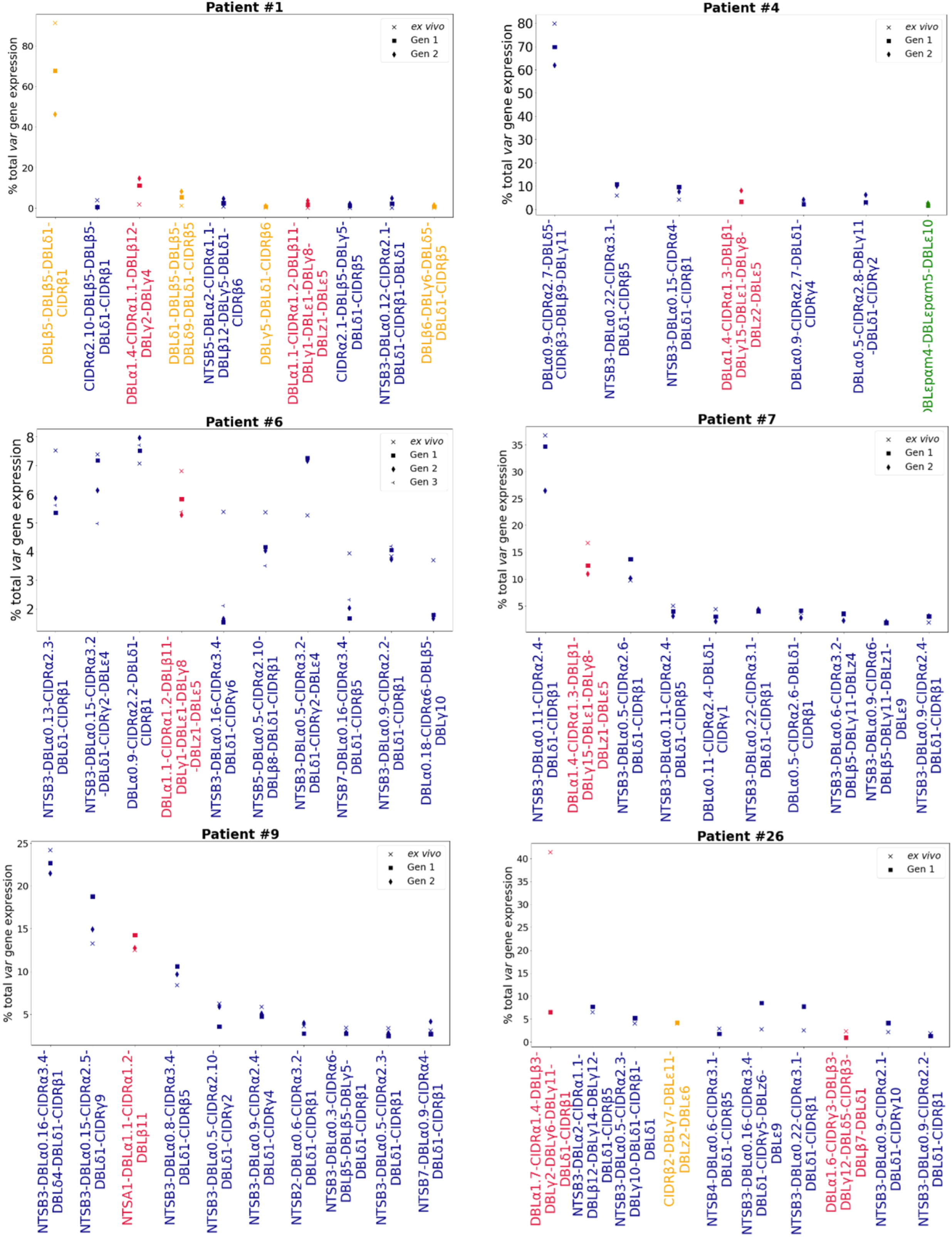
Rank *var* gene expression analysis. For each patient, the paired *ex vivo* (n=13) and *in vitro* samples (generation 1: n=13, generation 2: n=10, generation 3: n=1) were analysed. The assembled *var* transcripts with at least 1500nt and containing 3 significantly annotated *var* domains across all the generations for a patient were combined into a reference, redundancy was removed using cd-hit (at sequence identity = 99%), and expression was quantified using Salmon. *Var* transcript expression was ranked. Plots show the top 10 *var* gene expression rankings for each patient and their *ex vivo* and short-term *in vitro* cultured parasite samples. Group A *var* transcripts (red), group B or C *var* transcripts (blue), group E *var* transcripts (green) and transcripts of unknown *var* group (orange).

In line with these results, *var* expression homogeneity (VEH) on a per patient basis showed in some patients a clear change, with the *ex vivo* sample diversity curve distinct from those of *in vitro* generation 1 and generation 2 samples (patients #1, #2, #4) (Figure 4 – Figure supplement 2). Similarly, in other patient samples, we observed a clear difference in the curves of *ex vivo* and generation 1 samples (patient #25 and #26, both from first-time infected severe malaria patients). Some of these samples (#1 and #26, both from first-time infected severe malaria patients) also showed changes in their dominant *var* gene expression during culture, taken together indicating much greater *var* transcriptional changes *in vitro* compared to the other samples.

### Expression of conserved *var* gene variants through short-term *in vitro* culture

Due to the relatively high level of conservation observed in *var1, var2csa* and *var3*, they do not present with the same limitations as regular *var* genes. Therefore, changes in their expression through short-term culture was investigated across all samples together. We observed no significant differences in the expression of conserved *var* gene variants, *var1-IT* (p_adj_ *=* 0.61, paired Wilcoxon test)*, var1 -3D7* (p_adj_ =0.93, paired Wilcoxon test) and *var2csa* (p_adj_ =0.54, paired Wilcoxon test) between paired *ex vivo* and generation 1 parasites, but *var2csa* was significantly differentially expressed between generation 1 and generation 2 parasites (p_adj_ = 0.029, paired Wilcoxon test) (Figure 4 – Figure supplement 3). However, *var2csa* expression previously appeared to have decreased in some paired samples during the first cycle of cultivation (Figure 4 – Figure supplement 3).

### Differential expression of *var* domains from *ex vivo* to *in vitro* samples

There is overlap in PfEMP1 domain subtypes of different parasite isolates which can be associated with *var* gene groups and receptor binding phenotypes. This allows performing differential expression analysis on the level of encoded PfEMP1 domain subtypes, as done in previous studies (Tonkin-Hill *et al*., 2018, Wichers *et al*., 2021). PCA on *var* domain expression (Figure 5a) showed some patients’ *ex vivo* samples clustering away from their respective generation 1 sample (patient #1, #2, #4, #12, #17, #25), again indicating a greater *var* transcriptional change relative to the other samples during the first cycle of cultivation. However, in the pooled comparison of the generation 1 vs *ex vivo* of all isolates, a single domain was significantly differentially expressed, CIDRα2.5 associated with B-type PfEMP1 proteins and CD36-binding (Figure 5b). In the generation 2 vs *ex vivo* comparison, there were no domains significantly differentially expressed, however we observed large log_2_FC values in similar domains to those changing most in the *ex vivo*vs generation 1 comparison (Figure 5c). No differentially expressed domains were found in the generation 1 vs generation 2 comparison. These results suggest individual changes in *var* expression are not reflected in the pooled analysis and the per patient approach is more suitable.

**Figure 5:**
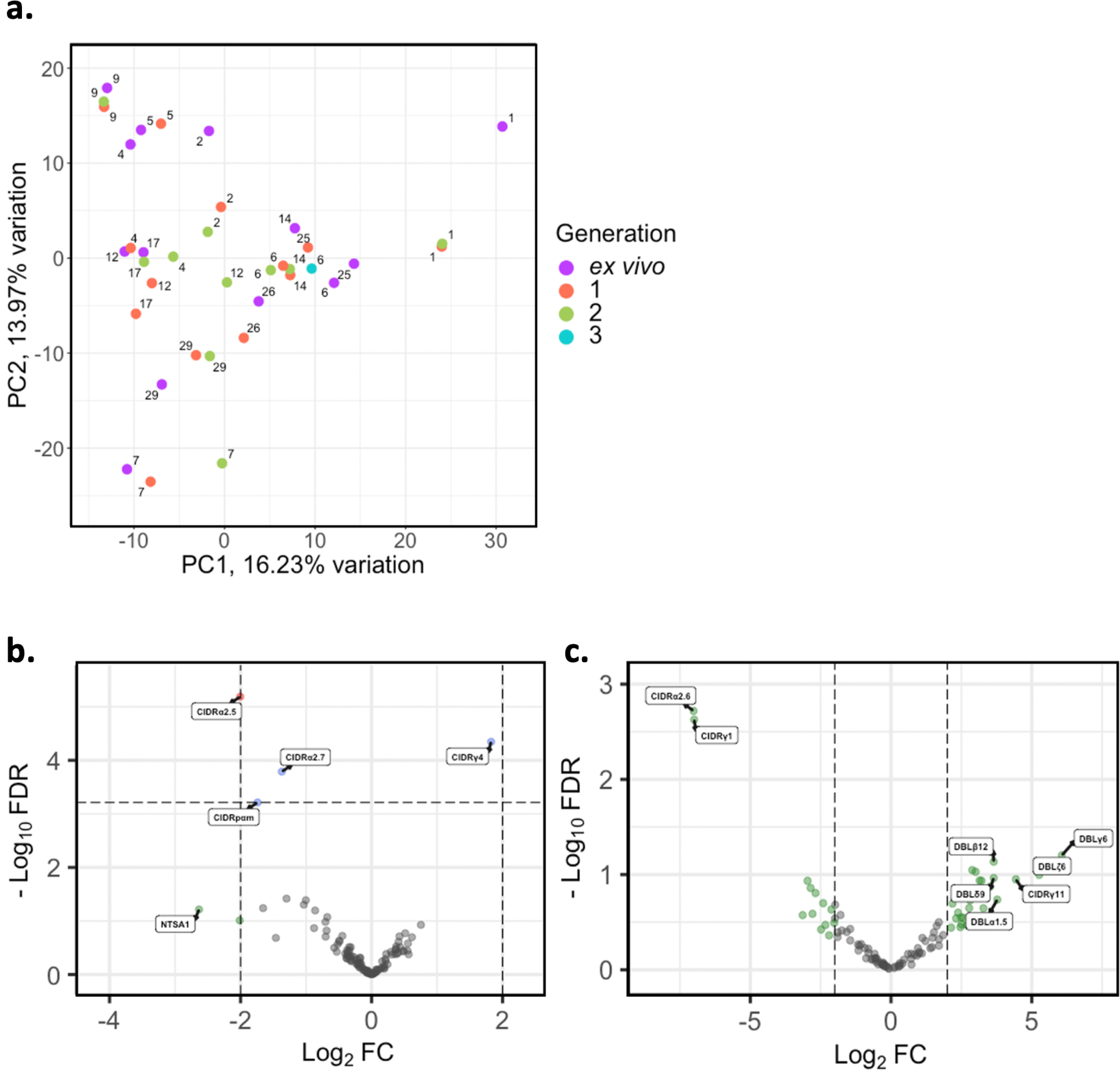
*Var* domain transcriptome analysis through short-term *in vitro* culture. *Var* transcripts for paired *ex vivo* (n=13), generation 1 (n=13), generation 2 (n=10) and generation 3 (n=1) were *de novo* assembled using the whole transcript approach. *Var* transcripts were filtered for those >= 1500nt in length and containing at least 3 significantly annotated *var* domains. Transcripts were annotated using HMM models built on the Rask et al., 2010 dataset (Rask *et al*., 2010). When annotating the whole transcript, the most significant alignment was taken as the best annotation for each region of the assembled transcript (e-value cut off 1e-5). Multiple annotations were allowed on the transcript if they were not overlapping, determined using cath-resolve-hits. *Var* domain expression was quantified using FeatureCounts and the domain counts aggregated **a)** PCA plot of log_2_ normalized read counts (adjusted for life cycle stage, derived from the mixture model approach). Points are coloured by their generation (*ex vivo*; purple, generation 1; red, generation 2; green and generation 3; blue) and labelled by their patient identity **b)** Volcano plot showing extent and significance of up-or down-regulation of *var* domain expression in *ex vivo* (n=13) compared with paired generation 1 cultured parasites (n=13) (red and blue, *P* < 0.05 after Benjamini-Hochberg adjustment for FDR; red and green, absolute log_2_ fold change log_2_FC in expression >= 2). Domains with a log_2_FC > = 2 represent those upregulated in generation 1 parasites. Domains with a log_2_FC <= -2 represent those downregulated in generation 1 parasites. **c)** Volcano plot showing extent and significance of up-or down-regulation of *var* domain expression in *ex vivo* (n=10) compared with paired generation 2 cultured parasites (n=10) (green, absolute log_2_ fold change log_2_FC in expression >= 2). Domains with a log_2_FC >= 2 represent those upregulated in generation 2 parasites. Domains with a log_2_FC <= -2 represent those downregulated in generation 2 parasites. Differential expression analysis was performed using DESeq2 (adjusted for life cycle stage, derived from the mixture model approach).

### *Var* group expression analysis

A previous study found group A *var* genes to have a rapid transcriptional decline in culture compared to group B *var* genes, however another study found a decrease in both group A and group B *var* genes in culture (Zhang *et al*., 2011, Peters *et al*., 2007). These studies were limited as the *var* type was determined by analysing the sequence diversity of DBLα domains, and by quantitative PCR (qPCR) methodology which restricts analysis to quantification of known/conserved sequences. Due to these results, the expression of group A *var* genes vs. group B and C *var* genes was investigated using a paired analysis on all the DBLα (DBLα1 vs DBLα0 and DBLα2) and NTS (NTSA vs NTSB) sequences assembled from *ex vivo* samples and across multiple generations in culture. A linear model was created with group A expression as the response variable, the generation and life cycle stage as independent variables and the patient information included as a random effect. The same was performed using group B and C expression levels.

In both approaches, DBLα and NTS, we found no significant changes in total group A or group B and C *var* gene expression levels (Figure 6). We observed high levels of group B and C *var* gene expression compared to group A in all patients, both in the *ex vivo* samples and the *in vitro* samples. In some patients we observed a decrease in group A *var* genes from *ex vivo*to generation 1 (patients #1, #2, #5, #6, #9, #12, #17, #26) (Figure 6a), however in all but four patients (patient #1, #2, #5, #6) the levels of group B and C *var* genes remained consistently high from *ex vivo* to generation 1 (Figure 6b). Interestingly, patients #6 and #17 also had a change in the dominant *var* gene expression through culture. Taken together with the preceding results, it appears that observed differences in *var* transcript expression occurring with transition to short-term culture are not due to modulation of recognised *var* classes, but due to differences in expression of particular *var* transcripts.

**Figure 6.**
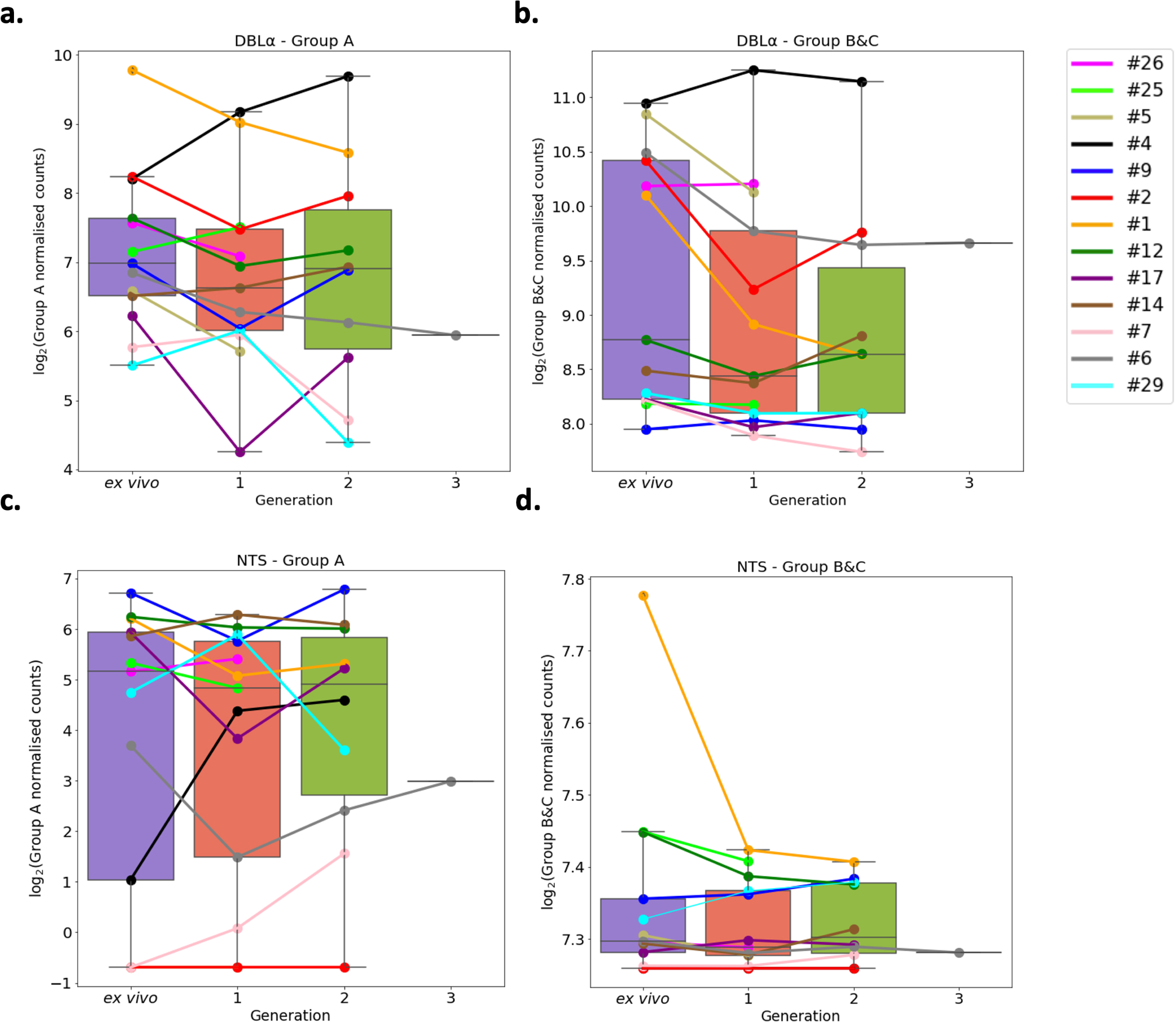
*Var* group expression analysis through short-term *in vitro* culture. The DBLα domain sequence for each transcript was determined and for each patient a reference of all assembled DBLα domains combined. Group A *var* genes possess DBLα1 domains, some group B encode DBLα2 domains and groups B and C encode DBLα0 domains. Domains were grouped by type and their expression summed. The relevant sample’s non-core reads were mapped to this using Salmon and DBLα expression quantified. DESeq2 normalisation was performed, with patient identity and life cycle stage proportions included as covariates. A similar approach was repeated for NTS domains. Group A *var* genes encode NTSA compared to group B and C *var* genes which encode NTSB. Boxplots show log_2_ normalised Salmon read counts for **a)** group A *var* gene expression through cultured generations assessed using the DBLα domain sequences, **b)** group B and C *var* gene expression through cultured generations assessed using the DBLα domain sequences, **c)** group A *var* gene expression through cultured generations assessed using the NTS domain sequences, and **d)** group B and C *var* gene expression through cultured generations assessed using the NTS domain sequences. Different coloured lines connect paired patient samples through the generations: *ex vivo* (n=13), generation 1 (n=13), generation 2 (n=10) and generation 3 (n=1). Axis shows different scaling.

### Quantification of total *var* gene expression

We observed a trend of decreasing total *var* gene expression between generations irrespective of the assembler used in the analysis (Figure 6 – Figure supplement 1). A similar trend is seen with the LARSFADIG count, which is commonly used as a proxy for the number of different *var* genes expressed (Otto *et al*., 2019). A linear model was created (using only paired samples from *ex vivo* and generation 1) (Supplementary file 1) with proportion of total gene expression dedicated to *var* gene expression as the response variable, the generation and life cycle stage as independent variables and the patient information included as a random effect. This model showed no significant differences between generations, suggesting that differences observed in the raw data may be a consequence of small changes in developmental stage distribution in culture.

### Validation of *var* expression profiling by DBLα-tag sequencing

Deep sequencing of RT-PCR-amplified DBLα expressed sequence tags (ESTs) combined with prediction of the associated transcripts and their encoded domains using the Varia tool (Mackenzie *et al*., 2022) was performed to supplement the RNA-sequencing analysis. The raw Varia output file is given in Supplementary file 2. Overall, we found a high agreement between the detected DBLα-tag sequences and the *de novo* assembled *var* transcripts. A median of 96% (IQR: 93–100%) of all unique DBLα-tag sequences detected with >10 reads were found in the RNA-sequencing approach. This is a significant improvement on the original approach (p= 0.0077, paired Wilcoxon test), in which a median of 83% (IQR: 79–96%) was found (Wichers *et al*., 2021). To allow for a fair comparison of the >10 reads threshold used in the DBLα-tag approach, the upper 75^th^ percentile of the RNA-sequencing-assembled DBLα domains were analysed. A median of 77.4% (IQR: 61–88%) of the upper 75^th^ percentile of the assembled DBLα domains were found in the DBLα-tag approach. This is a lower median percentage than the median of 81.3% (IQR: 73– 98%) found in the original analysis (p= 0.28, paired Wilcoxon test) and suggests the new assembly approach is better at capturing all expressed DBLα domains.

The new whole transcript assembly approach also had high consistency with the domain annotations predicted from Varia. Varia predicts *var* sequences and domain annotations based on short sequence tags, using a database of previously defined *var* sequences and annotations (Mackenzie *et al*., 2022). A median of 85% of the DBLα annotations and 73% of the DBLα-CIDR domain annotations, respectively, identified using the DBLα-tag approach were found in the RNA sequencing approach. This further confirms the performance of the whole transcript approach and it was not restricted by the pooled approach of the original analysis. We also observed consistent results with the per patient analysis, in terms of changes in the dominant *var* gene expression (described above) (Supplementary file 2). In line with the RNA-sequencing data, the DBLα-tag approach revealed no significant differences in Group A and Group B and C groups during short-term culture, further highlighting the agreement of both methods (Figure 6 – Figure supplement 2).

## Differential expression analysis of the core transcriptome between *ex vivo* and *in vitro* samples

Given the modest changes in *var* gene expression repertoire upon culture we wanted to investigate the extent of any accompanying changes in the core parasite transcriptome. PCA was performed on core gene (*var, rif, stevor, surf* and rRNA genes removed) expression, adjusted for life cycle stage. We observed distinct clustering of *ex vivo*, generation 1, and generation 2 samples, with patient identity having much less influence (Figure 7a). There was also a change from the heterogeneity between the *ex vivo* samples to more uniform clustering of the generation 1 samples (Figure 7a), suggesting that during the first cycle of cultivation the core transcriptomes of different parasite isolates become more alike.

**Figure 7:**
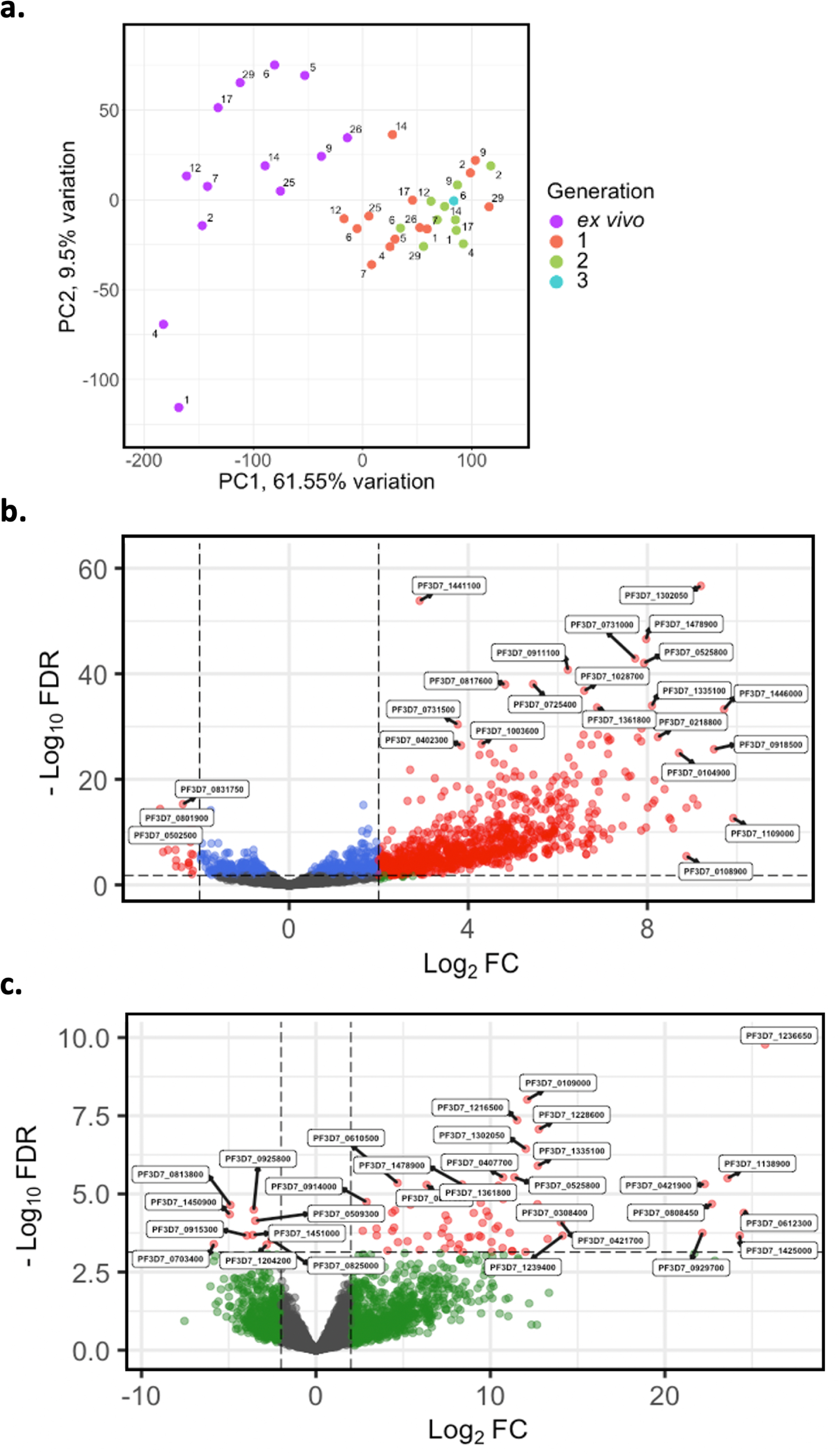
Core gene transcriptome analysis of *ex vivo* and short-term *in vitro* cultured samples. Core gene expression was assessed for paired *ex vivo* (n=13), generation 1 (n=13), generation 2 (n=10) and generation 3 (n=1) parasite samples. Subread align was used, as in the original analysis, to align the reads to the human genome and *P. falciparum* 3D7 genome, *with var, rif, stevor, surf* and *rRNA* genes removed. HTSeq count was used to quantify gene counts. **a)** PCA plot of log_2_ normalized read counts. Points are coloured by their generation (*ex vivo*: purple, generation 1: red, generation 2: green, and generation 3: blue) and labelled by their patient identity. **b)** Volcano plot showing extent and significance of up- or down-regulation of core gene expression in *ex vivo*(n=13) compared with paired generation 1 cultured parasites (n=13) and **c)** in *ex vivo* (n=10) compared with paired generation 2 cultured parasites (n=10). Dots in red and blue represent those genes with *P* < 0.05 after Benjamini-Hochberg adjustment for FDR, red and green dots label genes with absolute log_2_ fold change log_2_FC in expression >= 2). Accordingly, genes with a log_2_FC > = 2 represent those upregulated in generation 1 parasites and genes with a log_2_FC <= -2 represent those downregulated in generation 1 parasites. Normalized read counts of the core gene analysis were adjusted for life cycle stage, derived from the mixture model approach.

In total, 920 core genes (19% of the core transcriptome) were found to be differentially expressed after adjusting for life cycle stages using the mixture model approach between *ex vivo and* generation 1 samples (Supplementary file 3). The majority were upregulated, indicating a substantial transcriptional change during the first cycle of *in vitro* cultivation (Figure 7b). 74 genes were found to be upregulated in generation 2 when compared to the *ex vivo* samples, many with log_2_FC greater than those in the *ex vivo* vs generation 1 comparison (Figure 7c). No genes were found to be significantly differentially expressed between generation 1 and generation 2. However, five genes had a log_2_FC >= 2 and were all upregulated in generation 2 compared to generation 1. Interestingly, the gene with the greatest fold change, encoding ROM3 (PF3D7_0828000), was also found to be significantly downregulated in generation 1 parasites in the *ex vivo* vs generation 1 analysis. The other four genes were also found to be non-significantly downregulated in generation 1 parasites in the *ex vivo* vs generation 1 analysis. This suggests changes in gene expression during the first cycle of cultivation are the greatest compared to the other cycles.

The most significantly upregulated genes (in terms of fold change) in generation 1 contained several small nuclear RNAs, splicesomal RNAs and non-coding RNAs (ncRNAs). 16 ncRNAs were found upregulated in generation 1, with several RNA-associated proteins having large fold changes (log_2_FC > 7) Significant gene ontology (GO) terms and Kyoto encyclopedia of genes and genomes (KEGG) pathways for the core genes upregulated in generation 1 included “entry into host”, “movement into host” and “cytoskeletal organisation” suggesting the parasites undergo a change in invasion efficiency, which is connected to the cytoskeleton, during their first cycle of *in vitro* cultivation (Figure 7 – Figure supplement 1). We observed eight AP2 transcription factors upregulated in generation 1 (PF3D7_0404100/AP2-SP2, PF3D7_0604100/SIP2, PF3D7_0611200/AP2-EXP2, PF3D7_0613800, PF3D7_0802100/AP2-LT, PF3D7_1143100/AP2-O, PF3D7_1239200, PF3D7_1456000/AP2-HC) with no AP2 transcription factors found to be downregulated in generation 1. To confirm the core gene expression changes identified were not due to the increase in parasite age during culture, as indicated by upregulation of many schizont-related genes, core gene differential expression analysis was performed on paired *ex vivo* and generation 1 samples that contained no schizonts or gametocytes in generation 1. The same genes were identified as significantly differentially expressed with a Spearman’s rank correlation of 0.99 for the log_2_FC correlation between this restricted sample approach and those produced using all samples (Figure 7 – Figure supplement 2).

### Cultured parasites as surrogates for assessing the *in vivo* core gene transcriptome

In the original analysis of *ex vivo samples*, hundreds of core genes were identified as significantly differentially expressed between pre-exposed and naïve malaria patients. We investigated whether these differences persisted after *in vitro* cultivation. We performed differential expression analysis comparing parasite isolates from naïve (n=6) vs pre-exposed (n=7) patients, first between their *ex vivo* samples, and then between the corresponding generation 1 samples. Interestingly, when using the *ex vivo* samples, we observed 206 core genes significantly upregulated in naïve patients compared to pre-exposed patients (Figure 7 – Figure supplement 3a). Conversely, we observed no differentially expressed genes in the naïve vs pre-exposed analysis of the paired generation 1 samples (Figure 7 – Figure supplement 3b). Taken together with the preceding findings, this suggests one cycle of cultivation shifts the core transcriptomes of parasites to be more alike each other, diminishing inferences about parasite biology *in vivo*.

A summary describing the rationale, results and interpretation of each approach in our analysis pipeline can be found in Table 4.

**Table 4:**
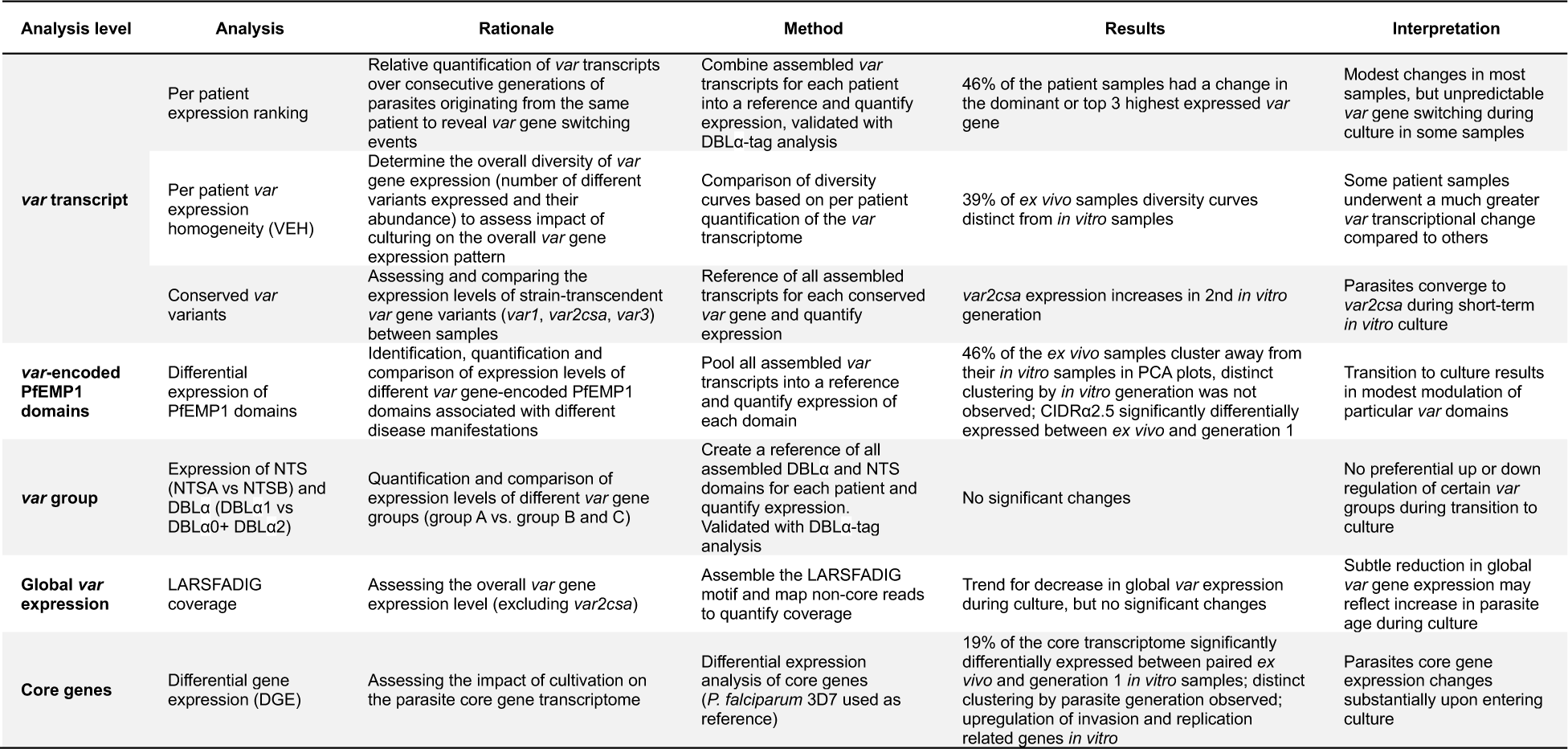
Summary of the different levels of analysis performed to assess the effect of short-term parasite culturing on *var* and core gene expression, their rational, method, results, and interpretation.

## Discussion

Multiple lines of evidence point to PfEMP1 as a major determinant of malaria pathogenesis, but previous approaches for characterising *var* expression profiles in field samples have limited *in vivo* studies of PfEMP1 function, regulation, and association with clinical symptoms (Tarr *et al*., 2018, Lee *et al*., 2018, Warimwe *et al*., 2013, Rorick *et al*., 2013, Zhang *et al*., 2011, Taylor *et al*., 2002). A more recent approach, based on RNA-sequencing, overcame many of the limitations imposed by the previous primer-based methods (Tonkin-Hill *et al*., 2018, Wichers *et al*., 2021). However, depending on the expression level and sequencing depth, *var* transcripts were found to be fragmented and only a partial reconstruction of the *var* transcriptome was achieved (Tonkin-Hill *et al*., 2018, Wichers *et al*., 2021, Andrade *et al*., 2020, Guillochon *et al*., 2022, Yamagishi *et al*., 2014). The present study developed a novel approach for *var* gene assembly and quantification that overcomes many of these limitations.

Our new approach used the most geographically diverse reference of *var* gene sequences to date, which improved the identification of reads derived from *var* transcripts. This is crucial when analysing patient samples with low parasitaemia where *var* transcripts are hard to assemble due to their low abundancy (Guillochon *et al*., 2022). Our approach has wide utility due to stable performance on both laboratory-adapted and clinical samples. Concordance in the different *var* expression profiling approaches (RNA-sequencing and DBLα-tag) on *ex vivo* samples increased using the new approach by 13%, when compared to the original approach (96% in the whole transcript approach compared to 83% in Wichers *et al*., 2021. This suggests the new approach provides a more accurate method for characterising *var* genes, especially in samples collected directly from patients. Ultimately, this will allow a deeper understanding of relationships between *var* gene expression and clinical manifestations of malaria.

Having a low number of long contigs is desirable in any *de novo* assembly. This reflects a continuous assembly, as opposed to a highly fragmented one where polymorphic and repeat regions could not be resolved (Lischer & Shimizu, 2017). An excessive number of contigs cannot be reasonably handled computationally and results from a high level of ambiguity in the assembly (Yang *et al*., 2012). We observed a greater than 50% reduction in the number of contigs produced in our new approach, which also had a 21% increase in the maximum length of the assembled *var* transcripts, when compared to the original approach. It doubled the assembly continuity and assembled an average of 13% more of the *var* transcripts. This was particularly apparent in the N-terminal region, which has often been poorly characterised by existing approaches. The original approach failed to assemble the N-terminal region in 58% of the samples, compared to just 4% in the new approach with assembly consistently achieved with an accuracy > 90%. This is important because the N-terminal region is known to contribute to the adhesion phenotypes of most PfEMP1 proteins.

The new approach allows for *var* transcript reconstruction across a range of expression levels, which is required when characterising *var* transcripts from multi-clonal infections. Assembly completeness of the lowly expressed *var* genes increased five-fold using the new approach. Biases towards certain parasite stages have been observed in non-severe and severe malaria cases, so it is valuable to assemble the *var* transcripts from different life cycle stages (Tonkin-Hill *et al*., 2018). Our new approach is not limited by parasite stage. It was able to assemble the whole *var* transcript, in a single contig, at later stages in the *P. falciparum* 3D7 intra-erythrocytic cycle, something previously unachievable. The new approach allows for a more accurate and complete picture of the *var* transcriptome. It provides new perspectives for relating *var* expression to regulation, co-expression, epigenetics and malaria pathogenesis. It can be applied for example in analysis of patient samples with different clinical outcomes and longitudinal tracking of infections *in vivo*. It represents a crucial improvement for quantifying the *var* transcriptome. In this work, the improved approach for *var* gene assembly and quantification was used to characterise *var* gene expression during transition from *in vivo* to short-term culture.

This study had substantial power through the use of paired samples. However, many *var* gene expression studies do not have longitudinal sampling. Future work should focus on identifying the best approach for analysing the *var* transcripts in cross-sectional samples. Higher level *var* classification systems, such as the PfEMP1 predicted binding phenotype or domain cassettes, could be applied to test for over-representation of different *var* gene features in different groups of interest, because the assumption of overlapping *var* repertoires at these levels of classification would be more realistic. This was briefly explored in our analysis through *var* domain differential expression analysis, which found minimal changes in *var* domain expression through short-term culture, supporting the per patient analysis results. This could be further improved by advancing the classifications of domain subtypes. This has recently been studied using MEME to identify short nucleotide motifs that are representative of domain subtypes (Otto et al., 2019). Other research could investigate clustering *var* transcripts based on sequence identity and testing for clusters associated with specific malaria disease groups.

Studies have been performed investigating differences between long-term laboratory-adapted clones and clinical isolates, with hundreds of genes found to be differentially expressed (Hoo *et al*., 2019, Tarr *et al*., 2018, Mackinnon *et al*., 2009). Surprisingly, studies investigating the impact of short-term culture on parasites are extremely limited, despite it being commonly undertaken for making inferences about the *in vivo* transcriptome (Vignali *et al*., 2011). Using the new *var* assembly approach, we found that *var* gene expression remains relatively stable during transition to culture. However, the conserved *var2csa* had increased expression from generation 1 to generation 2. It has previously been suggested that long-term cultured parasites converge to expressing *var2csa*, but our findings suggest this begins within two cycles of cultivation (Zhang *et al*., 2022, Mok *et al*., 2008). Switching to *var2csa* has been shown to be favourable and is suggested to be the default *var* gene upon perturbation to *var* specific heterochromatin (Ukaegbu *et al*., 2015). These studies also suggested *var2csa* has a unique role in *var* gene switching and our results are consistent with the role of *var2csa* as the dominant “sink node” (Zhang *et al*., 2022, Ukaegbu *et al*., 2015, Ukaegbu *et al*., 2014, Mok *et al*., 2008). A previous study suggested *in vitro* cultivation of controlled human malaria infection samples resulted in dramatic changes in *var* gene expression (Lavstsen *et al*., 2005, Peters *et al*., 2007). Almost a quarter of samples in our analysis showed more pronounced and unpredictable changes. In these individuals, the dominant *var* gene being expressed changed within one cycle of cultivation. This implies short-term culture can result in unpredictable *var* gene expression as observed previously using a semi-quantitative RT-PCR approach (Bachmann *et al*., 2011) and that one would need to confirm *in vivo* expression matches *in vitro* expression. This can be achieved using the assembly approach described here.

We observed no generalised pattern of up- or downregulation of specific *var* groups following transition to culture. This implies there is probably not a selection event occurring during culture but may represent a loss of selection that is present *in vivo.* A global down regulation of certain *var* groups might only occur as a selective process over many cycles in extended culture. Determining changes in *var* group expression levels are difficult using degenerate qPCR primers bias and previous studies have found conflicting results in terms of changes of expression of *var* groups through cultivation. Zhang *et al*., 2011 found a rapid transcriptional decline of group A and group B *var* genes, however Peters *et al*., 2007 found group A *var* genes to have a high rate of downregulation, when compared to group B *var* genes. These studies differed in the stage distribution of the parasites and were limited in measuring enough variants through their use of primers. Our new approach allowed for the identification of more sequences, with 26.6% of assembled DBLα domains not found via the DBLα-tag approach. This better coverage of the expressed *var* diversity was not possible in these previous studies and may explain discrepancies observed.

Generally, there was a high consensus between all levels of *var* gene analysis and changes observed during short-term *in vitro* cultivation. However, the impact of short-term culture was the most apparent at the *var* transcript level and became less clear at the *var* domain, *var* type and global *var* gene expression level. This highlights the need for accurate characterisation of full length *var* transcripts and analysis of the *var* transcriptome at different levels, both of which can be achieved with the new approach developed here.

We saw striking changes in the core gene transcriptomes between *ex vivo* and generation 1 parasites with 19% of the core genome being differentially expressed. A previous study showed that expression of 18% of core genes were significantly altered after ∼50 cycles through culture (Mackinnon *et al*., 2009), but our data suggest that much of this change occurs early in the transition to culture. We observed genes with functions unrelated to ring-stage parasites were among those most significantly expressed in the generation 1 vs *ex vivo* analysis, suggesting the culture conditions may temporarily dysregulate stage-specific expression patterns or result in the parasites undergoing a rapid adaptation response (Andreadaki *et al*., 2020, Beeson *et al*., 2016). Several AP2 transcription factors (AP2-SP2, AP2-EXP2, AP2-LT, AP2-O and AP2-HC) were upregulated in generation 1. AP2-HC has been shown to be expressed in asexual parasites (Carrington *et al*., 2021). AP2-O is thought to be specific for the ookinete stage and AP2-SP2 plays a key role in sporozoite stage specific gene expression (Kaneko *et al*., 2015, Yuda *et al*., 2010). Our findings are consistent with another study investigating the impact of long-term culture (Mackinnon *et al*., 2009) which also found genes like merozoite surface proteins differentially expressed, however they were downregulated in long-term cultured parasites, whereas we found them upregulated in generation 1. This suggests short-term cultured parasites might be transcriptionally different from long-term cultured parasites, especially in their invasion capabilities, something previously unobserved. Several genes involved in the stress response of parasites were upregulated in generation 1, for example DnaJ proteins, serine proteases and ATP dependent CLP proteases (Oakley *et al*., 2007). The similarity of the core transcriptomes of the *in vitro* samples compared to the heterogeneity seen in the *ex vivo* samples could be explained by a stress response upon entry to culture. Studies investigating whether the dysregulation of stage specific expression and the expression of stress associated genes persist in long-term culture are required to understand whether they are important for growth in culture. Critically, the marked differences presented here suggest the impact of short-term culture can override differences observed in both the *in vivo* core and *var* transcriptomes of different disease manifestations.

In summary, we present an enhanced approach for *var* transcript assembly which allows for *var* gene expression to be studied in connection to *P. falciparum’s* core transcriptome through RNA-sequencing. This will be useful for expanding our understanding of *var* gene regulation and function in *in vivo* samples. As an example of the capabilities of the new approach, the method was used to quantify differences in gene expression upon short-term culture adaptation. This revealed that inferences from clinical isolates of *P. falciparum* put into short-term culture must be made with a degree of caution. Whilst *var* gene expression is often maintained, unpredictable switching does occur, necessitating that the similarity of *in vivo* and *in vitro* expression should be confirmed. The more extreme changes in the core transcriptome could have much bigger implications for understanding other aspects of parasite biology such as growth rates and drug susceptibility and raise a need for additional caution. Further work is needed to examine *var* and core transcriptome changes during longer term culture on a larger sample size. Understanding the ground truth of the *var* expression repertoire of *Plasmodium* field isolates still presents a unique challenge and this work expands the database of *var* sequences globally. The increase in long-read sequencing and the growing size of *var* gene databases containing isolates from across the globe will help overcome this issue in future studies.

## Materials and Methods

**Table 5:**
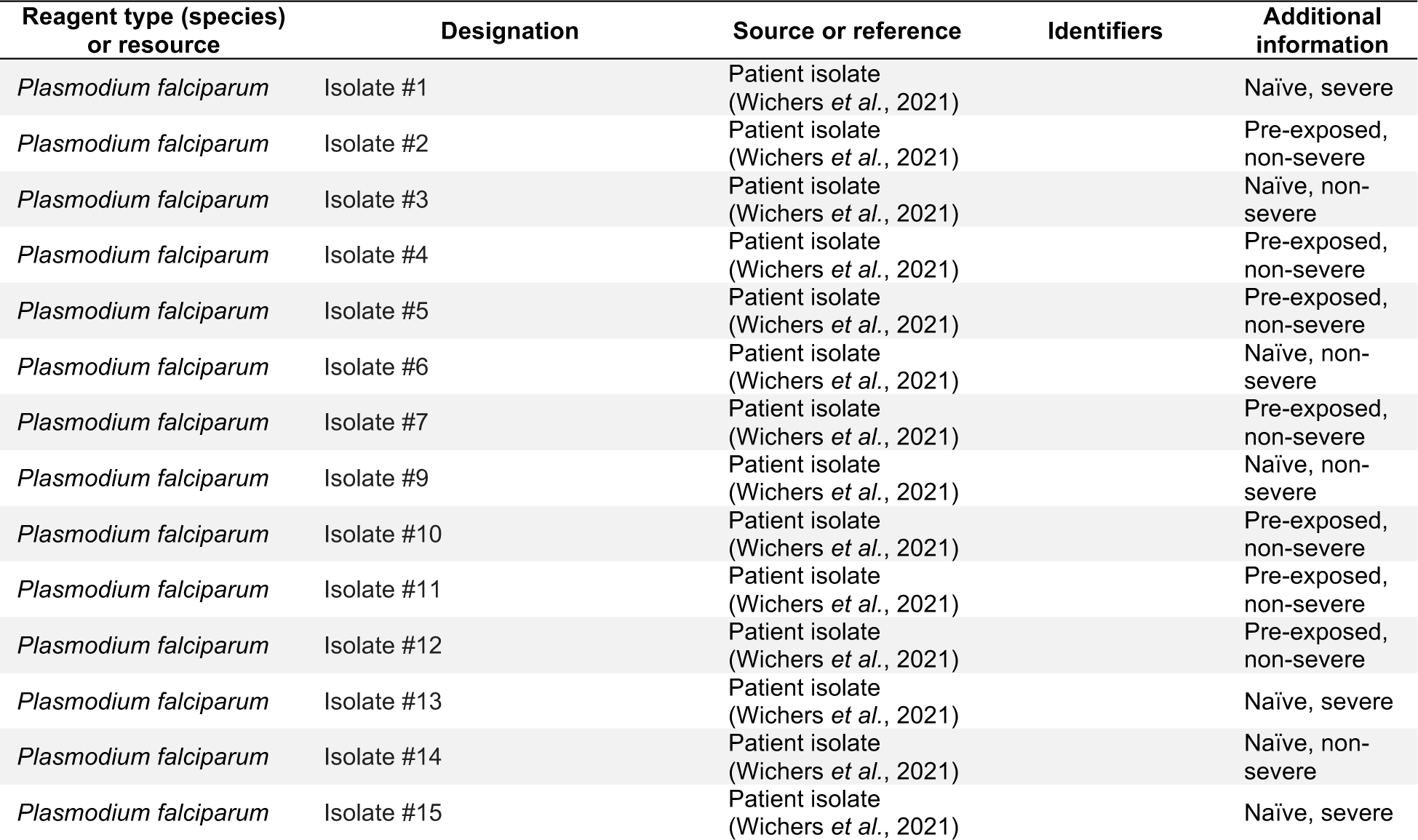
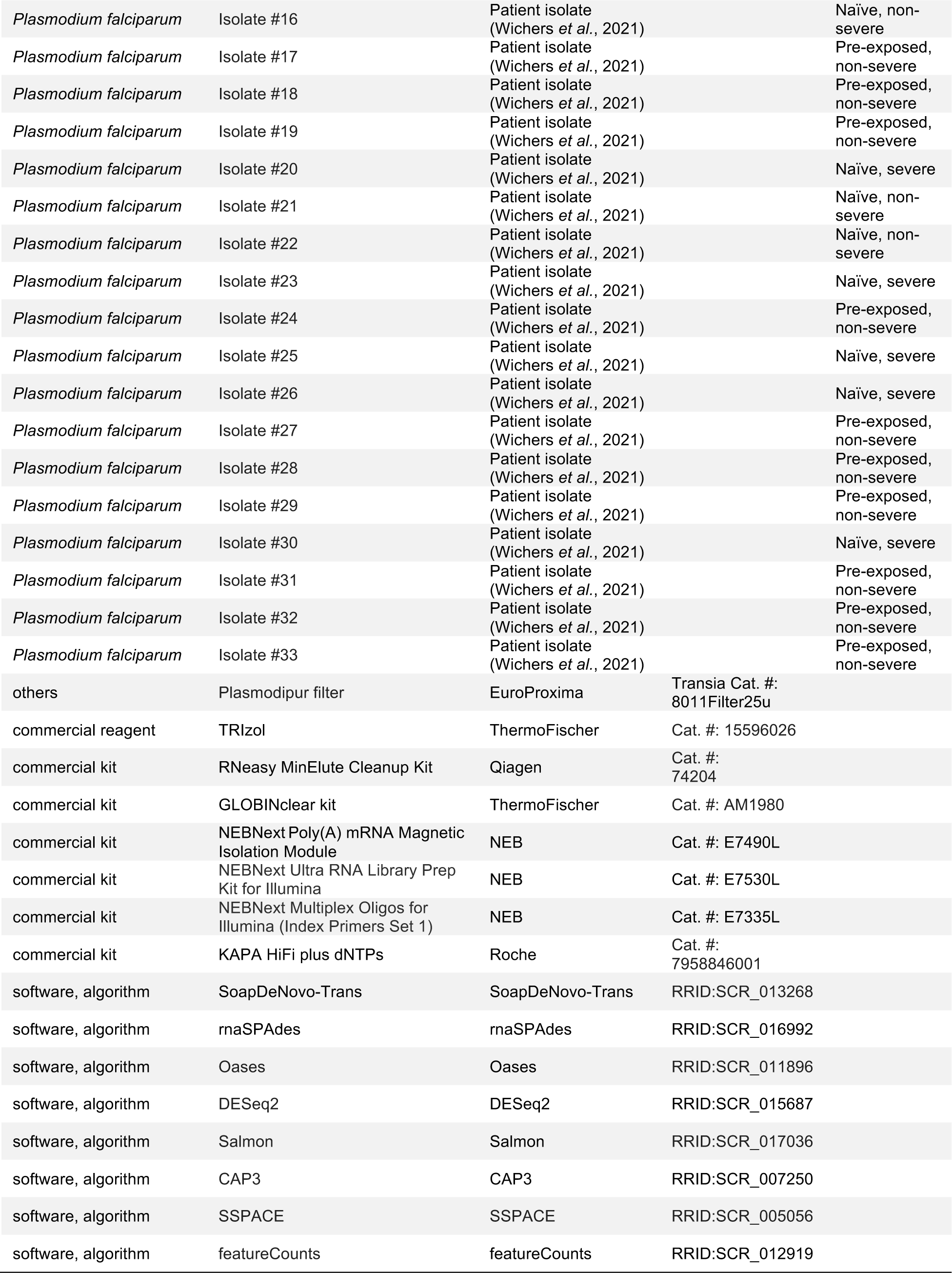
Key Resources Table.

## Materials availability statement

The authors confirm that the data supporting the findings of this study are available within the article and its supplementary materials. Additionally, the raw sequencing data are openly available in National Center for Biotechnology Information (NCBI) at BioProject ID: PRJNA679547 (https://www.ncbi.nlm.nih.gov/bioproject/?term=PRJNA679547). The already published laboratory dataset (3D7 time course RNA-sequencing data) was deposited in the European nucleotide archive (ENA): PRJEB31535 (https://www.ebi.ac.uk/ena/browser/view/PRJEB31535) (Wichers *et al*., 2019) and is also available on plasmoDB. All scripts are available in the GitHub repository (https://github.com/ClareAndradiBrown/varAssembly).

## Ethics statement

The study was conducted according to the principles of the Declaration of Helsinki, 6th edition, and the International Conference on Harmonization-Good Clinical Practice (ICH-GCP) guidelines. All 32 patients were treated as inpatients or outpatients in Hamburg, Germany (outpatient clinic of the University Medical Center Hamburg-Eppendorf (UKE) at the Bernhard Nocht Institute for Tropical Medicine, UKE, Bundeswehrkrankenhaus) (Wichers *et al*., 2021). Blood samples for this analysis were collected after patients had been informed about the aims and risks of the study and had signed an informed consent form for voluntary blood collection (n=21). In the remaining cases, no intended blood samples were collected but residuals from diagnostic blood samples were used (n=11). The study was approved by the responsible ethics committee (Ethics Committee of the Hamburg Medical Association, reference numbers PV3828 and PV4539).

## Blood sampling, processing and *in vitro* cultivation of *P. falciparum*

EDTA blood samples (1–30 mL) were collected from 32 adult falciparum malaria patients for *ex vivo* transcriptome profiling as reported by Wichers *et al*., 2021 (Wichers *et al*., 2021), hereafter termed “the original analysis”. Blood was drawn and either immediately processed (#1, #2, #3, #4, #11, #12, #14, #17, #21, #23, #28, #29, #30, #31, #32) or stored overnight at 4°C until processing (#5, #6, #7, #9, #10, #13, #15, #16, #18, #19, #20, #22, #24, #25, #26, #27, #33). If samples were stored overnight, the *ex vivo* and *in vitro* samples were still processed at the same time (so paired samples had similar storage). Erythrocytes were isolated by Ficoll gradient centrifugation, followed by filtration through Plasmodipur filters (EuroProxima) to remove residual granulocytes. At least 400 µl of the purified erythrocytes were quickly lysed in 5 volumes of pre-warmed TRIzol (ThermoFisher Scientific) and stored at -80°C until further processing (“*ex vivo* samples”). When available, the remainder was then transferred to *in vitro* culture either without the addition of allogeneic red cells or with the addition of O+ human red cells (blood bank, UKE) for dilution according to a protocol adopted from Trager and Jensen (Table S4). Cultures were maintained at 37°C in an atmosphere of 1% O2, 5% CO2, and 94% N2 using RPMI complete medium containing 10% heat-inactivated human serum (A+, Interstate Blood Bank, Inc., Memphis, USA). Cultures were sampled for RNA purification at the ring stage by microscopic observation of the individual growth of parasite isolates, and harvesting was performed at the appropriate time without prior synchronization treatment (“*in vitro* samples”). 13 of these *ex vivo* samples underwent one cycle of *in vitro* cultivation, ten of these generation 1 samples underwent a second cycle of *in vitro* cultivation. One of these generation 2 samples underwent a third cycle of *in vitro*cultivation (Table 2). In addition, an aliquot of *ex vivo* erythrocytes (approximately 50–100 µl) and aliquots of *in vitro* cell cultures collected as indicated in Supplementary file 4 were processed for gDNA purification and MSP1 genotyping as described elsewhere (Wichers *et al*., 2021, Robert *et al*., 1996).

## RNA purification, RNA-sequencing library preparation, and sequencing

RNA purification was performed as described in Wichers *et al*., 2021, using TRIzol in combination with the RNeasy MinElute Kit (Qiagen) and DNase digestion (DNase I, Qiagen). Human globin mRNA was depleted from all samples except from samples #1 and #2 using the GLOBINclear kit (ThermoFisher Scientific). The median RIN value over all *ex vivo*samples was 6.75 (IQR: 5.93–7.40), although this measurement has only limited significance for samples containing RNA of two species. Accordingly, the RIN value increased upon cultivation for all *in vitro* samples (Supplementary file 5). Customized library construction in accordance to Tonkin-Hill *et al.,* 2018, including amplification with KAPA polymerase and HiSeq 2500 125 bp paired-end sequencing was performed by BGI Genomics Co. (Hong Kong).

## Methods for assembling *var* genes

Previously Oases, Velvet, SoapDeNovo-Trans or MaSuRCA have been used for *var* transcript assembly (Wichers *et al*., 2021, Andrade *et al*., 2020, Otto *et al*., 2019, Tonkin-Hill *et al*., 2018). Previous methods either did not incorporate read error correction or focussed on gene assembly, as opposed to transcript assembly (Schulz *et al*., 2012, Zerbino & Birney, 2008, Xie *et al*., 2014, Zimin *et al*., 2013). Read error correction is important for *var* transcript assembly due to the highly repetitive nature of the *P. falciparum* genome. Recent methods have also focused on whole transcript assembly, as opposed to initial separate domain assembly followed by transcript assembly (Wichers *et al*., 2021, Andrade *et al*., 2020, Otto *et al*., 2019, Tonkin-Hill *et al*., 2018). The original analysis used SoapDeNovo-Trans to assemble the *var* transcripts, however it is currently not possible to run all steps in the original approach, due to certain tools being improved and updated. Therefore, SoapDeNovo-Trans (k=71) was used and termed the original approach.

Here, two novel methods for whole *var* transcript and *var* domain assembly were developed and their performance was evaluated in comparison to the original approach (Figure 2b). In both methods the reads were first mapped to the human g38 genome and any mapped reads were removed. Next, the unmapped reads were mapped to a modified *P. falciparum* 3D7 genome with *var* genes removed, to identify multi-mapping reads commonly present in *Plasmodium* RNA-sequencing datasets. Any mapped reads were removed. In parallel, the unmapped RNA reads from the human mapping stage were mapped against a reference of field isolate *var* exon 1 sequences and the mapped reads identified (Otto *et al*., 2019). These reads were combined with the unmapped reads from the 3D7 genome mapping stage and duplicate reads removed. All mapping was performed using sub-read align as in the original analysis (Wichers *et al*., 2021). The reads identified at the end of this process are referred to as “non-core reads”.

## Whole *var* transcript and *var* domain assembly methods

For whole *var* transcript assembly the non-core reads, for each sample separately, were assembled using rnaSPAdes (k-mer =71, read_error_correction on) (Bushmanova *et al*., 2019). Contigs were joined into larger scaffolds using SSPACE (parameters -n 31 -x 0 -k 10) (Boetzer *et al*., 2011). Transcripts < 500nt were excluded, as in the original approach. The included transcripts were annotated using hidden Markov models (HMM) (Finn *et al*., 2011) built on the *Rask et al.*, 2010 dataset and used in Tonkin-Hill *et al*., 2018. When annotating the whole transcript, the most significant alignment was taken as the best annotation for each region of the assembled transcript (e-value cut off 1e-5). Multiple annotations were allowed on the transcript if they were not overlapping, determined using cath-resolve-hits (Lewis *et al*., 2019). Scripts are available in the GitHub repository (https://github.com/ClareAndradiBrown/varAssembly)

In the *var* domain assembly approach, separate domains were assembled first and then joined up to form transcripts. First, the non-core reads were mapped (nucleotide basic local alignment tool (blastn) short read option) to the domain sequences as defined in Rask *et al*., 2010. This was found to produce similar results when compared to using tblastx. An e-value threshold of 1e-30 was used for the more conserved DBLα domains and an e-value of 1e-10 for the other domains. Next, the reads mapping to the different domains were assembled separately. rnaSPAdes (read_error_correction on, k-mer = 15), Oases (kmer = 15) and SoapDeNovo2 (kmer = 15) were all used to assemble the reads separately (Bushmanova *et al*., 2019, Xie *et al*., 2014, Schulz *et al*., 2012). The output of the different assemblers was combined into a per sample reference of domain sequences. Redundancy was removed in the reference using cd-hit (-n 8-c 0.99) (at sequence identity = 99%) (Fu *et al*., 2012). Cap3 was used to merge and extend the domain assemblies (Huang & Madan, 1999). SSPACE was used to join the domains together (parameters - n 31 -x 0 -k 10) (Boetzer *et al*., 2011). Transcript annotation was performed as in the whole transcript approach, with transcripts < 500 nt removed. Significantly annotated (1e-5) transcripts were identified and selected. The most significant annotation was selected as the best annotation for each region, with multiple annotations allowed on a single transcript if the regions were not overlapping. For both methods, a *var* transcript was selected if it contained at least one significantly annotated domain (in exon 1). *Var* transcripts that encoded only the more conserved exon 2 (acidic terminal segment (ATS) domain) were discarded.

## Validation on RNA-sequencing dataset from *P. falciparum* reference strain 3D7

Both new approaches and the original approach (SoapDeNovo-Trans, k =71) (Wichers *et al*., 2021, Tonkin-Hill *et al*., 2018) were run on a public RNA-sequencing dataset of the intra-erythrocytic life cycle stages of cultured *P. falciparum* 3D7 strain, sampled at 8-hr intervals up until 40 hrs post infection and then at 4 hr intervals up until 48 hrs post infection (ENA: PRJEB31535) (Wichers *et al*., 2019). This provided a validation of all three approaches due to the true sequence of the *var* genes being known in *P. falciparum*3D7 strain. Therefore, we compared the assembled sequences from all three approaches to the true sequence. The first best hit (significance threshold = 1e-10) was chosen for each contig. The alignment score was used to evaluate the performance of each method. The alignment score represents ⎷accuracy* recovery. The accuracy is the proportion of bases that are correct in the assembled transcript and the recovery reflects what proportion of the true transcript was assembled. Misassemblies were counted as transcripts that had a percentage identity < 99% to their best hit (i.e. the *var* transcript is not 100% contained against the reference).

## Comparison of approaches for *var* assembly on *ex vivo* samples

The *var* transcripts assembled from the 32 *ex vivo* samples using the original approach were compared to those produced from the whole transcript and domain assembly approaches. The whole transcript approach was chosen for subsequent analysis and all assembled *var* transcripts from this approach were combined into a reference, as in the original method (Wichers *et al*., 2021).

Removal of *var* transcripts with sequence id >= 99% prior to mapping was not performed in the original analysis. To overcome this, *var* transcripts were removed if they had a sequence id >= 99% against the full complement in the whole transcript approach, using cd-hit-est (Fu *et al*., 2012). Removing redundancy in the reference of assembled *var* transcripts across all samples led to the removal of 1,316 assembled contigs generated from the whole transcript approach.

This reference then represented all assembled *var* transcripts across all samples in the given analysis. The same method that was used in the original analysis was applied for quantifying the expression of the assembled *var* transcripts. The non-core reads were mapped against this reference and quantification was performed using Salmon (Patro *et al*., 2017). DESeq2 was used to perform differential expression analysis between severe versus non-severe groups and naïve versus pre-exposed groups in the original analysis (Love *et al*., 2014). Here, the same approach, as used in the original analysis, was applied to see if concordant expression estimates were obtained. As genomic sequencing was not available, this provided a confirmation of the whole transcript approach after the domain annotation step. The assembled *var* transcripts produced by the whole transcript assembly approach had their expression quantified at the transcript and domain level, as in the original method, and the results were compared to those obtained by the original method. To quantify domain expression, featureCounts was used, as in the original method with the counts for each domain aggregated (Liao *et al*., 2014). Correlation analysis between the domain’s counts from the whole transcript approach and the original method was performed for each *ex vivo* sample. Differential expression analysis was also performed using DESeq2, as in the original analysis and the results compared (Love *et al*., 2014, Wichers *et al*., 2021).

## Estimation of parasite lifecycle stage distribution in *ex vivo* and short-term *in vitro* samples

To determine the parasite life cycle stage proportions for each sample the mixture model approach of the original analysis (Tonkin-Hill *et al*., 2018, Wichers *et al*., 2021) and the SCDC approach were used (Dong *et al*., 2021, Howick *et al*., 2019). Recently, it has been determined that species-agnostic reference datasets can be used for efficient and accurate gene expression deconvolution of bulk RNA-sequencing data from any *Plasmodium* species and for correct gene expression analyses for biases caused by differences in stage composition among samples (Tebben *et al*., 2022). Therefore, the *Plasmodium berghei* single cell atlas was used as reference with restriction to 1:1 orthologs between *P. berghei* and *P. falciparum*. This reference was chosen as it contained reference transcriptomes for the gametocyte stage. To ensure consistency with the original analysis, proportions from the mixture model approach were used for all subsequent analyses (Wichers *et al*., 2021). For comparison, the proportion of different stages of the parasite life cycle in the *ex vivo* and *in vitro* samples was determined by two independent readers in Giemsa-stained thin blood smears. The same classification as the mixture model approach was used (8, 19, 30, and 42 hours post infection corresponding to ring, early trophozoite, late trophozoite and schizont stages respectively). Significant differences in ring stage proportions were tested using pairwise Wilcoxon tests. For the other stages, a modified Wilcoxon rank test for zero-inflated data was used (Wang *et al*., 2021). *Var* gene expression is highly stage dependent, so any quantitative comparison between samples needs adjustment for developmental stage. The life cycle stage proportions determined from the mixture model approach were used for adjustment.

## Characterising *var* transcripts

The whole transcript approach was applied to the paired *ex vivo* and *in vitro* samples. Significant differences in the number of assembled *var* transcripts and the length of the transcripts across the generations was tested using the paired Wilcoxon test. Redundancy was removed from the assembled *var* transcripts and transcripts and domains were quantified using the approach described above. Three additional filtering steps were applied separately to this reference of assembled *var* transcripts to ensure the *var* transcripts that went on to have their expression quantified represented true *var* transcripts. The first method restricted *var* transcripts to those greater than 1500nt containing at least 3 significantly annotated *var* domains, one of which had to be a DBLα domain. The second restricted *var* transcripts to those greater than 1500nt and containing a DBLα domain. The third approach restricted *var* transcripts to those greater than 1500nt with at least 3 significant *var* domain annotations.

## Per patient *var* transcript expression

A limitation of *var* transcript differential expression analysis is that it assumes all *var* sequences have the possibility of being expressed in all samples. However, since each parasite isolate has a different set of *var* gene sequences, this assumption is not completely valid. To account for this, *var* transcript expression analysis was performed on a per patient basis. For each patient, the paired *ex vivo* and *in vitro* samples were analysed. The assembled *var* transcripts (at least 1500nt and containing 3 significantly annotated *var* domains) across all the generations for a patient were combined into a reference, redundancy was removed as described above, and expression was quantified using Salmon (Patro *et al*., 2017). *Var* transcript expression was ranked, and the rankings compared across the generations.

## *Var* expression homogeneity (VEH)

VEH is defined as the extent to which a small number of *var* gene sequences dominate an isolate’s expression profile (Warimwe *et al*., 2013). Previously, this has been evaluated by calculating a commonly used α diversity index, the Simpson’s index of diversity. Different α diversity indexes put different weights on evenness and richness. To overcome the issue of choosing one metric, α diversity curves were calculated (Wagner *et al*., 2018). Equation 1 is the computational formula for diversity curves. *D* is calculated for *q* in the range 0 to 3 with a step increase of 0.1 and *p* in this analysis represented the proportion of *var* gene expression dedicated to *var* transcript *k*. *q* determined how much weight is given to rare vs abundant *var* transcripts. The smaller the *q* value, the less weight was given to the more abundant *var* transcript. VEH was investigated on a per patient basis.

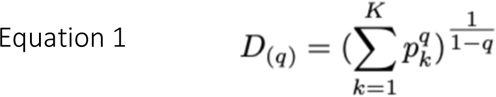

## Conserved *var* gene variants

To check for the differential expression of conserved *var* gene variants *var1-3D7*, *var1-IT* and *var2csa*, all assembled transcripts significantly annotated as such were identified. For each conserved gene, Salmon normalised read counts (adjusted for life cycle stage) were summed and expression compared across the generations using a pairwise Wilcoxon rank test.

## Differential expression of *var* domains from *ex vivo* to *in vitro* samples

Domain expression was quantified using featureCounts, as described above (Liao *et al*., 2014). DESeq2 was used to test for differential domain expression, with five expected read counts in at least three patient isolates required, with life cycle stage and patient identity used as covariates. For the *ex vivo* versus *in vitro* comparisons, only *ex vivo* samples that had paired samples in generation 1 underwent differential expression analysis, given the extreme nature of the polymorphism seen in the *var* genes.

## *Var* group expression analysis

The type of the *var* gene is determined by multiple parameters: upstream sequence (ups), chromosomal location, direction of transcription and domain composition. All regular *var* genes encode a DBLα domain in the N-terminus of the PfEMP1 protein (Figure 1c). The type of this domain correlates with previously defined *var* gene groups, with group A encoding DBLα1, groups B and C encoding DBLα0 and group B encoding a DBLα2 (chimera between DBLα0 and DBLα1) (Figure 1c). The DBLα domain sequence for each transcript was determined and for each patient a reference of all assembled DBLα domains combined. The relevant sample’s non-core reads were mapped to this using Salmon and DBLα expression quantified (Patro *et al*., 2017). DESeq2 normalisation was performed, with patient identity and life cycle stage proportions included as covariates and differences in the amounts of *var* transcripts of group A compared with groups B and C assessed (Love *et al*., 2014). A similar approach was repeated for NTS domains. NTSA domains are found encoded in group A *var* genes and NTSB domains are found encoded in group B and C *var* genes (Figure 1c).

## Quantification of total *var* gene expression

The RNA-sequencing reads were blastn (with the short-blastn option on and significance = 1e-10) against the LARSFADIG nucleotide sequences (142 unique LARSFADIG sequences) to identify reads containing the LARSFADIG motifs. This approach has been described previously (Andrade *et al*., 2020). Once the reads containing the LARSFADIG motifs had been identified, they were used to assembled the LARSFADIG motif. Trinity (Henschel, 2012) and rnaSPAdes (Bushmanova *et al*., 2019) were used separately to assemble the LARSFADIG motif, and the results compared. The sequencing reads were mapped back against the assemblies using bwa mem (Li, 2013), parameter

-k 31 -a (as in Andrade *et al*., 2020). Coverage over the LARSFADIG motif was assessed by determining the coverage over the middle of the motif (S) using Samtools depth (Danecek *et al*., 2021). These values were divided by the number of reads mapped to the *var* exon 1 database and the 3D7 genome (which had *var* genes removed) to represent the proportion of total gene expression dedicated to *var* gene expression (similar to an RPKM). The results of both approaches were compared. This method has been validated on 3D7, IT and HB3 *Plasmodium* strains. *Var2csa* does not contain the LARSFADIG motif, hence this quantitative analysis of global *var* gene expression excluded *var2csa* (which was analysed separately). Significant differences in total *var* gene expression were tested by constructing a linear model with the proportion of gene expression dedicated to *var* gene expression as the response variable, the generation and life cycle stage as an independent variables and the patient identity included as a random effect.

## *Var* expression profiling by DBLα-tag sequencing

DBLα-tag sequence analysis was performed as in the original analysis (Wichers *et al*., 2021), with Varia used to predict domain composition (Mackenzie *et al*., 2022). The proportion of transcripts encoding NTSA, NTSB, DBLα1, DBLα2 and DBLα0 domains were determined for each sample. These expression levels were used as an alternative approach to see whether there were changes in the *var* group expression levels through culture.

The consistency of domain annotations was also investigated between the DBLα-tag approach and the assembled transcripts. This was investigated on a per patient basis, with all the predicted annotations from the DBLα-tag approach for a given patient combined. These were compared to the annotations from all assembled transcripts for a given patient. DBLα annotations and DBLα-CIDR annotations were compared. This provided another validation of the whole transcript approach after the domain annotation step and was not dependent on performing differential expression analysis.

For comparison of both approaches (DBLα-tag sequencing and our new whole transcript approach), the same analysis was performed as in the original analysis (Wichers *et al*., 2021). All conserved variants (*var1, var2csa* and *var3*) were removed as they were not properly amplified by the DBLα-tag approach. To identify how many assembled transcripts, specifically the DBLα region, were found in the DBLα-tag approach, we applied BLAST. As in the original analysis, a BLAST database was created from the DBLα-tag cluster results and screened for the occurrence of those assembled DBLα regions with more than 97% seq id using the “megablast” option. This was restricted to the assembled DBLα regions that were expressed in the top 75^th^ percentile to allow for a fair comparison, as only DBLα-tag clusters with more than 10 reads were considered. Similarly, to identify how many DBLα-tag sequences were found in the assembled transcripts, a BLAST database was created from the assembled transcripts and screened for the occurrence of the DBLα-tag sequences with more than 97% seq id using the “megablast” option. This was performed for each sample.

## Core gene differential expression analysis

Subread align was used, as in the original analysis, to align the reads to the human genome and *P. falciparum* 3D7 genome, *with var, rif, stevor, surf* and *rRNA* genes removed (Liao *et al*., 2013). HTSeq count was used to quantify gene counts (Anders *et al*., 2015). DESeq2 was used to test for differentially expressed genes with five read counts in at least three samples being required (Love *et al*., 2014). Parasite life cycle stages and patient identity were included as covariates. GO and KEGG analysis was performed using ShinyGo and significant terms were defined by having a Bonferroni corrected p-value < 0.05 (Ge *et al*., 2020).

## Funding / Acknowledgements

CAB received support from the Wellcome Trust (4-Year PhD programme, grant number 220123/Z/20/Z). Infrastructure support for this research was provided by the NIHR Imperial Biomedical Research Centre and Imperial College Research Computing Service, DOI: 10.14469/hpc/2232. JSWM, YDH and AB were funded by the German Research Foundation (DFG) grants BA 5213/3-1 (project #323759012) and BA 5213/6-1 (project #433302244). TO is supported by the Wellcome Trust grant 104111/Z/14/ZR. The funders had no role in study design, data collection and analysis, decision to publish, or preparation of the manuscript. JB acknowledges support from Wellcome (100993/Z/13/Z)

## Author contribution

Conceptualization: CAB, TDO, AB, AJC

Methodology: CAB, MFD, TL, TDO

Software: CAB

Validation: CAB, AB

Formal analysis: CAB

Investigation: JSW, HvT, YDH, JAMS, HSH, EFH, AB

Resources: TL, AJC, AB

Data curation: CAB, AB

Writing – original draft: CAB, TDO, AJC, AB

Writing – review & editing: CAB, JSW, MFD, TL, TWG, JB, TDO, AJC, AB

Visualization: CAB

Supervision: TWG, JB, TDO, AJC, AB

Project Administration: CAB, AJC, AB

Funding acquisition: AJC, AB

All authors read and approved the manuscript.

---

**Figure 2 – Figure supplement 1:**
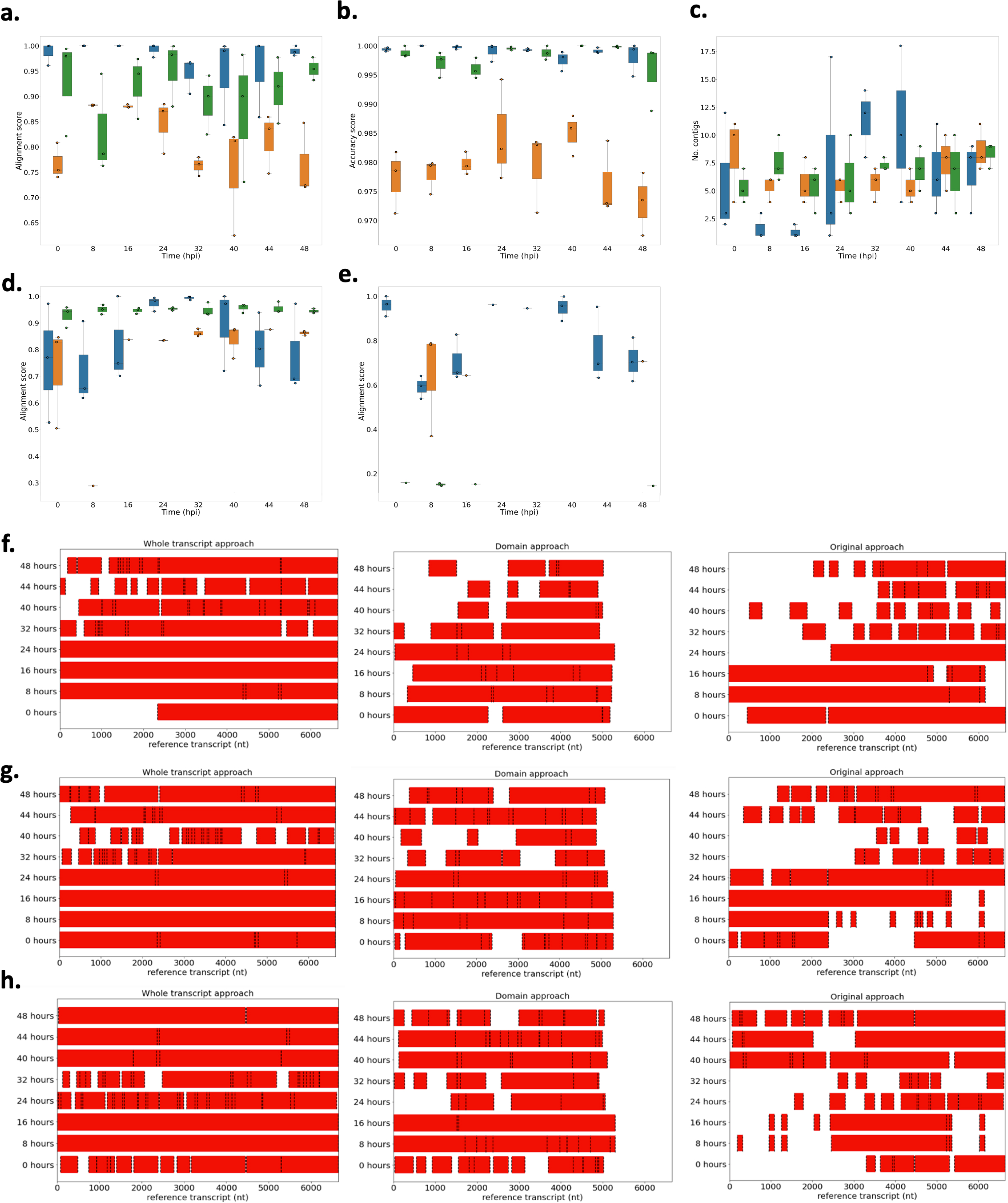
Performance of novel computational pipelines for *var* assembly on *Plasmodium falciparum* 3D7: The three approaches (whole transcript: blue, domain approach: orange, original approach: green) were applied to a public RNA-seq dataset (ENA: PRJEB31535) of the intra-erythrocytic life cycle stages of 3 biological replicates of cultured *P. falciparum* 3D7, sampled at 8-hour intervals up until 40hrs post infection (hpi) and then at 4-hour intervals up until 48 hpi (Wichers et al., 2019). Boxplots show the data from the 3 biological replicates for each time point in the intra-erythrocytic life cycle: **a)** alignment scores for the dominantly expressed *var* gene (PF3D7_0712600), **b)** accuracy scores for the dominantly expressed *var* gene (PF3D7_0712600), **c)** number of contigs required to assemble the dominant *var* gene (PF3D7_0712600), **d)** alignment scores for a middle ranking expressed *var* gene (PF3D7_0937800), **e)** alignment scores for the lowest expressed *var* gene (PF3D7_0200100). The first best blast hit (significance threshold = 1e-10) was chosen for each contig. The alignment score was used to evaluate the performance of each method. The alignment score represents ⎷accuracy* recovery. The accuracy is the proportion of bases that are correct in the assembled transcript and the recovery reflects what proportion of the true transcript was assembled. Assembly completeness of the dominant *var* gene (PF3D7_0712600, length = 6648nt) for the three approaches was assessed for each biological replicate: **f)** biological replicate 1, **g)** biological replicate 2, **h)** biological replicate 3. Dotted lines represent the start and end of the contigs required to assemble the *var* gene. Red bars represent assembled sequences relative to the dominantly expressed whole *var* gene sequence, where we know the true sequence (termed “reference transcript”).

**Figure 2 – Figure supplement 2:**
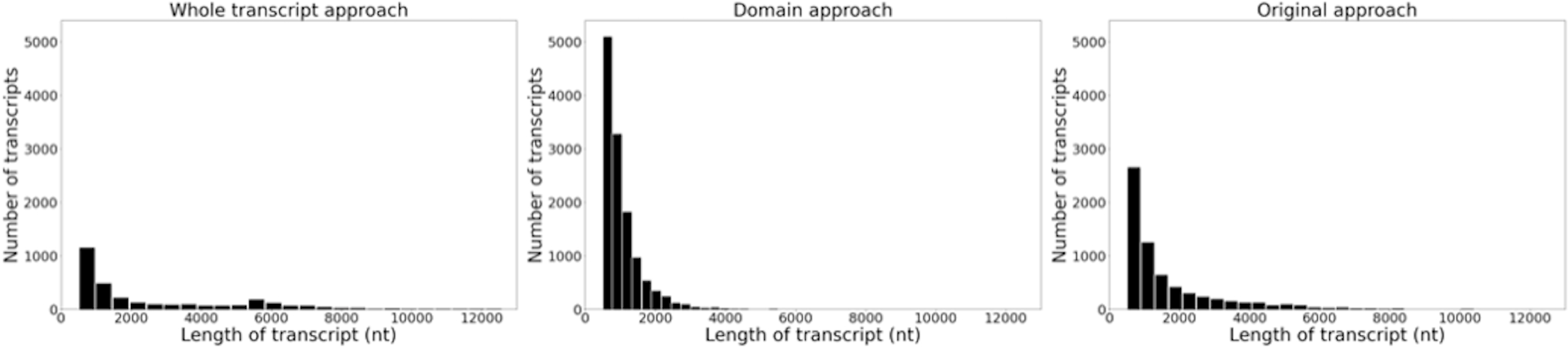
Histograms of the length and frequency of *var* transcripts produced by the different *var* assembly approaches. Approaches were applied to *ex vivo* samples (n =32) from Wichers *et al*., 2021. Assembled *var* transcripts < 500nt and not containing a significantly annotated *var* domain were removed.

**Figure 2 – Figure supplement 3:**
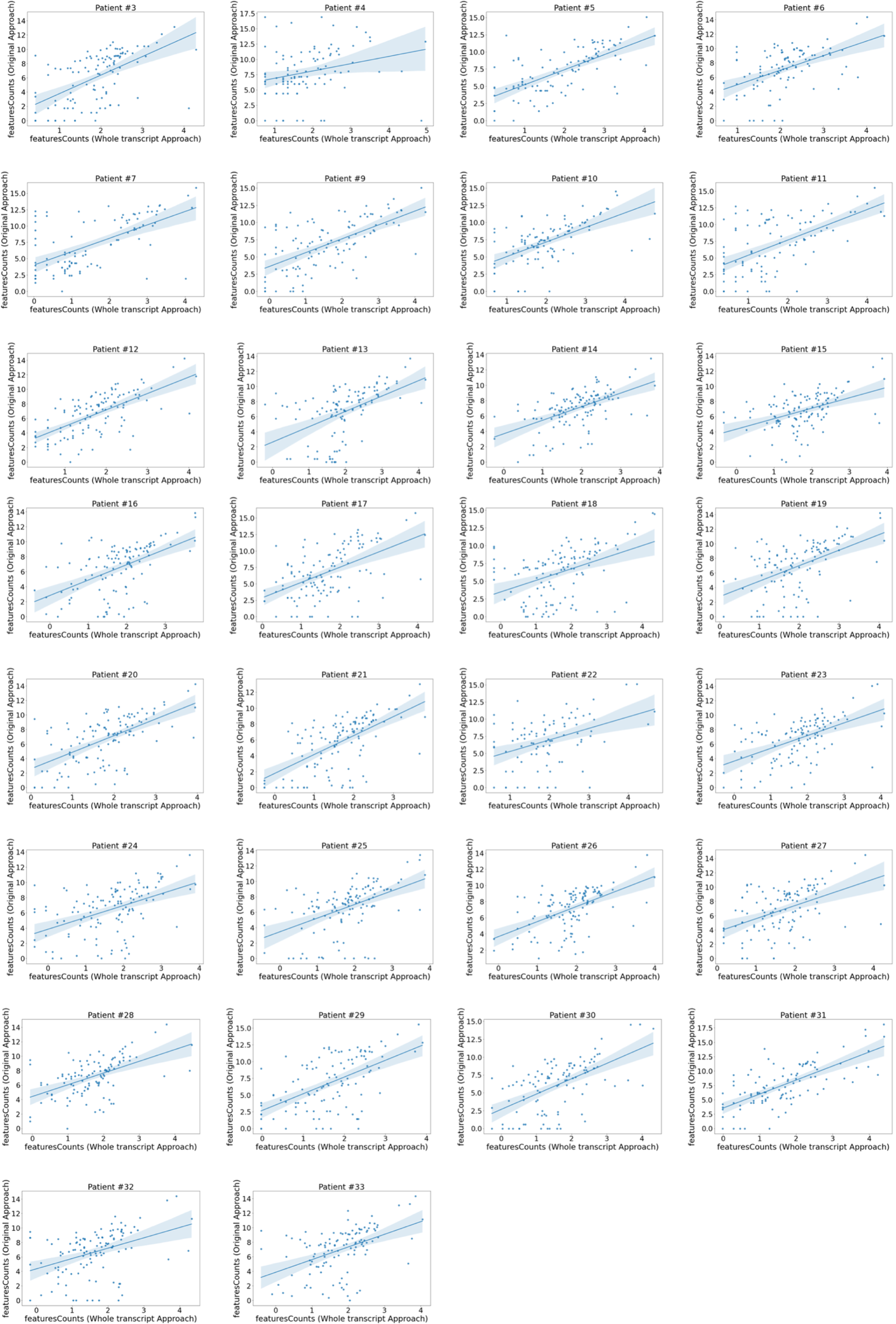
Domain expression correlation between the whole transcript approach and the original approach. Correlation plots of the original approach *var* domain counts vs those produced by the whole transcript approach. Each plot represents a patient. Domain counts were determined using featureCounts in both approaches and counts for each domain aggregated. Plots show log_2_ transformed normalized counts. The line represents a linear regression model fit.

**Figure 2 – Figure supplement 4:**
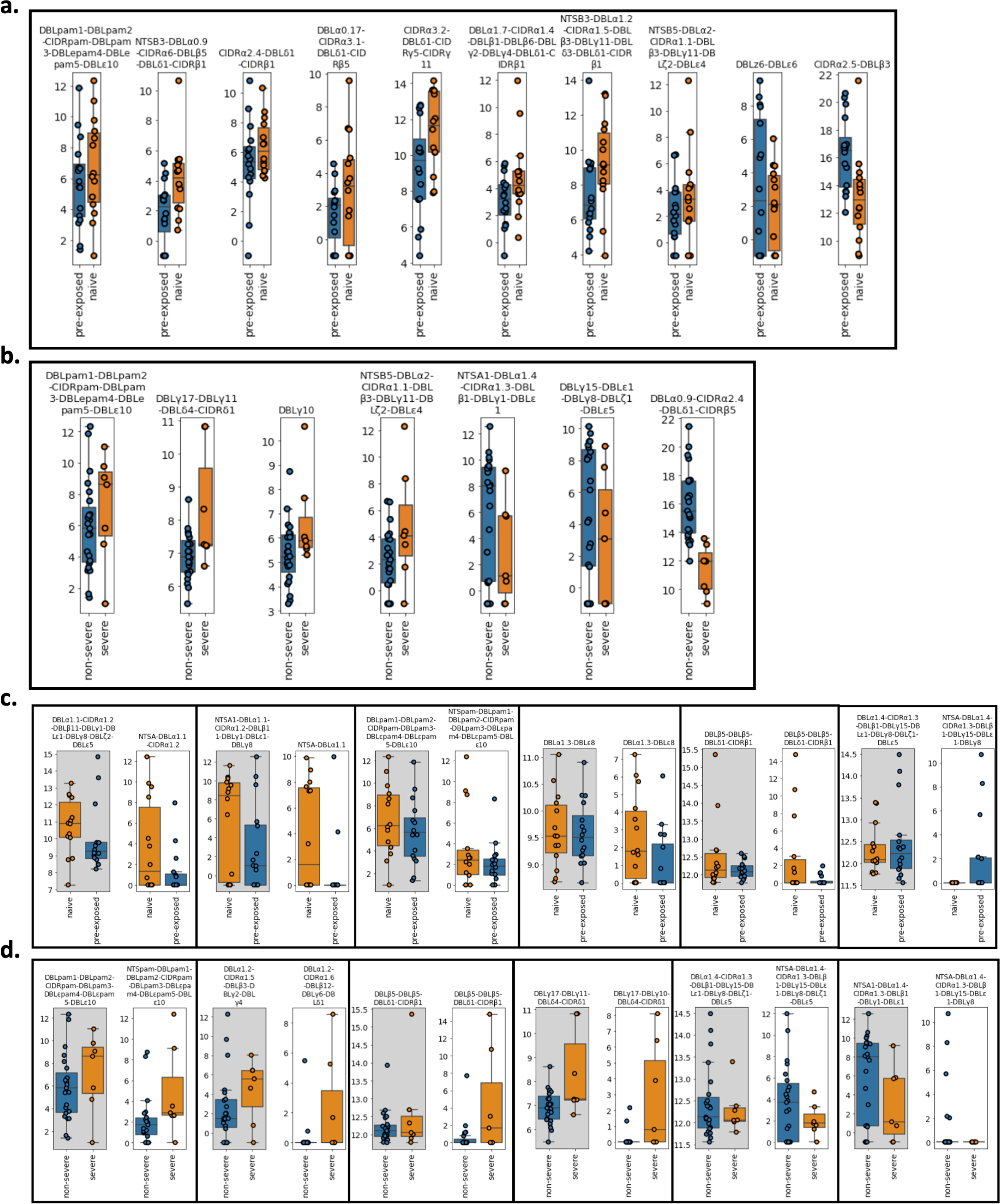
*Var* transcript expression differences derived from the whole transcript approach between a) pre-exposed (n=17) (blue) vs naïve (n=15) (orange) cases and b) severe (n=8) (orange) vs non-severe (n=24) (blue) cases. Boxplots show log_2_ normalized Salmon counts adjusted for life cycle stage (derived from the mixture model approach). Comparison of *var* transcript expression differences found in the original approach with those assembled using the whole transcript approach. These transcripts were found to be significantly differentially expressed in the **c)** pre-exposed (n=17) (blue) vs naïve (n=15) (orange) cases and **d)** severe (n=8) (orange) vs non-severe (n=24) (blue) cases in the original analysis. Each box represents a comparison of the transcript assembled using the whole transcript approach (grey background) vs using the original approach (white background). Boxplots show log_2_ Salmon normalized counts. RNA-sequencing reads of each patient sample were matched to *de novo* assembled *var* contigs with varying lengths and domains using the whole transcript approach. Shown are significantly differently expressed *var* transcripts. Differential expression analysis was performed using DESeq2.

**Figure 3 – Figure supplement 1:**
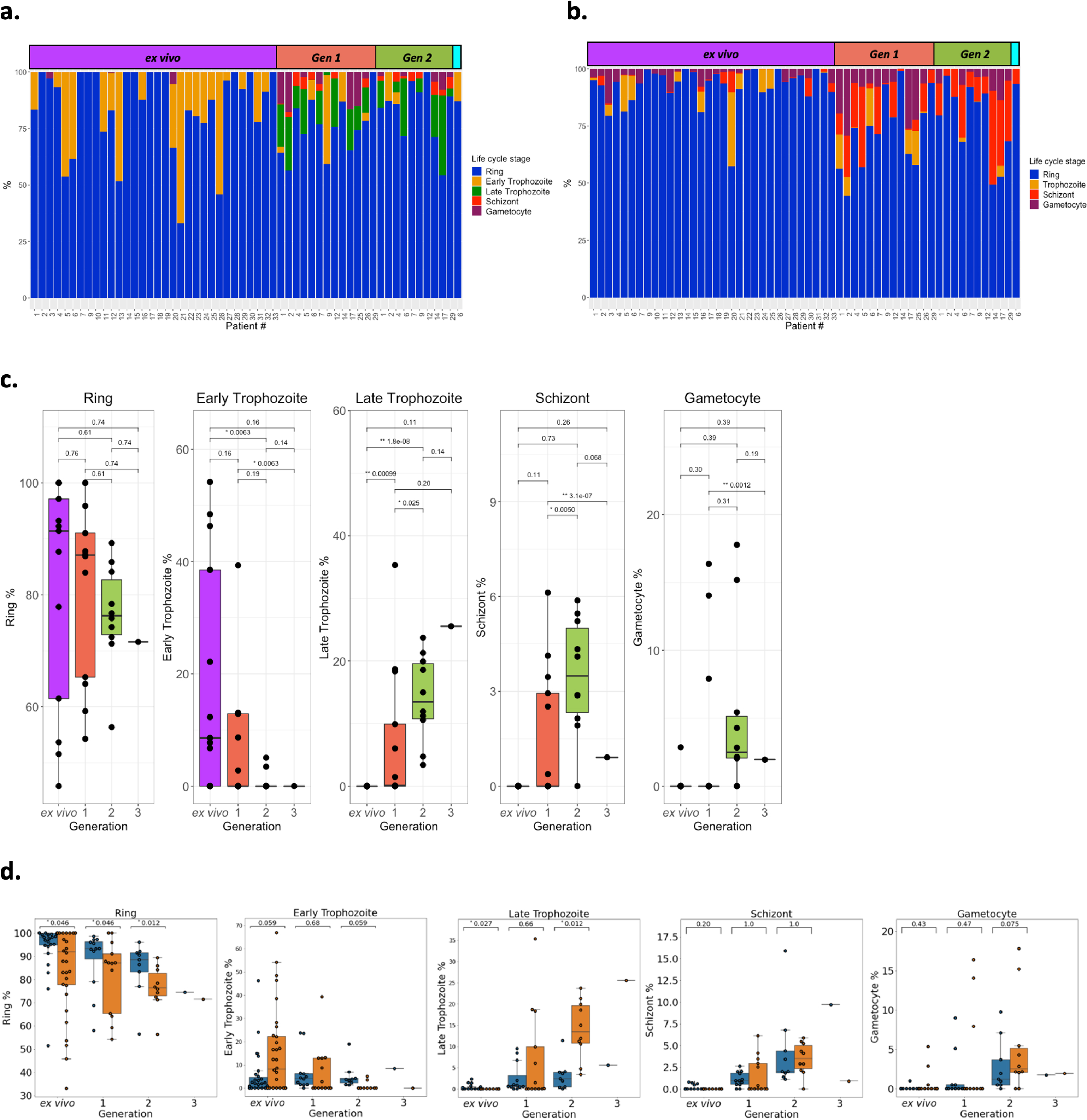
Parasite life cycle stage estimates across the generations. Life cycle stage percentages were estimated using two approaches. **a)** The first, termed the mixture model approach, was the same as used in the original analysis with 8, 19, 30, and 42 hours post infection corresponding to ring (blue), early trophozoite (orange), late trophozoite (green), schizont (red) and gametocyte (purple) stages, respectively. **b)** The second approach, termed SCDC, used SCDC, a deconvolution tool, that uses single cell reference transcriptomes of known life cycle stages, ring (blue), trophozoite (orange), schizont (red), and gametocyte (purple). The Malaria Cell Atlas for *P. berghei* was used as the reference transcriptome due to it containing gametocyte single cells. Only *P. berghei* genes that had 1:1 orthologs with *P. falciparum* 3D7 were used. Samples are ordered by generation: *ex vivo*; yellow, generation 1; pink, generation 2; blue, generation 3; green. In panels a and b life cycle stage proportions are shown for all *ex vivo* samples (n =32), paired generation 1 samples (n=13), paired generation 2 samples (n=10) and a paired generation 3 Figure shows life cycle stage proportions for all *ex vivo* samples (n =32), paired generation 1 samples (n=13), paired generation 2 samples (n=10) and a paired generation 3 sample (n=1) **c)** Significant differences in the parasite life cycle stage estimates (derived from the mixture model approach) for the paired *ex vivo* (n=13), generation 1 (n=13), generation 2 (n=10) and generation 3 (n=1) samples were tested using paired Wilcox tests (for the ring stage comparison) and a modified Wilcox on rank test for zero inflated data on the other life cycle stages. The numbers above the boxplots represent significant adjusted p-values observed (Benjamini-Hochberg adjusted). **d)** Concordance of life cycle stage estimations using the mixture model approach and counting Giemsa-stained thin blood smears. Giemsa-stained thin blood smears counting was performed by two independent researchers. Orange boxplots represent percentages determined using the mixture model approach and blue boxplots represent percentages determined using Giemsa-stained thin blood smears. Data shown represents the *ex vivo* (n=11) samples and the paired *in vitro* generations (generation 1; n=12, generation 2; n=9 and generation 3;n=1) Numbers above the boxplots represent adjusted p-values (Benjamini-Hochberg adjusted) from a pairwise Wilcoxon test. The same classification as the mixture model approach was used for the Giemsa-stained thin blood smears counting (8, 19, 30, and 42 hours post infection corresponding to ring, early trophozoite, late trophozoite and schizont stages respectively).

**Figure 3 – Figure supplement 2:**
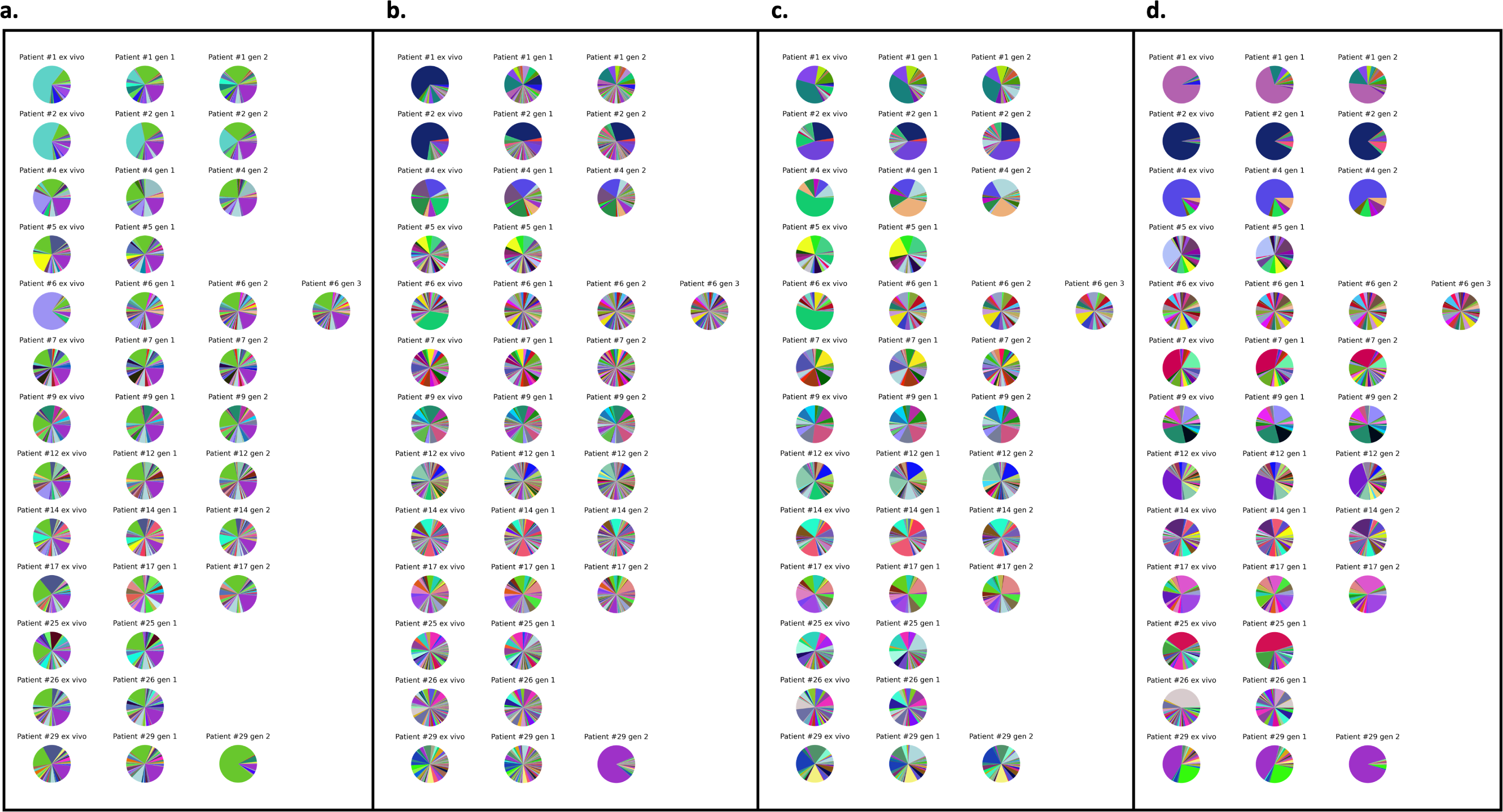
*Var* expression profiles across different mapping approaches. Different mapping approaches were used to quantify the *var* expression profiles of each sample (*ex vivo* (n=13), generation 1 (n=13), generation 2 (n=10) and generation 3 (n=1). The pooled sample approach in which all significantly assembled *var* transcripts (1500nt and containing 3 significantly annotated *var* domains) across samples were combined into a reference and redundancy was removed using cd-hit (at sequence identity = 99%) (a–c). The non-core reads of each sample were mapped to this pooled reference using **a)** Salmon, **b)** bowtie2 filtering for uniquely mapping paired reads with MAPQ >0, and **c)** bowtie2 filtering for uniquely mapping paired reads with a MAPQ > 20. **d)** The per patient approach was applied. For each patient, the paired *ex vivo* and *in vitro* samples were analysed. The assembled *var* transcripts (at least 1500nt and containing 3 significantly annotated *var* domains) across all the generations for a patient were combined into a reference, redundancy was removed using cd-hit (at sequence identity = 99%), and expression was quantified using Salmon. Pie charts show the *var* expression profile with the relative size of each slice representing the relative percentage of total *var* gene expression of each *var* transcript. Different colours represent different assembled *var* transcripts with the same colour code used across a-d.

**Figure 4 – Figure supplement 1:**
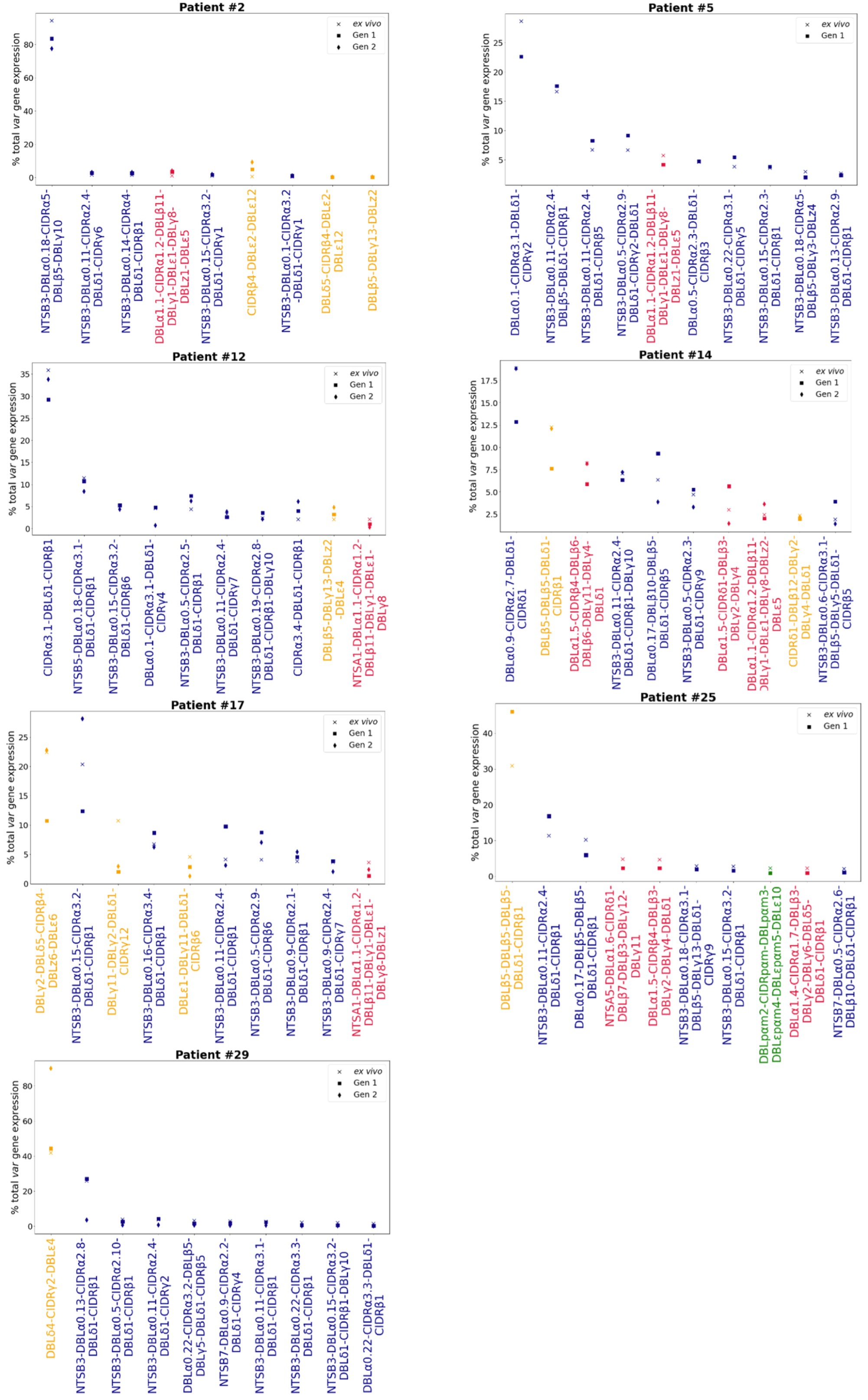
Rank *var* gene expression analysis. For each patient, the paired *ex vivo* (n=13) and *in vitro* samples (generation 1: n=13, generation 2: n=10, generation 3: n=1) were analysed. The assembled *var* transcripts with at least 1500nt and containing 3 significantly annotated *var* domains across all the generations for a patient were combined into a reference, redundancy was removed using cd-hit (at sequence identity = 99%), and expression was quantified using Salmon. *Var* transcript expression was ranked. Plots show the top 10 *var* gene expression rankings for each patient and their *ex vivo*and short-term *in vitro*cultured parasite samples. Group B/C *var* transcript (blue), group A *var* transcript (red), group E *var* transcript (green) and unknown (orange).

**Figure 4 – Figure supplement 2:**
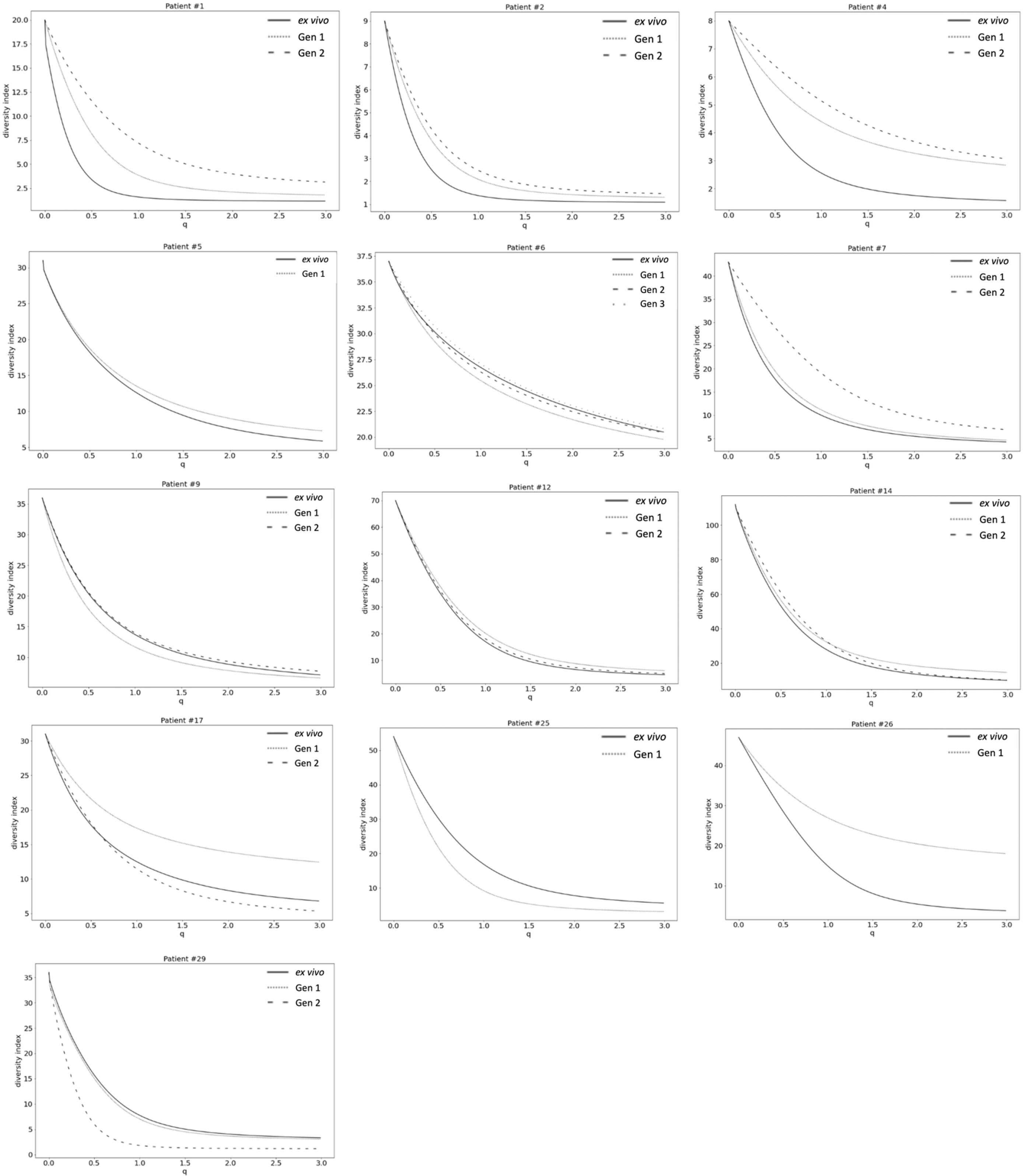
*Var* expression homogeneity (VEH). α diversity curves were determined on a per patient basis using Equation 1. *D* is calculated for *q* in the range 0 to 3 with a step increase of 0.1 and *p* in this analysis represented the proportion of *var* gene expression dedicated to *var* transcript *k*. *q* determined how much weight is given to rare vs abundant *var* transcripts. The smaller the *q* value, the less weight was given to the more abundant *var* transcript.

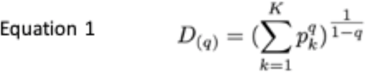

**Figure 4 – Figure supplement 3:**
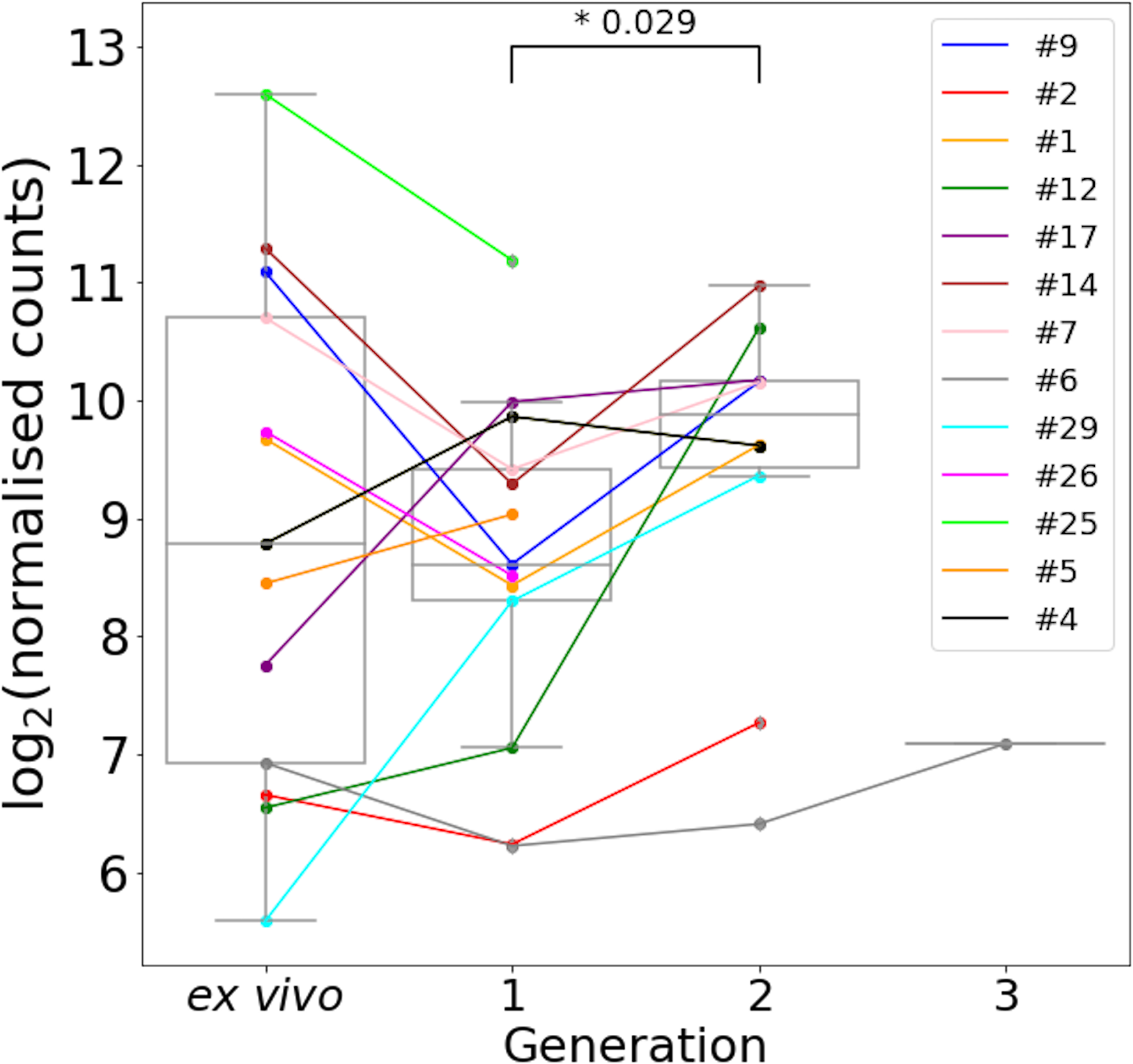
*Var2csa* expression through short-term *in vitro* cultivation. All significantly annotated *var2csa* transcripts had their log_2_ Salmon normalised count summed. Significant differences in expression levels between generations were assessed using a paired Wilcoxon test and adjusted using the Benjamini-Hochberg method. Numbers above the boxplots represent significant differences observed (FDR <=0.05). Coloured lines connect paired patient samples through the generations.

**Figure 6 – Figure supplement 1:**
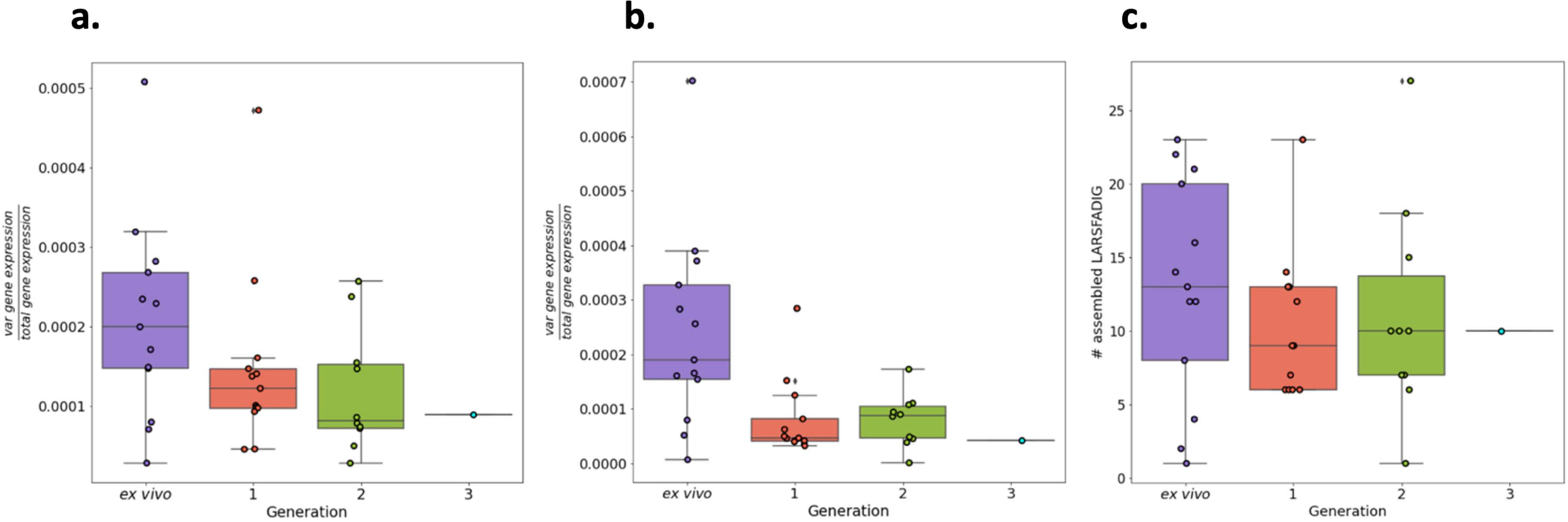
Total *var* gene expression through short-term *in vitro* culture. Boxplots show data from paired *ex vivo* (n=13), generation 1 (n=13), generation 2 (n=10) and generation 3 (n=1) parasite samples. The RNA-sequencing reads were blastn (with the short-blastn option on and significance = 1e-10) against the LARSFADIG nucleotide sequences (142 unique LARSFADIG sequences) to identify reads containing the LARSFADIG motifs. Once the reads containing the LARSFADIG motifs had been identified, they were used to assemble the LARSFADIG motif. rnaSPAdes and Trinity were used to assemble the LARSFADIG motif. The sequencing reads were mapped back against the assemblies using bwa mem, and coverage over the middle of the motif (S) determined. These values were divided by the number of reads mapped to the *var* exon 1 database and the 3D7 genome (which had *var* genes removed) to represent the proportion of total gene expression dedicated to *var* gene expression (similar to an RPKM). % total gene expression dedicated to *var* gene expression quantified using LARSFADIG motifs assembled **a)** by Trinity, **b)** by rnaSPAdes, or **c)** # assembled LARSFADIG motifs, assembled using rnaSPAdes.

**Figure 6 – Figure supplement 2:**
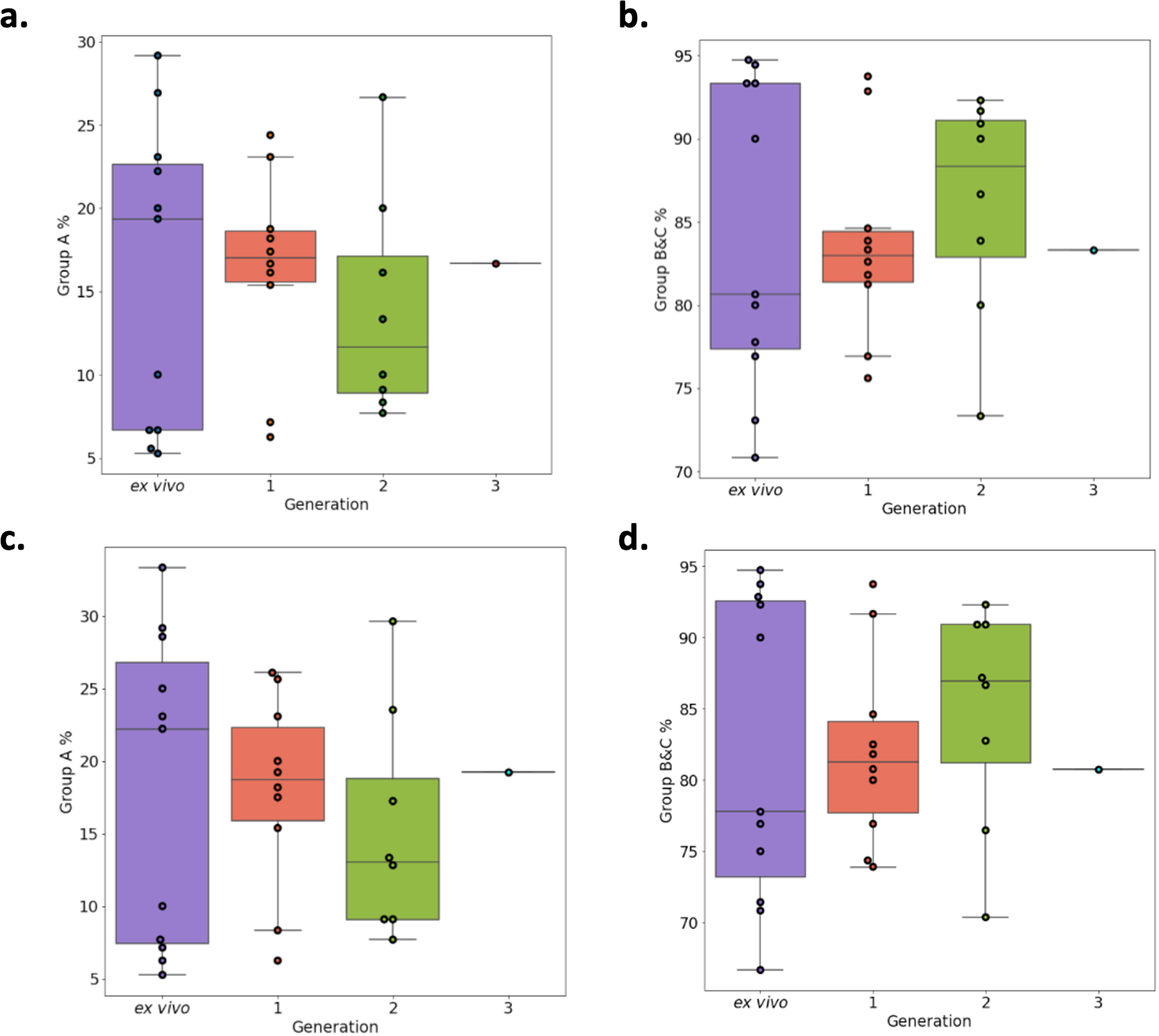
Verification of RNA-sequencing results using DBLα-tag sequencing. Amplified DBLα-tag sequences were blasted against the ∼2400 genomes on varDB to obtain subclassification into DBLα0/1/2 and prediction of adjacent head structure N-terminal segment and cysteine-rich interdomain region domains and their related binding phenotype. Proportion of each NTS and DBLα subclass were calculated, and group A and group B and C expression proportions determined. Group A *var* genes encode NTSA and DBLα1 and groups B and C encode NTSB and DBLα0. Group B *var* genes encode DBLα2. **a)** Group A transcript proportions determined by the DBLα1 domain. **b)** Group B and C transcript proportions determined by DBLα0 and DBLα2 domains. **c)** Group A transcript proportions determined by the NTSA domain. **d)** Group B and C transcript proportions determined by the NTSB domain.

**Figure 7 – Figure supplement 1:**
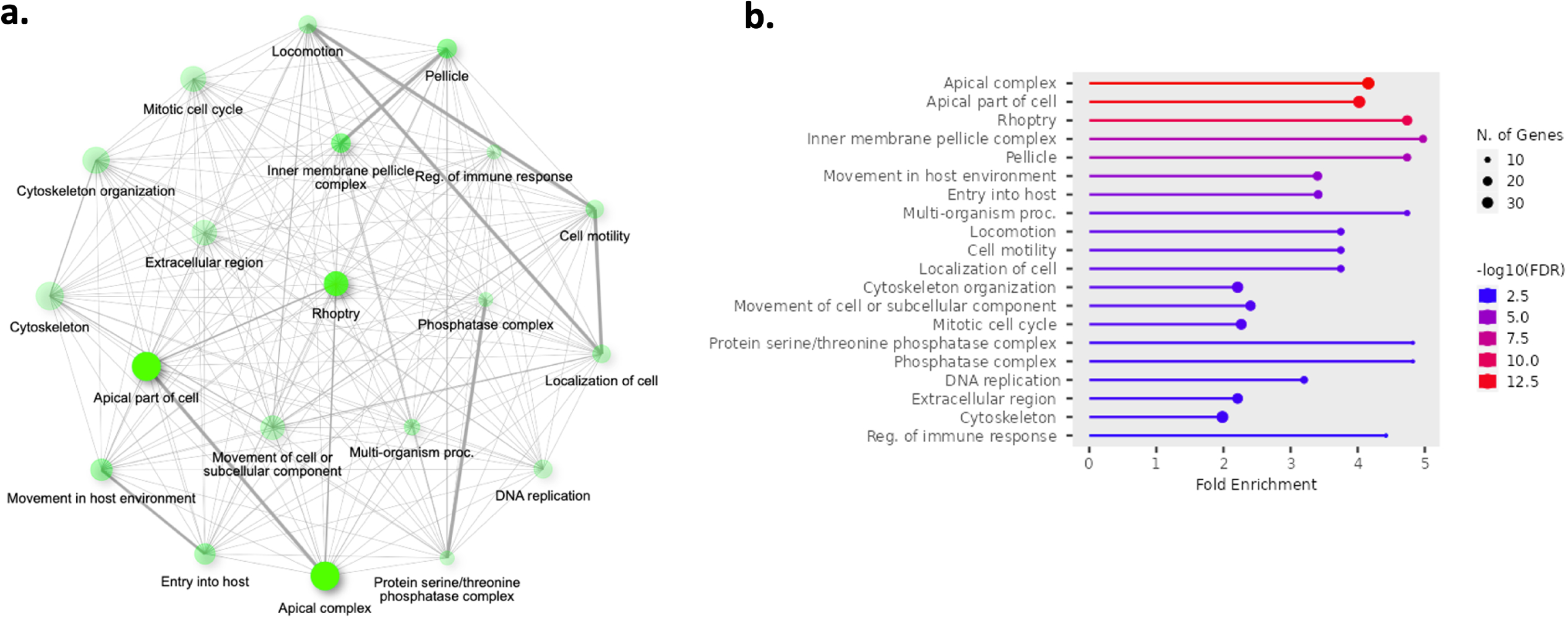
Top 20 enriched pathways in the core genes found to be significantly upregulated in generation 1 parasite samples (n=13) compared to *ex vivo* parasite samples (n=13). The 902 genes found significantly upregulated in generation 1 parasite samples underwent pathway analysis. Gene ontology and KEGG analysis was performed using ShinyGo and significant terms were defined by having a Bonferroni corrected p-value < 0.05. **a)** Network of enriched pathways. Two pathways (nodes) are connected if they share 20% or more genes. Darker nodes are more significantly enriched gene sets. Bigger nodes represent larger gene sets. Thicker edges represent more overlapped genes. **b)** Fold enrichment, significance level and the number of genes found in the pathways.

**Figure 7 – Figure supplement 2:**
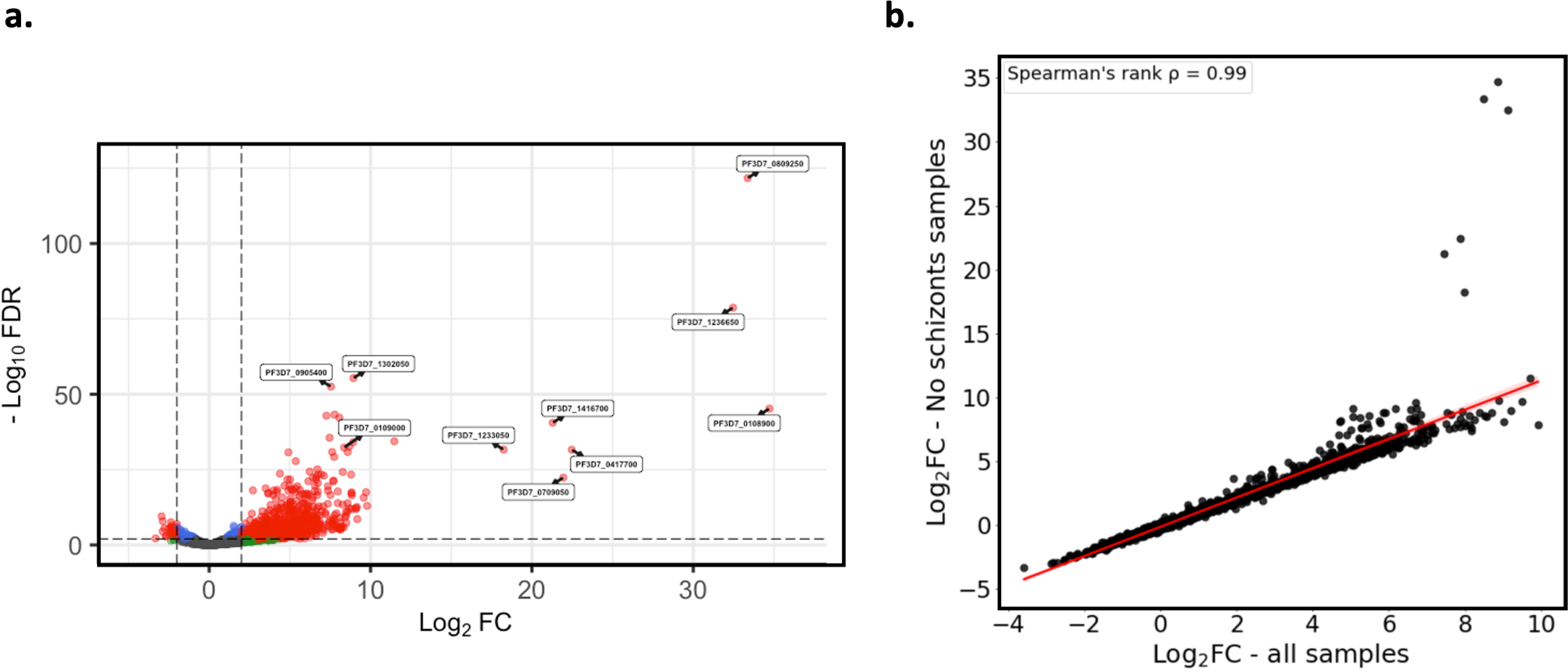
Verification of core gene expression analysis excluding schizont and gametocyte stage parasite samples. Core gene expression was assessed for paired *ex vivo*(n=6) and generation 1 (n=6) parasite samples where there were no schizont or gametocyte stage parasites, as determined using the mixture model approach. Subread align was used, as in the original analysis, to align the reads to the human genome and *P. falciparum*3D7 genome, *with var, rif, stevor, surf* and *rRNA* genes removed. HTSeq count was used to quantify gene counts. **a)** Volcano plot showing extent and significance of up-or down-regulation of core gene expression in *ex vivo* compared with paired generation 1 cultured parasites (red and blue: *P* < 0.05 after Benjamini-Hochberg adjustment for FDR, red and green: absolute log_2_ fold change log_2_FC in expression >= 2). Genes with a log_2_FC > = 2 represent those upregulated in generation 1 parasites. Genes with a log_2_FC <= -2 represent those downregulated in generation 1 parasites. **b)** Spearman’s rank correlation analysis of the log_2_FC values for the *ex vivo* vs generation 1 core gene differential expression analysis using all paired *ex vivo* (n=13) and generation 1 (n=13) samples vs *ex vivo* (n=9) and generation 1 (n=9) samples containing no schizont or gametocyte stage parasites.

**Figure 7 – Figure supplement 3:**
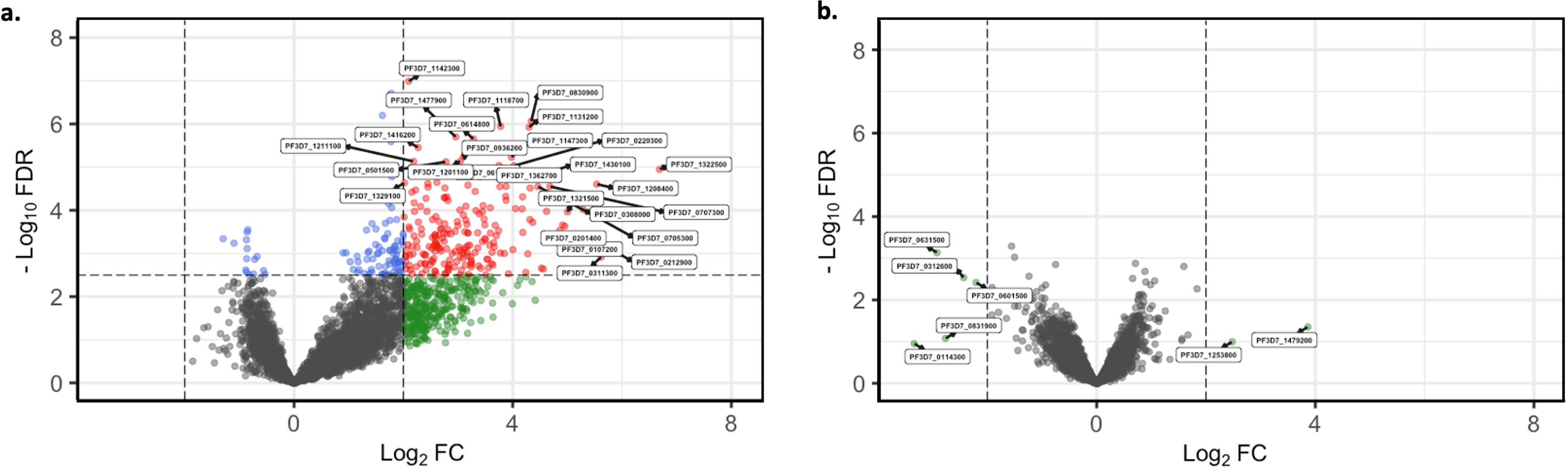
Short-term *in vitro* cultured parasites as surrogates for assessing *the in vivo* core gene transcriptome. Core gene expression was assessed for paired *ex vivo* (n=13) and generation 1 (n=13) parasite samples. Subread align was used, as in the original analysis, to align the reads to the human genome and *P. falciparum* 3D7 genome, *with var, rif, stevor, surf* and *rRNA* genes removed. HTSeq count was used to quantify gene counts (Anders et al., 2015). **a)** Volcano plot showing extent and significance of up- or down-regulation of core gene expression in malaria naïve patient *ex vivo* samples (n=6) compared with previously exposed malaria patient *ex vivo*samples (n=7) (red and blue: *P* < 0.05 after Benjamini-Hochberg adjustment for FDR; red and green: absolute log_2_ fold change log_2_FC in expression >= 2). Genes with a log_2_FC > = 2 represent those upregulated in malaria naïve patients. Genes with a log_2_FC <= -2 represent those downregulated in malaria naïve patients. **b)** Volcano plot showing extent and significance of up- or down-regulation of core gene expression in malaria naïve patient samples that underwent one cycle of *in vitro* cultivation (n=6) compared with previously exposed malaria patient samples that underwent one cycle of *in vitro* cultivation (n=7) (red and blue, *P* < 0.05 after Benjamini-Hochberg adjustment for FDR; red and green, absolute log_2_ fold change log_2_FC in expression >=2). Genes with a log_2_FC > = 2 represent those upregulated in malaria naïve patients. Genes with a log_2_FC <= -2 represent those downregulated in malaria naïve patients. Differential expression analysis was performed using DESeq2 (adjusted for life cycle stage, derived from the mixture model approach).

**Supplementary file 1: Output of the linear model to quantify total *var* gene expression during the transition of parasites into *in vitro* culture.** *Var* gene expression as a proportion of total gene expression was used as a response variable, generation and life cycle stage as independent variables and patient information as a random effect. Only paired samples of *ex vivo* parasites and generation 1 parasites were used.

**Supplementary file 2: Varia results output.** For each patient sample, the corresponding paired cluster is labelled (e.g., #6_1st_invitro_cluster_154=cluster 90ev shows cluster 154 of patient #6 generation 1 sample corresponds to cluster 90 of patient #6 *ex vivo* sample. Identical cluster sequences across paired samples were defined as having % sequence ID = > 99%.

**Supplementary file 3: Differentially expressed core genes in the *ex vivo* vs generation 1 paired analysis.** Log2FoldChange represents the log2 fold change for the *ex vivo* and generation 1 analysis. Values >0 represent genes up-regulated in generation 1 samples and values <0 represent genes down-regulated in generation 1 samples. pvalue and padj represent the p-value and adjusted pvalue (Benjamini Hochberg) respectively.

**Supplementary file 4: MSP1 genotyping results from *ex vivo* and *in vitro*-adapted *P. falciparum* isolates.**

**Supplementary file 5: Quality of the RNA samples analyzed in this study.** To characterize the overall RNA quality prior to library synthesis, the Bioanalyzer automated RNA electrophoresis system was used to visualize the samples and calculate the RIN values. The measurement of the RIN value for samples from mixed species (*H. sapiens*, *P. falciparum*) is not very meaningful, as the RIN increases the higher the proportion of a single species. This can be observed by the increase in the RIN value during *in vitro* cultivation of the parasites, as the parasite RNA content increases over time. Of the four rRNA peaks visible in particular in the *ex vivo* samples, the inner peaks represent the 18S and 28S rRNA of *P. falciparum*, the outer peaks are of human origin.

