## Supplementary material for "A novel computational pipeline for *var* gene expression augments the discovery of changes in the *Plasmodium falciparum* transcriptome during transition from *in vivo* to short-term *in vitro* culture": Table S1

**Table S1: Comparison of misassemblies produced by each *var* assembly approach.** Table shows the number (absolute) and proportion (relative to all assembled contigs) of misassemblies produced for the dominant *var* gene (PF3D7_0712600) assembly in the *P. falciparum* 3D7 dataset (ENA:PRJEB31535: A public RNA-seq dataset of the intra-erythrocytic life cycle stages of cultured *P. falciparum* 3D7 strain, sampled at 8-hour intervals up until 40 hours post infection and then at 4 hr intervals up until 48 hours post infection). A misassembly was defined as a contig whose best hit was to PF3D7_0712600 and had a sequence identity < 99% (i.e. was not 100% contained within the reference *var* transcript).

| **Sample ID** | **Time point (hrs post invasion)** | **Replicate** | **# Misassemblies** | | | | | |
| --- | --- | --- | --- | --- | --- | --- | --- | --- |
|  |  |  | **Original approach** | | **Whole transcript approach** | | **Domain approach** | |
|  |  |  | **Absolute #** | **Proportion** | **Absolute #** | **Proportion** | **Absolute #** | **Proportion** |
| ERR3196800 | 8 | 1 | 4 | 0.20 | 2 | 0.29 | 37 | 0.97 |
| ERR3196801 | 16 | 1 | 3 | 0.14 | 0 | 0.00 | 26 | 0.81 |
| ERR3196802 | 24 | 1 | 1 | 0.20 | 0 | 0.00 | 10 | 0.91 |
| ERR3196803 | 32 | 1 | 1 | 0.13 | 0 | 0.00 | 4 | 0.67 |
| ERR3196804 | 40 | 1 | 0 | 0.00 | 1 | 0.09 | 5 | 0.71 |
| ERR3196805 | 44 | 1 | 0 | 0.00 | 0 | 0.00 | 5 | 0.83 |
| ERR3196806 | 48 | 1 | 2 | 0.29 | 2 | 0.20 | 9 | 1.00 |
| ERR3196807 | 0 | 1 | 0 | 0.00 | 0 | 0.00 | 11 | 0.92 |
| ERR3196808 | 8 | 2 | 3 | 0.15 | 0 | 0.00 | 36 | 0.92 |
| ERR3196809 | 16 | 2 | 0 | 0.00 | 1 | 0.50 | 20 | 0.69 |
| ERR3196810 | 24 | 2 | 0 | 0.00 | 0 | 0.00 | 11 | 0.85 |
| ERR3196811 | 32 | 2 | 1 | 0.14 | 1 | 0.08 | 6 | 0.75 |
| ERR3196812 | 40 | 2 | 0 | 0.00 | 1 | 0.05 | 3 | 0.75 |
| ERR3196813 | 44 | 2 | 0 | 0.00 | 0 | 0.00 | 12 | 0.92 |
| ERR3196814 | 48 | 2 | 1 | 0.10 | 1 | 0.10 | 11 | 1.00 |
| ERR3196815 | 0 | 2 | 0 | 0.00 | 1 | 0.25 | 17 | 0.85 |
| ERR3196816 | 8 | 3 | 3 | 0.19 | 1 | 0.33 | 44 | 0.96 |
| ERR3196817 | 16 | 3 | 0 | 0.00 | 1 | 0.17 | 28 | 0.90 |
| ERR3196818 | 24 | 3 | 0 | 0.00 | 1 | 0.05 | 4 | 0.80 |
| ERR3196819 | 32 | 3 | 0 | 0.00 | 1 | 0.05 | 6 | 0.75 |
| ERR3196820 | 40 | 3 | 0 | 0.00 | 0 | 0.00 | 8 | 0.89 |
| ERR3196821 | 44 | 3 | 0 | 0.00 | 0 | 0.00 | 22 | 0.92 |
| ERR3196822 | 48 | 3 | 0 | 0.00 | 0 | 0.00 | 10 | 0.83 |
| ERR3196823 | 0 | 3 | 0 | 0.00 | 0 | 0.00 | 12 | 0.86 |
