## Supplementary material for "A novel computational pipeline for *var* gene expression augments the discovery of changes in the *Plasmodium falciparum* transcriptome during transition from *in vivo* to short-term *in vitro* culture": Table S2

**Table S2. Per sample *var* assembly results using the whole transcript approach.** Patient exposure and severity refer to the original patient exposure and severity that the sample originated from. # *var* transcripts before SSPACE represents the # *var* transcripts assembled before SSPACE was applied. # *var* transcripts after SSPACE represents the # *var* transcripts assembled after SSPACE was used to join contigs. # *var* significant annotated transcripts > =500nt represents the # *var* transcripts >= 500nt that contained at least one significantly annotated *var* domain. # *var* significant annotated transcripts > =1500nt and 3 domains represents the # *var* transcripts > =1500nt that contained at least three significantly annotated *var* domains. *Var* largest transcript (nt) represents the length of the longest assembled *var* transcript in that sample in nucleotides. *Var* N50 represents the length of the shortest *var* transcript where all transcripts greater than or equal to this length when summed together represent 50% of the total *var* transcript assembly length. # *var* transcripts >= 5% represents the number of *var* transcripts whose expression contributed to > 5% overall *var* gene expression. # assembled LARSFADIG represents the number of assembled LARSFADIG motifs (assembled using rnaSPAdes).

| **Sample ID** | **Patient #** | **Sex of patient** | **Generation** | **Patient exposure** | **Patient severity** | **# *var* transcripts before SSPACE** | **# *var* transcripts after SSPACE** | **# Significant *var* transcripts ≥500nt** | **# Significant *var* transcripts ≥1500nt & 3 domains** | ***Var* largest transcript (nt)** | ***Var* N50** | **# *Var* transcripts ≥5%** | **# Assembled LARSFADIG** |
| --- | --- | --- | --- | --- | --- | --- | --- | --- | --- | --- | --- | --- | --- |
| SRR13197346 | 1 | female | *ex vivo* | naive | severe | 161,641 | 161,641 | 16 | 1 | 6,383 | 2,281 | 4 | 2 |
| SRR13197345 | 2 | male | *ex vivo* | pre-exposed | non-severe | 184,373 | 184,373 | 19 | 3 | 5,665 | 5,129 | 2 | 4 |
| SRR13197323 | 4 | male | *ex vivo* | pre-exposed | non-severe | 65,961 | 65,955 | 15 | 2 | 7,726 | 5,908 | 4 | 1 |
| SRR13197320 | 5 | male | *ex vivo* | pre-exposed | non-severe | 47,979 | 47,977 | 61 | 26 | 11,337 | 5,831 | 4 | 14 |
| SRR13197319 | 6 | female | *ex vivo* | naïve | non-severe | 31,669 | 31,661 | 53 | 7 | 7,237 | 1,849 | 4 | 8 |
| SRR13197318 | 7 | male | *ex vivo* | pre-exposed | non-severe | 34,338 | 34,332 | 47 | 17 | 12,287 | 6,069 | 5 | 16 |
| SRR13197317 | 9 | female | *ex vivo* | naïve | non-severe | 68,101 | 68,100 | 83 | 31 | 9,693 | 6,169 | 3 | 20 |
| SRR13197344 | 12 | male | *ex vivo* | pre-exposed | non-severe | 38,035 | 38,026 | 99 | 30 | 7,572 | 5,138 | 4 | 12 |
| SRR13197342 | 14 | male | *ex vivo* | naïve | non-severe | 110,684 | 110,676 | 163 | 68 | 11,975 | 5,637 | 2 | 21 |
| SRR13197339 | 17 | female | *ex vivo* | pre-exposed | non-severe | 115,787 | 115,766 | 48 | 8 | 7,405 | 2,566 | 1 | 13 |
| SRR13197330 | 25 | female | *ex vivo* | naïve | severe | 43,335 | 43,326 | 84 | 46 | 11,475 | 7,011 | 2 | 22 |
| SRR13197329 | 26 | male | *ex vivo* | naïve | severe | 44,777 | 44,767 | 151 | 36 | 10,918 | 5,060 | 2 | 23 |
| SRR13197326 | 29 | male | *ex vivo* | pre-exposed | non-severe | 33,800 | 33,785 | 44 | 20 | 10,440 | 5,836 | 2 | 12 |
| SRR25659814 | 1 | female | 1 | naïve | severe | 19,902 | 19,901 | 61 | 9 | 8,648 | 3,449 | 3 | 9 |
| SRR25659802 | 2 | male | 1 | pre-exposed | non-severe | 32,014 | 32014 | 25 | 5 | 11,607 | 5,768 | 5 | 6 |
| SRR25659796 | 4 | male | 1 | pre-exposed | non-severe | 11,583 | 11,583 | 24 | 10 | 8,462 | 7,024 | 4 | 6 |
| SRR25659794 | 5 | male | 1 | pre-exposed | non-severe | 7,876 | 7,876 | 54 | 15 | 7,833 | 5,605 | 3 | 6 |
| SRR25659793 | 6 | female | 1 | naïve | non-severe | 6,765 | 6,764 | 45 | 20 | 11,345 | 6,408 | 4 | 7 |
| SRR25659812 | 7 | male | 1 | pre-exposed | non-severe | 7,316 | 7,312 | 26 | 10 | 11,369 | 5,987 | 3 | 6 |
| SRR25659810 | 9 | female | 1 | naïve | non-severe | 8,840 | 8,839 | 38 | 2 | 5,989 | 1,067 | 3 | 6 |
| SRR25659808 | 12 | male | 1 | pre-exposed | non-severe | 7,089 | 7,087 | 100 | 29 | 9,816 | 4,348 | 2 | 13 |
| SRR25659806 | 14 | male | 1 | naïve | non-severe | 11,776 | 11,775 | 138 | 38 | 10,219 | 4,622 | 2 | 23 |
| SRR25659804 | 17 | female | 1 | pre-exposed | non-severe | 7,431 | 7,430 | 67 | 25 | 8,996 | 5,301 | 2 | 14 |
| SRR25659801 | 25 | female | 1 | naïve | severe | 13,576 | 13,576 | 85 | 31 | 10,454 | 5,571 | 3 | 12 |
| SRR25659800 | 26 | male | 1 | naïve | severe | 5,833 | 5,833 | 76 | 16 | 8,452 | 4,611 | 2 | 13 |
| SRR25659799 | 29 | male | 1 | pre-exposed | non-severe | 10,424 | 10,423 | 65 | 15 | 7,101 | 4,223 | 3 | 13 |
| SRR25659813 | 1 | female | 2 | naïve | severe | 8,614 | 8,614 | 70 | 15 | 11,067 | 4,024 | 2 | 15 |
| SRR25659797 | 2 | male | 2 | pre-exposed | non-severe | 7,874 | 7,874 | 43 | 6 | 6,088 | 2,282 | 4 | 9 |
| SRR25659795 | 4 | male | 2 | pre-exposed | non-severe | 8,531 | 8,528 | 32 | 7 | 11,306 | 5,648 | 4 | 1 |
| SRR25659792 | 6 | female | 2 | naïve | non-severe | 5,628 | 5,627 | 46 | 11 | 8,547 | 5,486 | 3 | 10 |
| SRR25659811 | 7 | male | 2 | pre-exposed | non-severe | 7,260 | 7,253 | 78 | 36 | 11,412 | 5,567 | 3 | 18 |
| SRR25659809 | 9 | female | 2 | naïve | non-severe | 13,341 | 13,340 | 73 | 16 | 7,477 | 3,166 | 2 | 10 |
| SRR25659807 | 12 | male | 2 | pre-exposed | non-severe | 10,026 | 10,024 | 153 | 33 | 6,775 | 2,774 | 3 | 10 |
| SRR25659805 | 14 | male | 2 | naïve | non-severe | 9,615 | 9,612 | 159 | 54 | 9,079 | 4,472 | 3 | 27 |
| SRR25659803 | 17 | female | 2 | pre-exposed | non-severe | 9,303 | 9,301 | 55 | 12 | 8,628 | 4,087 | 4 | 7 |
| SRR25659798 | 29 | male | 2 | pre-exposed | non-severe | 16,306 | 16,304 | 80 | 20 | 7,164 | 3,468 | 4 | 7 |
| SRR25659791 | 6 | female | 3 | naïve | non-severe | 7,577 | 7,573 | 75 | 18 | 7,895 | 3,418 | 3 | 10 |
