## Supplementary material for "A novel computational pipeline for *var* gene expression augments the discovery of changes in the *Plasmodium falciparum* transcriptome during transition from *in vivo* to short-term *in vitro* culture": Table S3

**Table S3. Summary statistics of the *var* transcripts after 3 different filtering approaches were applied to the paired *ex vivo* (n =13), generation 1 (n=13), generation 2 (n=10) and generation 3 (n=1) samples.** The *var* transcripts were assembled using the whole transcript approach and all samples’ assemblies combined into a reference. The first approach filtered for *var* transcripts that contained at least 3 significantly annotated domains, one of which had to be DBLα and required the transcript to be >= 1500nt in length (3 domains, >= 1500 nt & DBLα). The second approach filtered for *var* transcripts at least 1500nt long and that contained a DBLα domain (>= 1500 nt & DBLα). The third approach filtered for transcripts that contained at least 3 significantly annotated *var* domains and were at least 1500nt in length (3 domains & >= 1500 nt). # Significantly annotated *var* transcripts represents the number of significantly annotated *var* transcripts in all samples combined. # Uniquely annotated *var* transcripts represents the number of unique *var* transcript annotations found in all samples combined. # *Var* transcripts (> =5 in at least 3 samples) represents the number of *var* transcripts after filtering for a Salmon estimated count of 5 in at least 3 samples (filtering threshold used prior to differential expression analysis). Max length of *var* transcript (nt) represents the longest transcript assembled in all samples combined. N50 represents the length of the shortest *var* transcript where all transcripts greater than or equal to this length when summed together represent 50% of the total *var* transcript assembly length. Transcripts were annotated using HMM models built on the Rask et al., 2010 dataset. When annotating the whole transcript, the most significant alignment was taken as the best annotation for each region of the assembled transcript (e-value cut off 1e-5). Multiple annotations were allowed on the transcript if they were not overlapping, determined using cath-resolve-hits.

|  | **3 domains, ≥1500 nt & DBLα** | **≥1500 nt & DBLα** | **3 domains, ≥1500 nt** |
| --- | --- | --- | --- |
| **# Significantly annotated *var* transcripts** | 543 | 568 | 746 |
| **# Unique significantly annotated *var* transcripts** | 313 | 330 | 459 |
| **# *Var* transcripts (≥5 counts in at least 3 samples)** | 309 | 161 | 455 |
| **Maximum length of *var* transcripts (nt)** | 12,287 | 12,287 | 12,287 |
| **N50** | 6,005 | 5,983 | 5,858 |
