## Supplementary material for "A novel computational pipeline for *var* gene expression augments the discovery of changes in the *Plasmodium falciparum* transcriptome during transition from *in vivo* to short-term *in vitro* culture": Table S4

**Table S4: *In vitro* culture conditions.** *In vitro* culture either had the addition of O+ human red cells (necessary due to high parasitemia or low sample volume obtained from patient) or were without allogenic red cells. Patient # represents the original *ex vivo* patient number that the sample derived from.

| **Patient #** | **Generation** | **Sample ID** | ***In vitro* condition** |
| --- | --- | --- | --- |
| 1 | 1 | SRR25659814 | Without allogeneic red cells |
|  | 2 | SRR25659813 | Without allogeneic red cells |
| 2 | 1 | SRR25659802 | Without allogeneic red cells |
|  | 2 | SRR25659797 | Without allogeneic red cells |
| 4 | 1 | SRR25659796 | Without allogeneic red cells |
|  | 2 | SRR25659795 | With O+ human red cells |
| 5 | 1 | SRR25659794 | Without allogeneic red cells |
| 6 | 1 | SRR25659793 | Without allogeneic red cells |
|  | 2 | SRR25659792 | Without allogeneic red cells |
|  | 3 | SRR25659791 | Without allogeneic red cells |
| 7 | 1 | SRR25659812 | With O+ human red cells |
|  | 2 | SRR25659811 | With O+ human red cells |
| 9 | 1 | SRR25659810 | With O+ human red cells |
|  | 2 | SRR25659809 | With O+ human red cells |
| 12 | 1 | SRR25659808 | Without allogeneic red cells |
|  | 2 | SRR25659807 | Without allogeneic red cells |
| 14 | 1 | SRR25659806 | Without allogeneic red cells |
|  | 2 | SRR25659805 | With O+ human red cells |
| 17 | 1 | SRR25659804 | Without allogeneic red cells |
|  | 2 | SRR25659803 | With O+ human red cells |
| 25 | 1 | SRR25659801 | With O+ human red cells |
| 26 | 1 | SRR25659800 | Without allogeneic red cells |
| 29 | 1 | SRR25659799 | With O+ human red cells |
|  | 2 | SRR25659798 | With O+ human red cells |
