## Supplementary figures and images for "A novel computational pipeline for *var* gene expression augments the discovery of changes in the *Plasmodium falciparum* transcriptome during transition from *in vivo* to short-term *in vitro* culture"

### Supplementary File 5

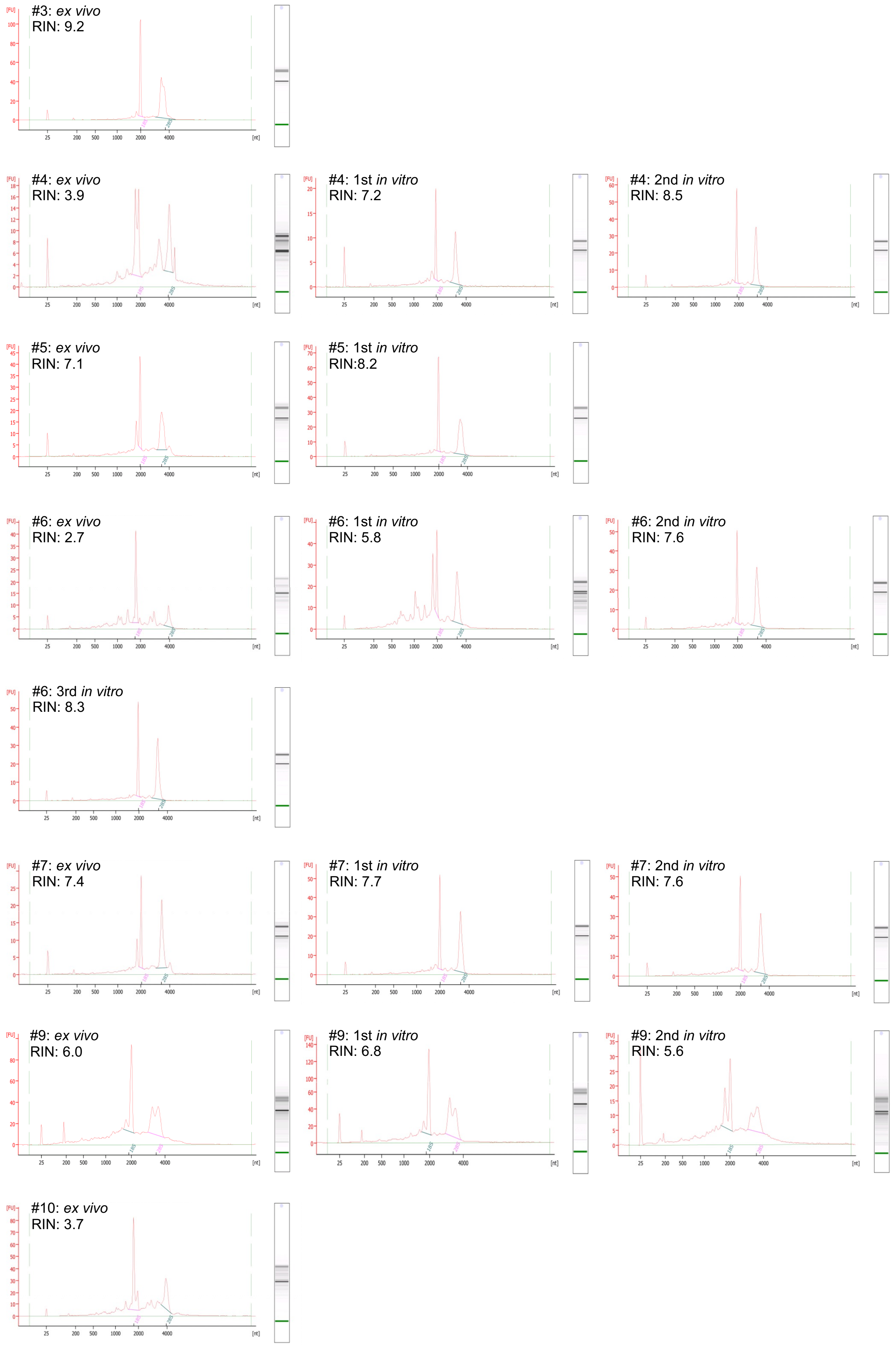

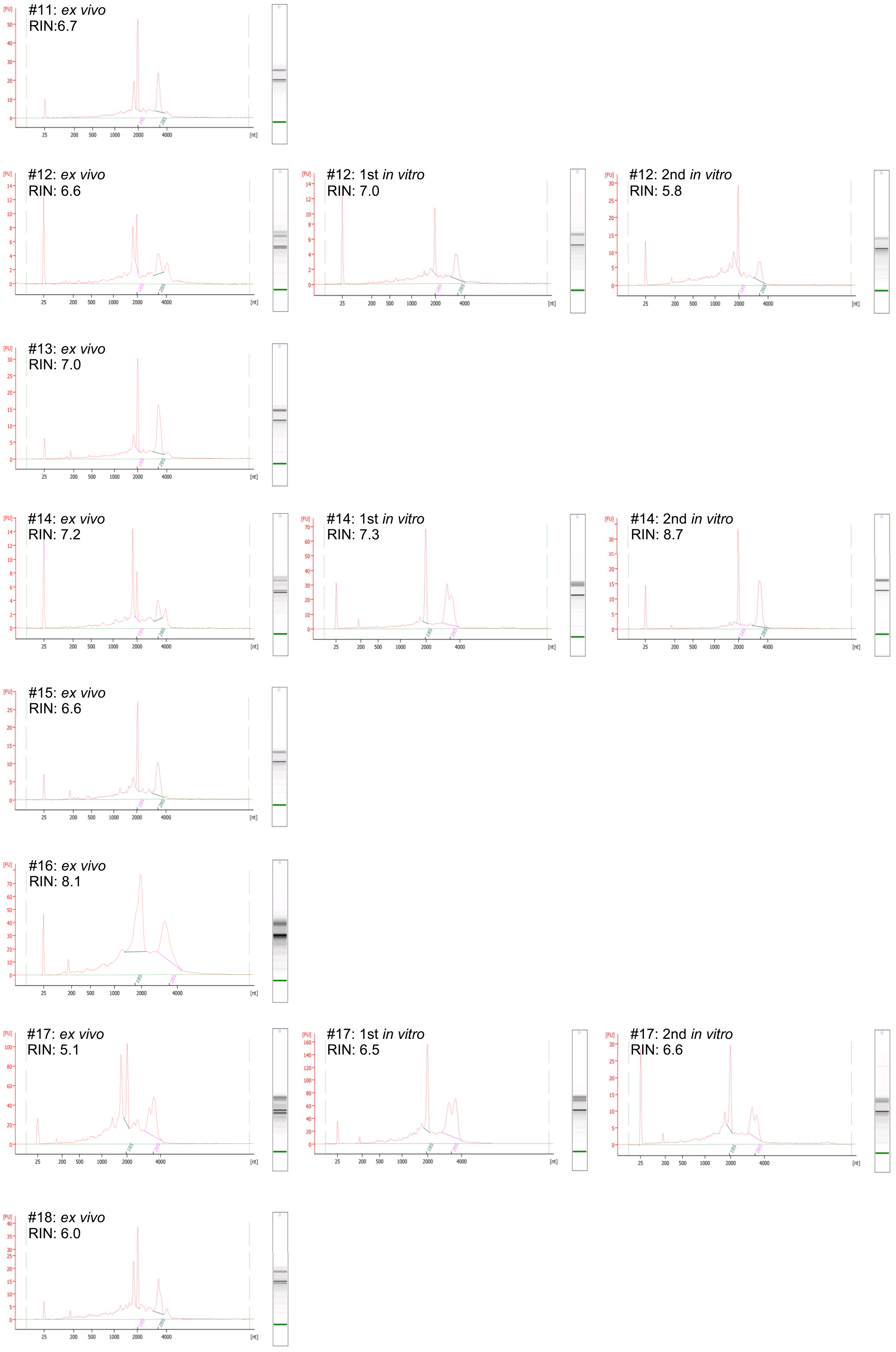

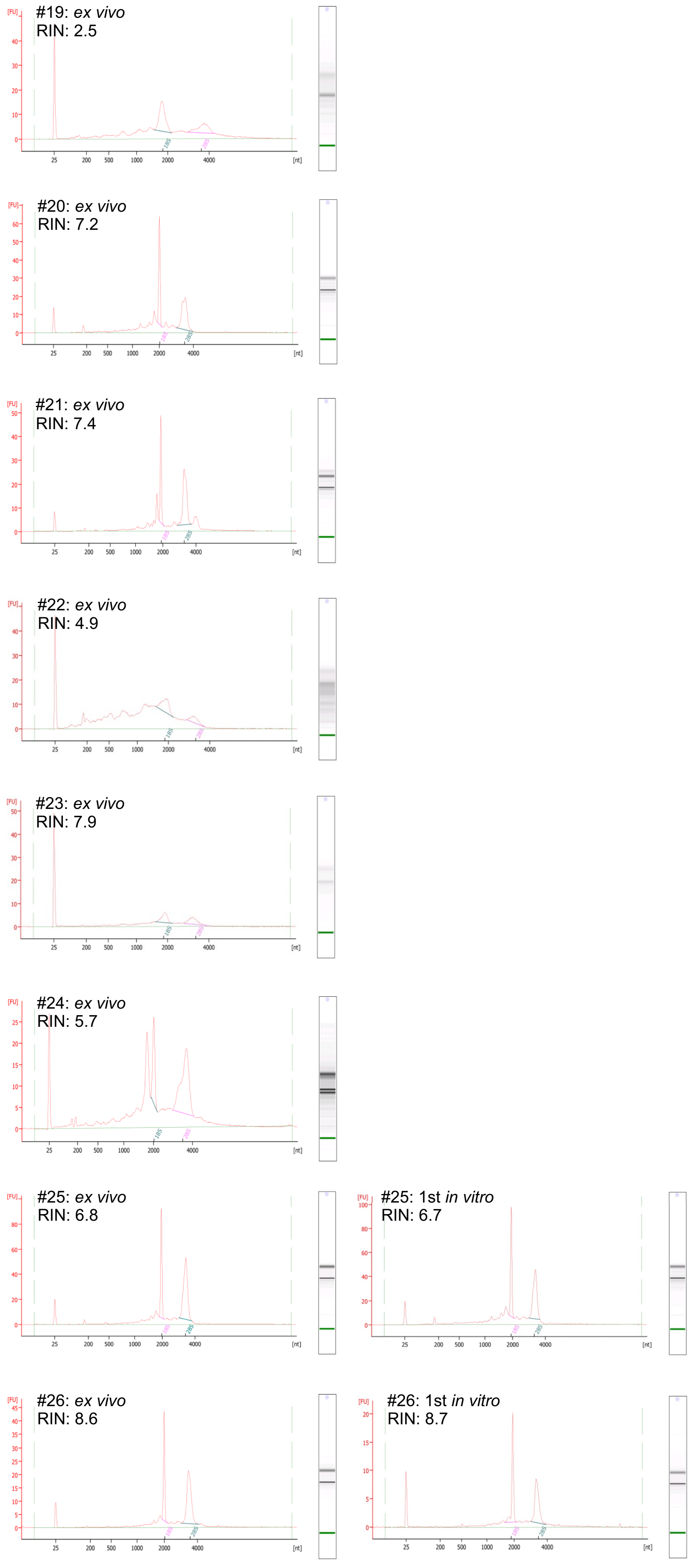

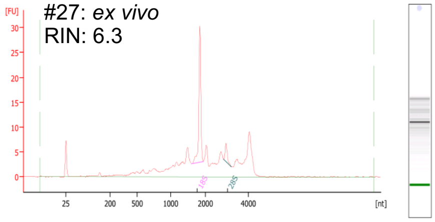

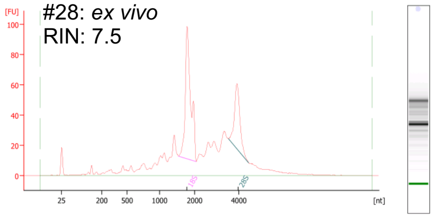

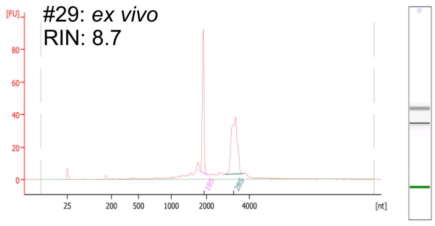

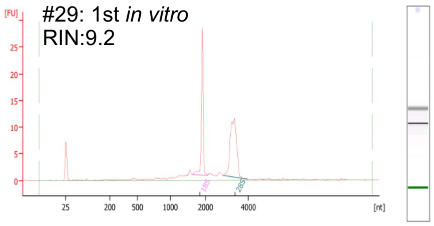

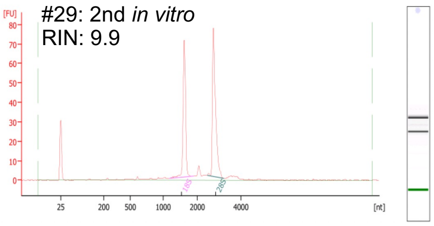

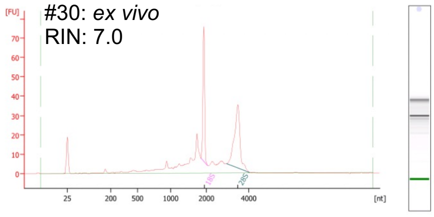

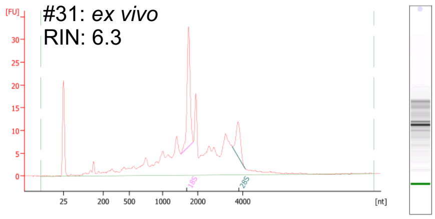

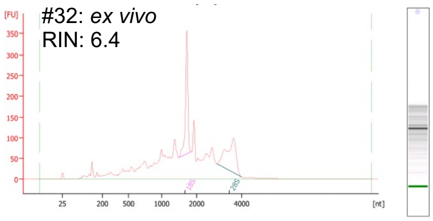

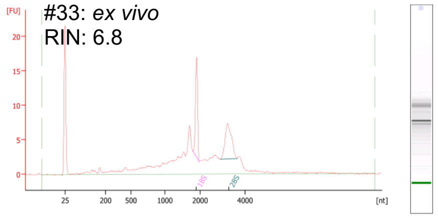
